# Thermal endurance by a hot-spring-dwelling phylogenetic relative of the mesophilic *Paracoccus*

**DOI:** 10.1101/2022.05.08.491110

**Authors:** Nibendu Mondal, Chayan Roy, Sumit Chatterjee, Jagannath Sarkar, Subhajit Dutta, Sabyasachi Bhattacharya, Ranadhir Chakraborty, Wriddhiman Ghosh

## Abstract

High temperature growth/survival was revealed in a phylogenetic relative (strain SMMA_5) of the mesophilic *Paracoccus* isolated from the 78-85°C water of a Trans-Himalayan sulfur-borax spring. After 12 h at 50°C, or 45 minutes at 70°C, in mineral salts thiosulfate (MST) medium, SMMA_5 retained ∼2% colony-forming units (CFUs), whereas comparator *Paracoccus* had 1.5% and 0% CFU left at 50°C and 70°C respectively. After 12 h at 50°C, the thermally-conditioned sibling SMMA_5_TC exhibited ∼1.5 time increase in CFU-count; after 45 minutes at 70°C, SMMA_5_TC had 7% of the initial CFU-count intact. 1000-times diluted Reasoner’s 2A medium, and MST supplemented with lithium, boron or glycine-betaine (solutes typical of the SMMA_5 habitat), supported higher CFU-retention/CFU-growth than MST. With or without lithium/boron/glycine-betaine in MST, a higher percentage of cells always remained viable (cytometry data), compared with what percentage remained capable of forming single colonies (CFU data). SMMA_5, compared with other *Paracoccus*, contained 335 unique genes, mostly for DNA replication/recombination/repair, transcription, secondary metabolites biosynthesis/transport/catabolism, and inorganic ion transport/metabolism. It’s also exclusively enriched in cell wall/membrane/envelope biogenesis, and amino acid metabolism, genes. SMMA_5 and SMMA_5_TC mutually possessed 43 nucleotide polymorphisms, of which 18 were in protein-coding genes with 13 nonsynonymous and seven radical amino acid replacements. Such biochemical and biophysical mechanisms could be involved in thermal stress mitigation which streamline the cells’ energy and resources towards system-maintenance and macromolecule-stabilization, thereby relinquishing cell-division for cell-viability. Thermal conditioning apparently helped memorize those potential metabolic states which are crucial for cell-system maintenance, while environmental solutes augmented the indigenous stability-conferring mechanisms.

**IMPORTANCE:** For a holistic understanding of microbial life’s high-temperature adaptation it is imperative to explore the biology of the phylogenetic relatives of mesophilic bacteria which get stochastically introduced to geographically and geologically diverse hot spring systems by local geodynamic forces. Here, *in vitro* endurance of high heat up to the extent of growth under special (habitat-inspired) conditions was discovered in a hot-spring-dwelling phylogenetic relative of the mesophilic *Paracoccus* species. Thermal conditioning, extreme oligotrophy, metabolic deceleration, presence of certain habitat-specific inorganic/organic solutes, and typical genomic specializations were found to be the major enablers of this conditional (acquired) thermophilicity. Feasibility of such phenomena across the taxonomic spectrum can well be paradigm-changing for the established scopes of microbial adaptation to the physicochemical extremes. Applications of conditional thermophilicity in microbial process biotechnology may be far reaching and multi-faceted.

## INTRODUCTION

Association of the typically thermophilic or hyperthermophilic bacteria and archaea with terrestrial hydrothermal habitats is axiomatic. Accordingly, our knowledge on microbial adaptation to high temperature is based largely on hot spring isolates that grow *in vitro* either obligately at ≥80°C (1–2) or facultatively between 30°C and 80°C (3–4). Members of mesophilic microbial groups (taxa having no member reported for laboratory growth at >45°C), on the other hand, though unexpected in high-temperature environments, often get stochastically introduced by local geodynamic forces to the hot spring systems (5–7), where they are detected mostly via sequencing and analysis of metagenomes (6–15), and sometimes as pure culture isolates (12, 16, 17). In such a scenario, for a holistic understanding of microbial life’s high-temperature adaptation, it becomes imperative to explore the biology of the phylogenetic relatives of mesophilic bacteria which happen to be there in geographically and geologically distinct hot spring habitats. That said, most studies on thermotolerance by mesophilic bacteria have thus far been centered on economically or clinically important strains isolated from environments other than hydrothermal ecosystems (18–22). Here we investigate high-temperature growth/survival in relation to the physiology, cell biology, and genomics of a novel, facultatively chemolithoautotrophic strain of the alphaproteobacterial genus *Paracoccus* (named SMMA_5) that was isolated (12) from the vent-water of a Trans-Himalayan sulfur-borax spring called Lotus Pond, located in the Puga geothermal area of eastern Ladakh (India), at an altitude of 4436 m, where water boils at ∼85°C (23). Remarkably, whereas no member of *Paracoccus* grows at >45°C *in vitro*, the temperature of the habitat of SMMA_5 ranges between 78°C and 85°C (7, 12, 23).

In order to understand how SMMA_5 responds to thermal stress, increase/decrease in the number of colony forming units (CFUs) present in the culture was tested at different temperatures (37°C-70°C); frequency of viable cells within the culture was determined via fluorescein diacetate (FDA) staining followed by flow cytometry. Potential effects of thermal conditioning on the isolate’s ability to grow or survive at high temperatures were tested by subjecting the organism to different iterations of “heat exposure and withdrawal” cycles. Previous geomicrobiological explorations of Lotus Pond had shown that the snow-melts and denuded soil/sediments, which infiltrate the shallow geothermal reservoir of Puga via local tectonic faults, introduce mesophilic bacteria into the hot spring system (7). Since the soil/sediment systems of the frigid deserts of Ladakh are typically poor in organic carbon (24), and because the vent-water of Lotus Pond has low dissolved solutes concentration (7), the plausible role of oligotrophy in thermal endurance by SMMA_5 was tested via high-temperature incubation in different dilution grades of Reasoner’s 2A (R2A) medium, typically used for growing oligotrophic bacteria (25). Dissolved solutes such as ions of boron and lithium, which are typical of the Puga hot springs including Lotus Pond (7, 14, 23), have been hypothesized previously as the *in situ* mitigators of the biomacromolecule-disordering effects of heat, and therefore the facilitators of the colonization of high-temperature sites by mesophilic bacteria (14). Accordingly, sodium tetraborate and lithium hydroxide were tested for their potential effects on the high-temperature growth/survival of SMMA_5. Furthermore, previous culture-independent community analyses of Lotus Pond’s vent-water (7, 12) had revealed the presence of several species belonging to glycine-betaine-producing genera such as *Bacillus*, *Ectothiorhodospira*, *Escherichia*, *Halomonas*, *Pseudomonas* and *Staphylococcus* (26–28). Since the osmoprotective compatible solute glycine-betaine is known to mitigate a wide range of physicochemical stressors (29), its potential effect on the high-temperature growth/survival of SMMA_5 was tested alongside boron and lithium. Finally, to know whether thermal endurance had a genetic foundation, the whole genome of SMMA_5 was sequenced and analyzed in conjunction with that of its thermally conditioned variant SMMA_5_TC.

## RESULTS

### Taxonomic identity and growth characteristics of SMMA_5

The 16S rRNA gene sequence of SMMA_5 exhibited closest (96.2-97.5%) similarities with homologs from diverse *Paracoccus* species; in the phylogenetic tree constructed, the isolate shared a major clade with *Paracoccus denitrificans*, *Paracoccus kondratievae*, *Paracoccus pantotrophus*, *Paracoccus versutus*, *Paracoccus bengalensis*, and several other species (Fig. S1). In the phylogeny reconstructed based on 92 conserved marker gene sequences (Fig. 1), SMMA_5 clustered in a major clade encompassing *Paracoccus aestuariivivens*, *Paracoccus aminophilus*, *Paracoccus aminovorans*, *P*. *denitrificans*, *P*. *kondratievae*, *Paracoccus laeviglucosivorans*, *Paracoccus limosus*, *Paracoccus litorisediminis*, *Paracoccus lutimaris*, *P*. *pantotrophus*, *Paracoccus sulfuroxidans*, *Paracoccus thiocyanatus*, *P*. *versutus* and *Paracoccus yeei* (notably, no genome sequence was available for *P*. *bengalensis*, so this organism could not be included in the phylogenomic analysis).

**Figure 1.**
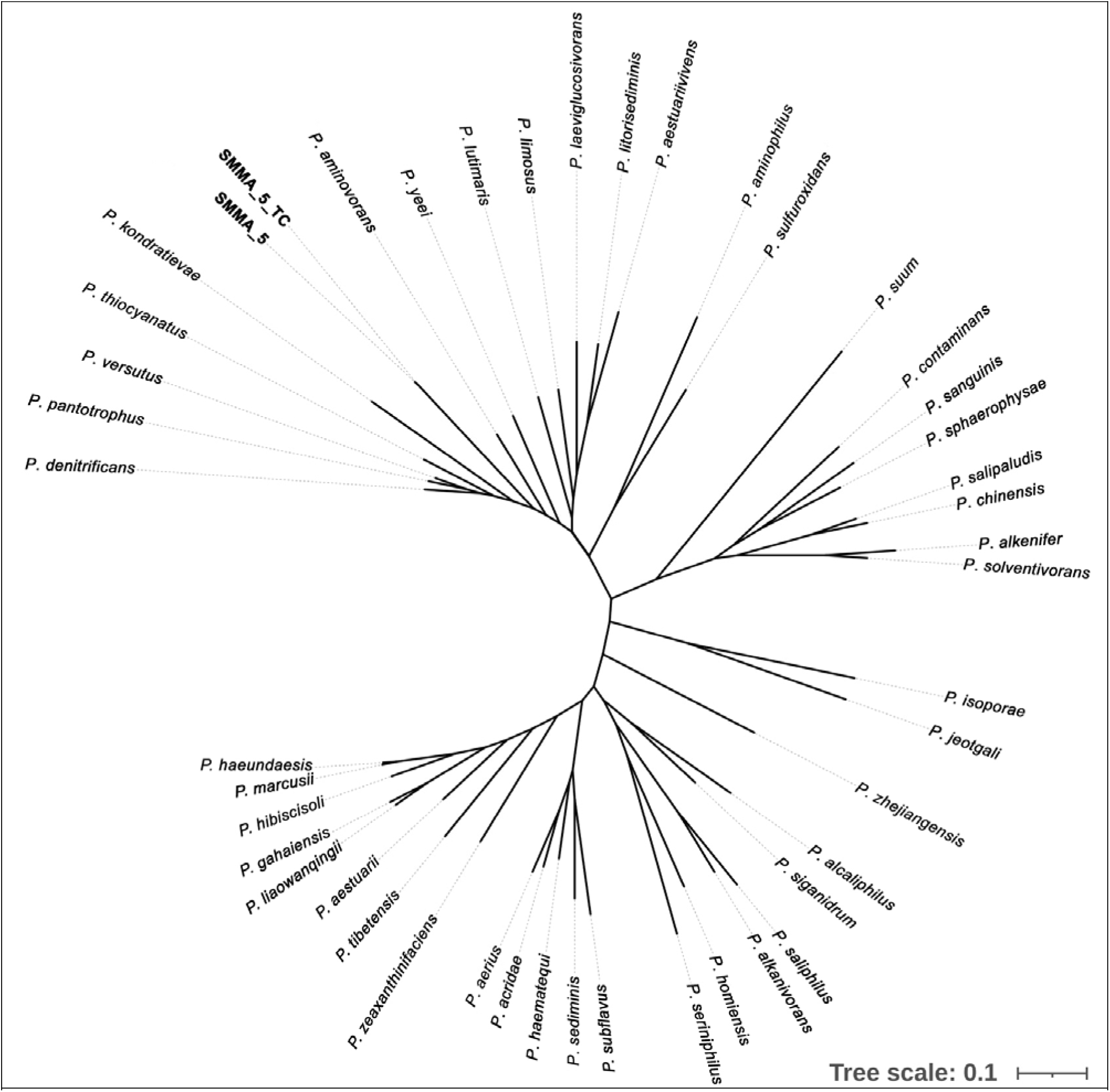
Phylogenetic relationships delineated based on 92 conserved marker gene sequences for SMMA_5, SMMA_5_TC, and 44 other *Paracoccus* species which have near-complete genome sequences available in the GenBank (see Table S1).

The SMMA_5 genome exhibited 23.7%, 23.8% and 21.6% DNA-DNA hybridization, *in silico*, with the genomes of the phylogenetically closest species *P*. *aminovorans*, *P*. *denitrificans* and *P*. *kondratievae* respectively (Table S1). Orthologous genes-based average nucleotide identities of SMMA_5 with the above three genomes were 81.2%, 81.4% and 78.8% respectively. These data indicated that SMMA_5 was a potentially novel species of *Paracoccus*.

When SMMA_5 was incubated in mineral salts thiosulfate (MST) medium for 12 h, >10^4^ times increases in CFU-count were observed at 37°C-45°C; at 50°C, 2% of the initial CFU-count remained in the culture, whereas at 60°C and 70°C no CFU was left (Fig. 2A and B). However, until 45 minutes in MST at 70°C, SMMA_5 had 1.6% of the initial CFU-count remaining in the culture (Fig. 3A and B).

**Figure 2.**
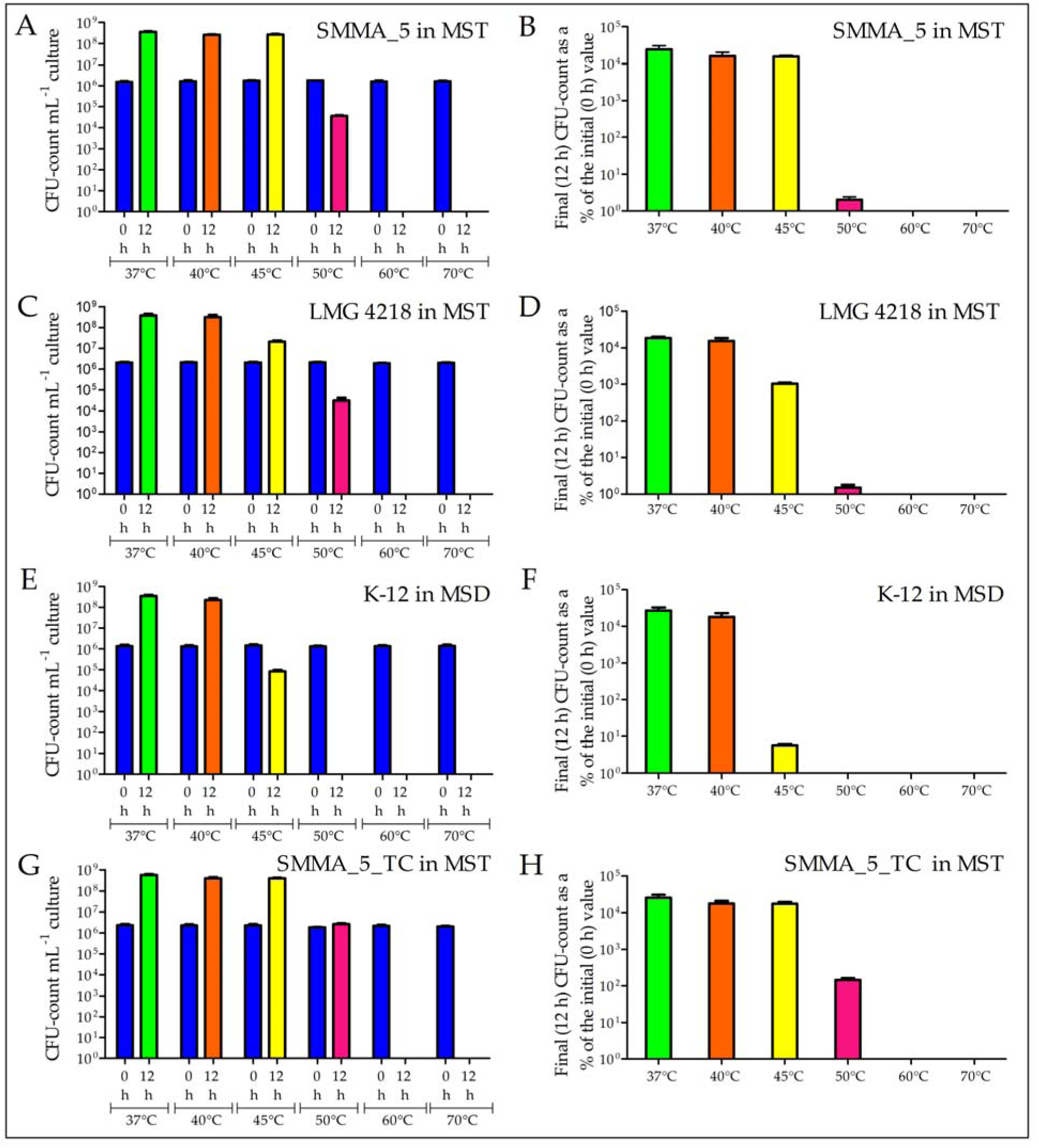
Increase or decrease in the CFU-counts of *Paracoccus* strains SMMA_5, LMG 4218 and SMMA_5_TC in MST, and *E*. *coli* strain K-12 in MSD, after 12 h of incubation at 37°C, 40°C, 45°C, 50°C, 60°C and 70°C. (**A**) 0 h and 12 h CFU-counts for SMMA_5 recorded at the different incubation temperatures; (**B**) representing each final (12 h) CFU-count datum of panel A as a percentage of the corresponding initial (0 h) CFU-count; (**C**) 0 h and 12 h CFU-counts for LMG 4218 recorded at the different incubation temperatures; (**D**) each 12 h CFU-count datum of panel C represented as a percentage of the corresponding 0 h CFU-count; (**E**) 0 h and 12 h CFU-counts for K-12 recorded at the different incubation temperatures; (**F**) each 12 h CFU-count datum of panel E represented as a percentage of the corresponding 0 h CFU-count; (**G**) 0 h and 12 h CFU-counts for SMMA_5_TC recorded at the different incubation temperatures; (**H**) each 12 h CFU-count datum of panel G represented as a percentage of the corresponding 0 h CFU-count. All the data shown in this figure are averages obtained from three different experiments; error bars indicate the standard deviations of the data. Across the panels, and irrespective of the bacterium considered, all 0 h data are represented by blue bars, while 12 h data for incubations at 37°C, 40°C, 45°C, 50°C, 60°C and 70°C are represented by green, orange, yellow, pink, red and purple bars respectively.

**Figure 3.**
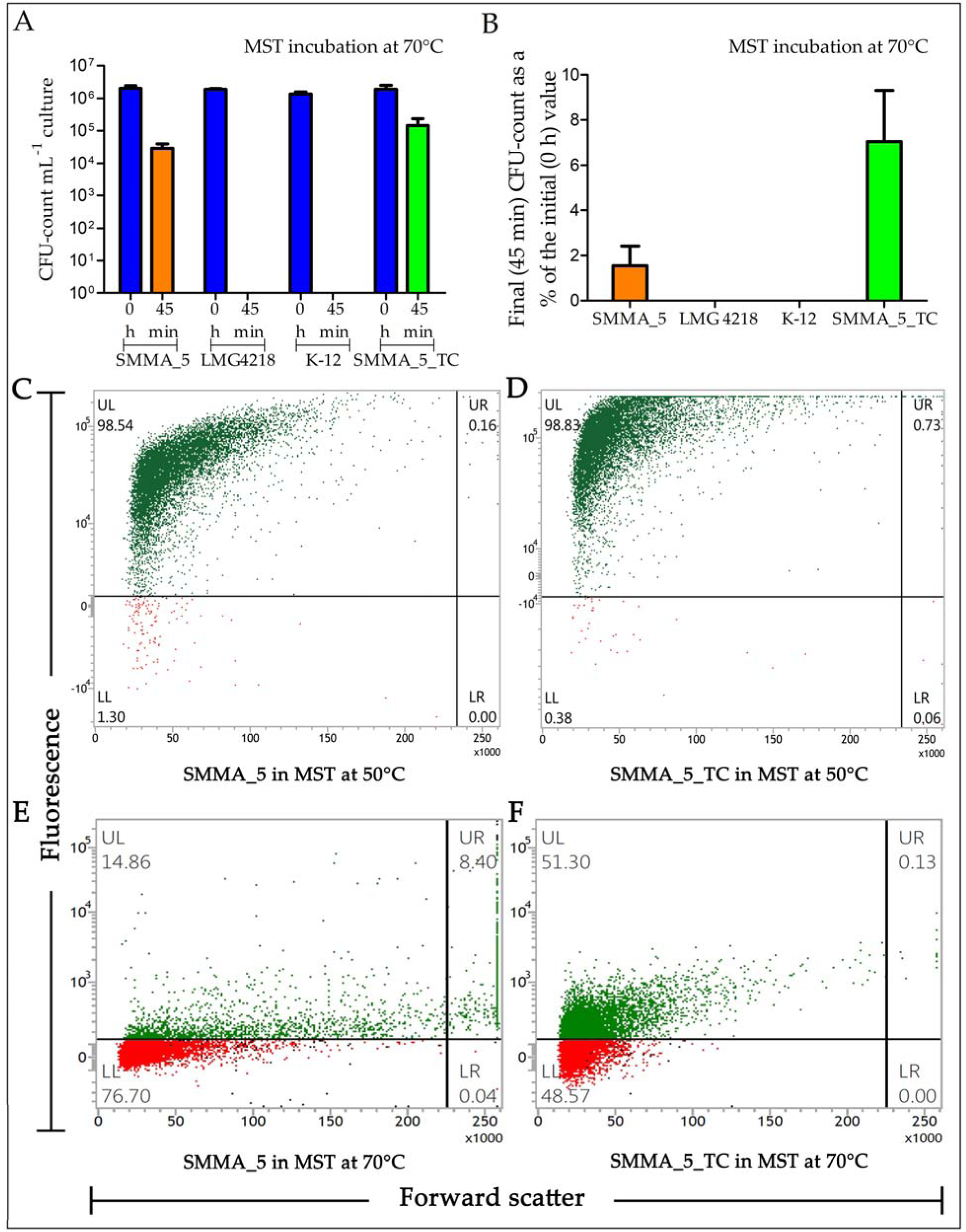
CFU-retention (panels A and B) by *Paracoccus* strains SMMA_5, LMG 4218 and SMMA_5_TC in MST, and *E*. *coli* strain K-12 in MSD, after 45 minutes of incubation at 70°C. Percentages of cells remaining viable (panels C to F) in MST cultures of SMMA_5 and SMMA_5_TC, after 12 h at 50°C and 45 minutes at 70°C, have also been shown. (**A**) number of CFUs present mL^-1^ of the different broth cultures, at 0 h (blue bars for all the cultures tested) and 45 minutes of incubation at 70°C, for SMMA_5 (orange bar), LMG 4218 (purple bar), K-12 (yellow bar) and SMMA_5_TC (green bar); (**B**) percentages of the initial CFU-counts that were retained in the SMMA_5, LMG 4218, SMMA_5_TC and K-12 after 45 minutes of incubation at 70°C (strain wise color code for the bars are same as that used in A). The data shown in panels A and B are averages obtained from three different experiments; error bars indicate the standard deviations of the data. Flow-cytometry-based dot plots showing the proportions of FDA-stained and FDA-unstained cells for (**C**) SMMA_5 after 12 h at 50°C, (**D**) SMMA_5_TC after 12 h at 50°C, (**E**) SMMA_5 after 45 minutes at 70°C, and (**F**) SMMA_5_TC after 45 minutes at 70°C. In panels C to F, fluorescence versus forward scatter data for 10000 randomly- taken cells have been potted; green and red dots represent cells that were stained and not stained by FDA respectively. Each flow-cytometry-based experiment was repeated for two more occasions and in every instance <2% deviations were observed from the values shown here for the proportions of FDA-stained and FDA-unstained cells.

The comparator chemolithoautotroph *P*. *pantotrophus* LMG 4218, when incubated in MST for 12 h, exhibited >10^4^ times increase in CFU-count at up to 40°C; at 45°C ∼10^3^ times increase in CFU-count was recorded, while at 50°C, 1.5% of the initial CFU-count remained in the culture; at 60°C and 70°C no CFU was left (Fig. 2C and D); no CFU was also there after 45 minutes in MST at 70°C (Fig. 3A and B).

*Escherichia coli* K-12, when incubated in mineral salts dextrose (MSD) for 12 h, showed >10^4^ times CFU-growth at up to 40°C; whereas only 5.8% CFU remained in the culture at 45°C, no CFU was left at ≥50°C (Fig. 2E and F); no CFU was also there after 45 minutes at 70°C (Fig. 3A and B).

### Thermal conditioning enhances high-temperature growth/survival

For maintenance, SMMA_5 was grown for 12 h at 37°C, and stored at 4°C for 28 days until the next transfer. On the other hand, its thermally conditioned cell-line (sibling strain), designated as SMMA_5_TC, was raised and maintained by first incubating for 12 h at 50°C, then reviving growth for 12 h at 37°C, and finally storing at 25°C for 14 days until the next transfer involving the same 50°C-37°C incubation procedure (Fig. 4A). All data presented for SMMA_5 and SMMA_5_TC were obtained from experiments carried out after 15 and 30 transfer cycles respectively.

**Figure 4.**
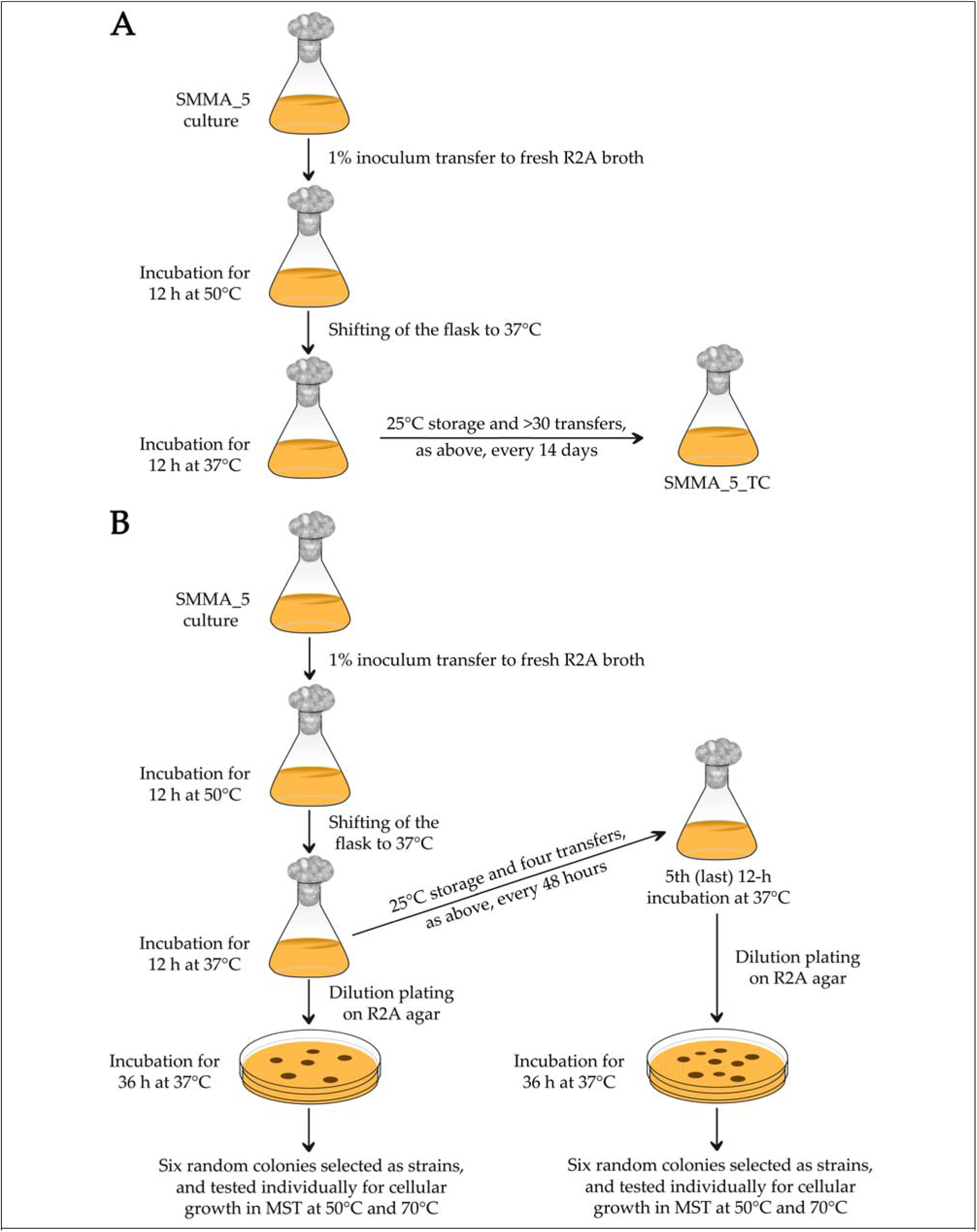
Schematic diagrams showing the procedure followed for thermal conditioning of SMMA_5. (**A**) steps followed in the creation of SMMA_5_TC; (**B**) steps followed in raising sibling strains of SMMA_5 after one or five cycles (generations) of thermal conditioning.

When tested for growth in MST medium at 37°C, 40°C and 45°C, SMMA_5_TC (Fig. 2G and H) exhibited phenotypes identical to those of SMMA_5 (Fig. 2A and B). However, after 12 h at 50°C in MST, SMMA_5_TC exhibited ∼1.5 times increase in the CFU-count of the culture with respect to the 0 h level (Fig. 2G and H), whereas under identical conditions SMMA_5 retained only 2% of the initial CFU-count (Fig. 2A and B). Albeit, SMMA_5_TC (Fig. 2G and H) did not have any CFU left after 12 h in MST at 60°C and 70°C, after 45 minutes in MST at 70°C it had 7% of the initial CFU-count intact (Fig. 3A and B).

Subsequently, experiments were carried out to know the effects of only a few thermal-conditioning cycles on the growth or CFU-retention by SMMA_5 at 50°C and 70°C (each cycle involved incubation for 12 h at 50°C, followed by 12 h at 37°C; Fig. 4B). When the six sibling strains generated after one round of thermal conditioning were individually incubated for 12 h at 50°C, 2.5%, 4.3%, 1.7%, 3.7%, 5.8% and 2.9% (on an average 3.5%) of their initial CFU-counts were retained (Fig. 5A and B). Likewise, when these six sibling strains were individually incubated for 45 minutes at 70°C, 3.8%, 1.8%, 4.2%, 3.2%, 2.7% and 4.0% (on an average 3.3%) of their initial CFU-counts were retained (Fig. 5E and F).

**Figure 5.**
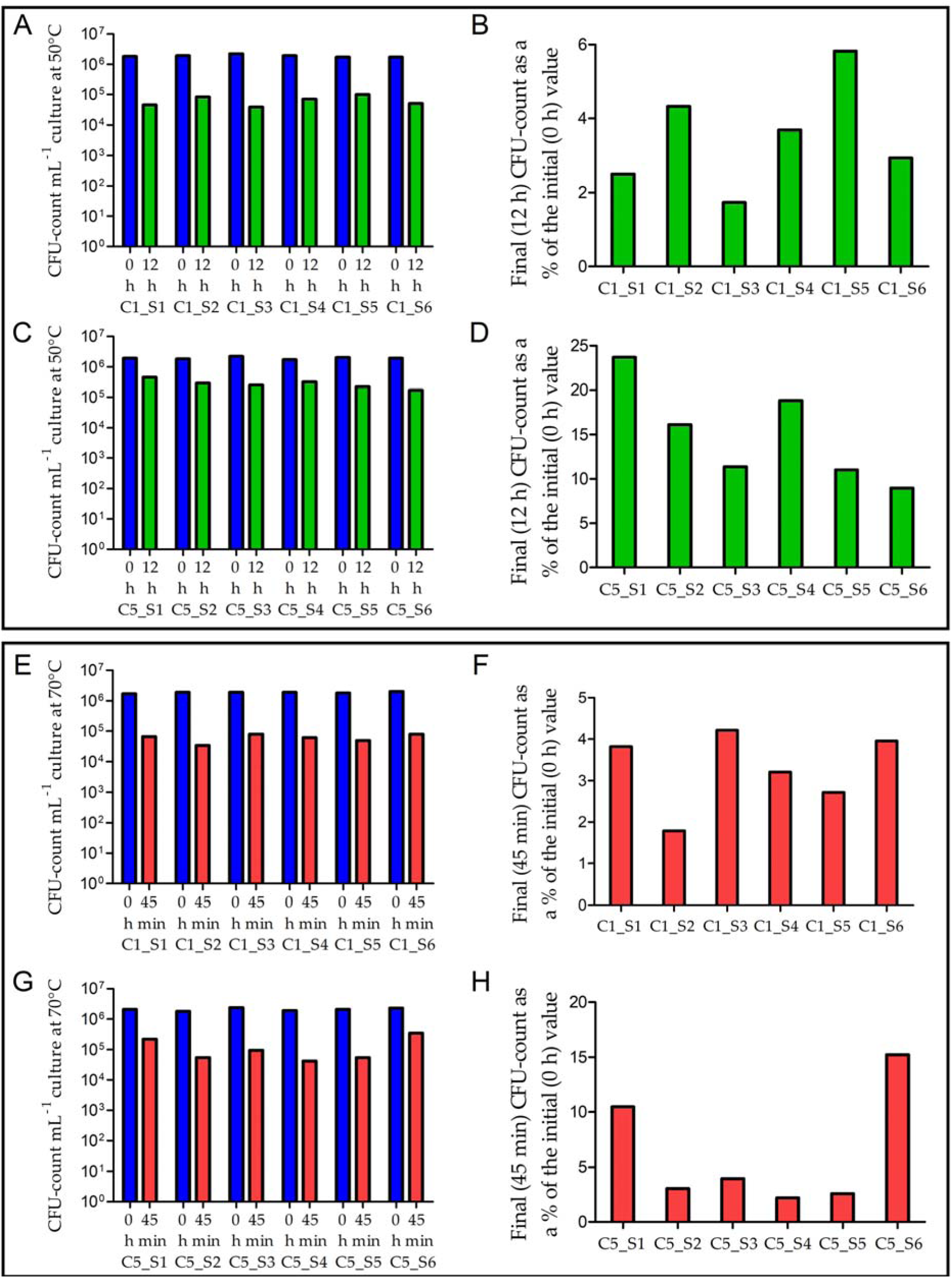
CFU-retention at 50°C (panels A to D) and 70°C (panels E to H) by multiple sibling strains of SMMA_5 generated after one (panels A, B, E and F) or five (panels C, D, G and H) cycles (generations) of thermal conditioning. (**A**) 0 h and 12 h CFU-counts in MST at 50°C for the six sibling strains of SMMA_5 (C1_S1 to C1_S6) that were raised via a single cycle of thermal conditioning; (**B**) final (12 h) CFU-count of each sibling strain shown in panel A, as a percentage of the corresponding initial (0 h) CFU-count; (**C**) 0 h and 12 h CFU-counts in MST at 50°C for the six sibling strains of SMMA_5 (C5_S1 to C5_S6) that were raised via five cycle of thermal conditioning; (**D**) final (12 h) CFU-count of each sibling strain shown in panel C, as a percentage of the corresponding initial (0 h) CFU-count; (**E**) 0 h and 45 minutes CFU-counts in MST at 70°C for the six sibling strains of SMMA_5 (C1_S1 to C1_S6) that were raised via a single cycle of thermal conditioning; (**F**) final (45 minutes) CFU-count of each sibling strain shown in panel E, as a percentage of the corresponding initial (0 h) CFU-count; (**G**) 0 h and 45 minutes CFU-counts in MST at 70°C for the six sibling strains of SMMA_5 (C5_S1 to C5_S6) that were raised via five cycle of thermal conditioning; (**H**) final (45 minutes) CFU-count of each sibling strain shown in panel G, as a percentage of the corresponding initial (0 h) CFU-count. Across the panels, 0 h, 12 h and 45 minutes data are represented by blue, green and red bars respectively.

When the six sibling strains generated after five rounds of thermal conditioning were individually incubated for 12 h at 50°C, 23.7%, 16.1%, 11.4%, 18.8%, 11.0% and 8.9% (on an average 15%) of their initial CFU-counts were retained (Fig. 5C and D). Furthermore, when these six sibling strains were individually incubated for 45 minutes at 70°C, 10.5%, 3.1%, 4.0%, 2.2%, 2.6% and 15.2% (on an average 6.2%) of their initial CFU-counts were retained (Fig. 5G and H).

### At high temperatures, cell viability is more than divisibility

After 12 h at 50°C, as well as after 45 minutes at 70°C, the percentages of SMMA_5 and SMMA_5_TC cells remaining viable or metabolically active were far higher than the percentages of cells remaining ready to divide and form single colonies (CFU-count data). As a mark of their metabolically active state, cell populations were tested post high-temperature incubation for their ability to uptake FDA and hydrolyze it to fluorescein, which in turn was detected via flow cytometry. In that way, after 12 h incubation at 50°C in MST, 98.7% and 99.56% cells of SMMA_5 and SMMA_5_TC were found to remain viable respectively (Fig. 3C and D); similarly, 23.26% and 51.43% cells of SMMA_5 and SMMA_5_TC remained viable after 45 minutes at 70°C in MST (Fig. 3E and F).

### Environmental solutes enhance thermal endurance or elicit moderate thermophilicity

When SMMA_5 was incubated at 50°C for 12 h in MSTB, MSTL and MSTG, i.e. MST supplemented with 16 mM boron, 1 mM lithium and 10 mM glycine-betaine respectively, 14.4%, 29.2% and 224% of the initial CFU-counts were present in the respective cultures (Fig. 6C and S2C). For SMMA_5_TC, 12 h incubation at 50°C in MSTB, MSTL and MSTG led to the CFU-growths equivalent to 170%, 230% and 370% of the initial counts respectively (Fig. 6D and S2D). Notably, SMMA_5 and SMMA_5_TC were also found to release 8-10 mM glycine-betaine in the medium after 12 h incubation in MST at 50°C.

**Figure 6.**
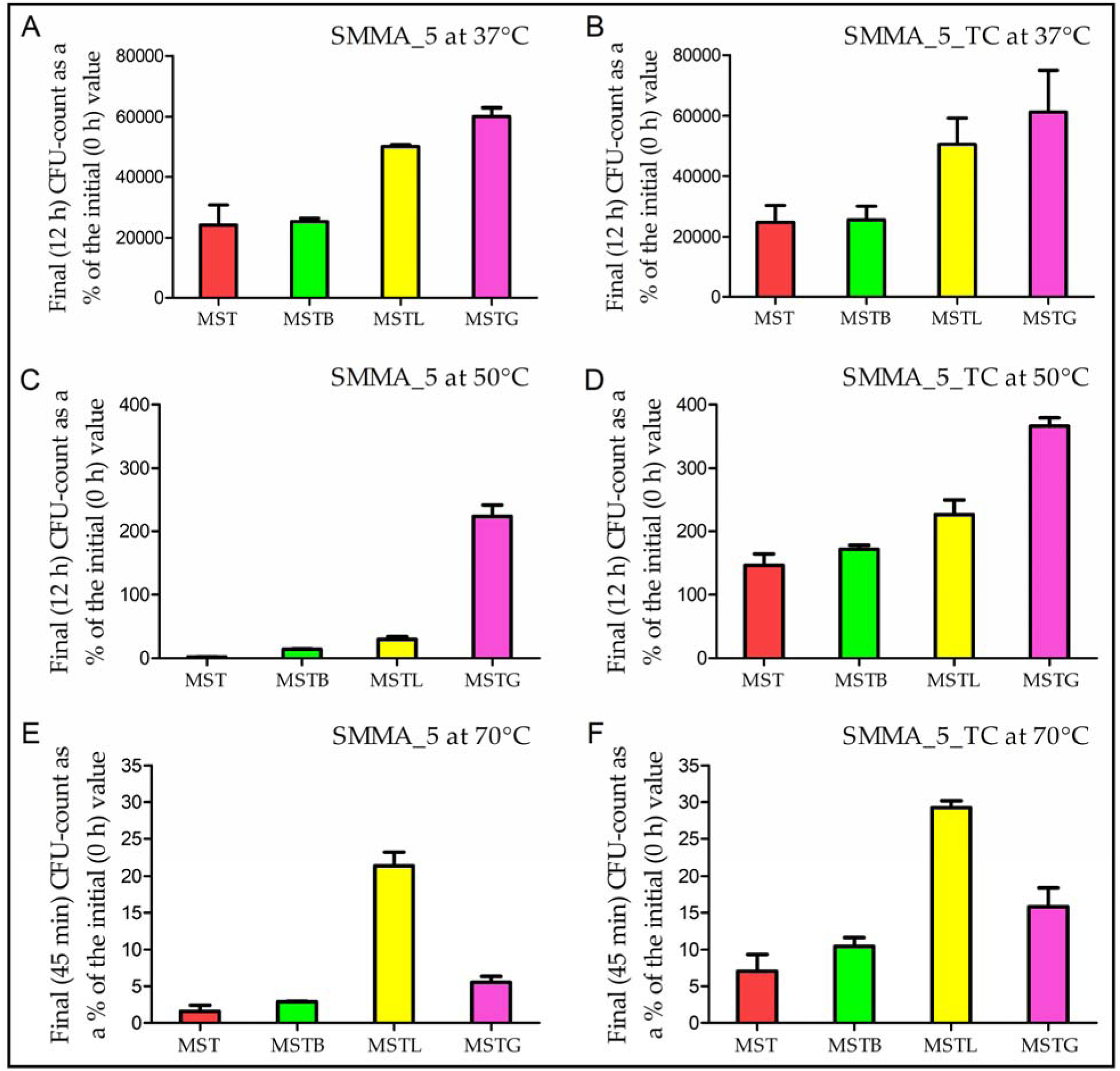
Increase or decrease in the CFU-count of SMMA_5 (panels A, C and E) and SMMA_5_TC (panels B, D and F) in MST, MSTB, MSTL and MSTG media after incubation at 37°C (panels A and B), 50°C (panels C and D) and 70°C (panels E and F). (**A**) final (12 h) CFU-counts of SMMA_5 in the different media at 37°C, represented as percentages of the corresponding initial (0 h) CFU-counts; (**B**) final (12 h) CFU-counts of SMMA_5_TC in the different media at 37°C, represented as percentages of the corresponding initial (0 h) CFU-counts; (**C**) final (12 h) CFU-counts of SMMA_5 in the different media at 50°C, represented as percentages of the corresponding initial (0 h) CFU-counts; (**D**) final (12 h) CFU-counts of SMMA_5_TC in the different media at 50°C, represented as percentages of the corresponding initial (0 h) CFU-counts; (**E**) final (45 minutes) CFU-counts of SMMA_5 in the different media at 70°C, represented as percentages of the corresponding initial (0 h) CFU-counts; (**F**) final (45 minutes) CFU- counts of SMMA_5_TC in the different media at 70°C, represented as percentages of the corresponding initial (0 h) CFU-counts. The data shown in panels A to F are averages obtained from three different experiments; error bars indicate the standard deviations of the data. Irrespective of the incubation temperature, data recorded for MST, MSTB, MSTL and MSTG media are represented by red, green, yellow and purple bars respectively.

After 45 minutes at 70°C in MSTB, MSTL or MSTG, SMMA_5 retained 2.9%, 21.4% and 5.5% of the initial CFU-count respectively (Fig. 6E and S2E). For SMMA_5_TC, 45 minute incubation at 70°C in MSTB, MSTL and MSTG led to the retention of 10.4%, 29.2% and 15.8% of the initial CFU-count respectively (Fig. 6F and S2F). Furthermore, 37°C incubation of either strain for 12 h in MSTB, MSTL and MSTG showed that boron, lithium and glycine-betaine were not only non-toxic to them but also considerably stimulatory to their growth at this temperature (Fig. 6A and B; Fig. S2A and B).

The three environmental solutes also exhibited pronounced influence on the viability of SMMA_5 and SMMA_5_TC cells at 70°C (notably, at 50°C, almost 100% cell viability was already recorded for both strains, without any environmental solute in the medium). After incubation in MSTB, MSTL and MSTG for 45 minutes at 70°C, 43.45%, 71.36% and 55.17% cells of SMMA_5 were found to remain viable respectively (Fig. 7A-C). Likewise, after 45 minutes in MSTB, MSTL and MSTG at 70°C, 66.04%, 89.31% and 76.35% cells of SMMA_5_TC were found to remain viable respectively (Fig. 7D to F).

**Figure 7.**
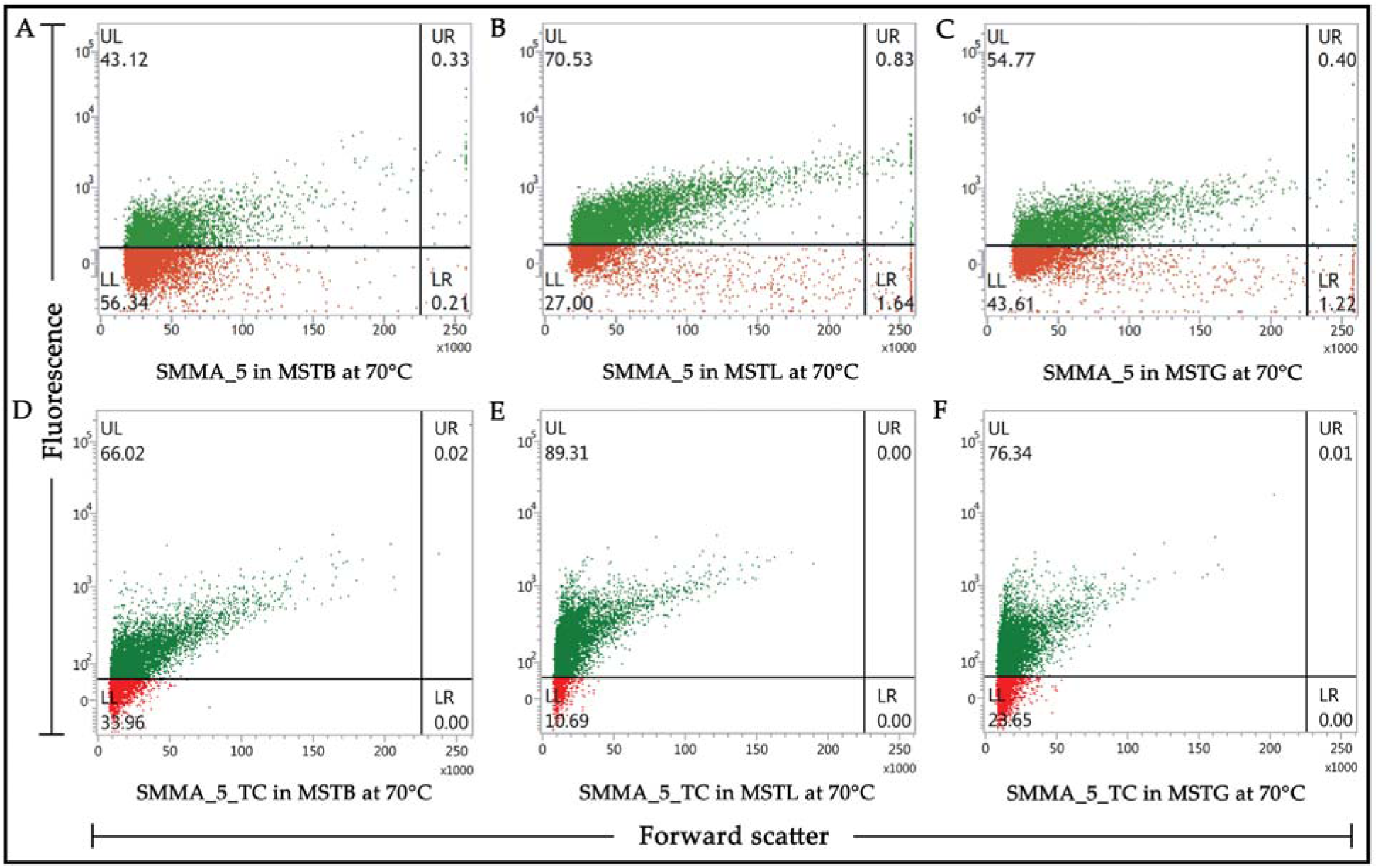
Flow-cytometry-based dot plots showing what percentage of cells remained viable (i.e. the proportion of FDA-stained and FDA-unstained cells) for SMMA_5 (panels A to C) and SMMA_5_TC (panels D to F), after 45 minute incubation at 70°C, in MSTB (panels A and D), MSTL (panels B and E) and MSTG (panels C and F). In each panel, fluorescence versus forward scatter data for 10000 randomly-taken cells have been potted; green and red dots represent cells that were stained and not stained by FDA respectively. Each experiment was repeated for two more occasions and in every instance <2% deviations were observed from the values shown here for the proportions of FDA-stained and FDA-unstained cells.

### Oligotrophy helps endure high temperature

When incubated in R2A for 12 h at 37°C, both SMMA_5 and SMMA_5_TC exhibited >10^4^ times increase in CFU-count (Fig. 8A and B; Fig. S3A and B); comparable growth was recorded for the two strains in Luria broth after 12 h at 37°C (data not shown). After 12 h at 50°C in R2A, SMMA_5 and SMMA_5_TC retained 2% and 2.5% of the initial CFU-counts respectively (Fig. 8C and D; Fig. S3C and D), but after 12 h at 50°C in Luria broth, neither strain retained any CFU. Likewise, after 45 minutes at 70°C in R2A, SMMA_5 and SMMA_5_TC retained 0.5% and 1.0% of the initial CFU-counts respectively (Fig. 8E and F; Fig. S3E and F), but their Luria broth cultures retained no CFU after 45 minutes at 70°C.

**Figure 8.**
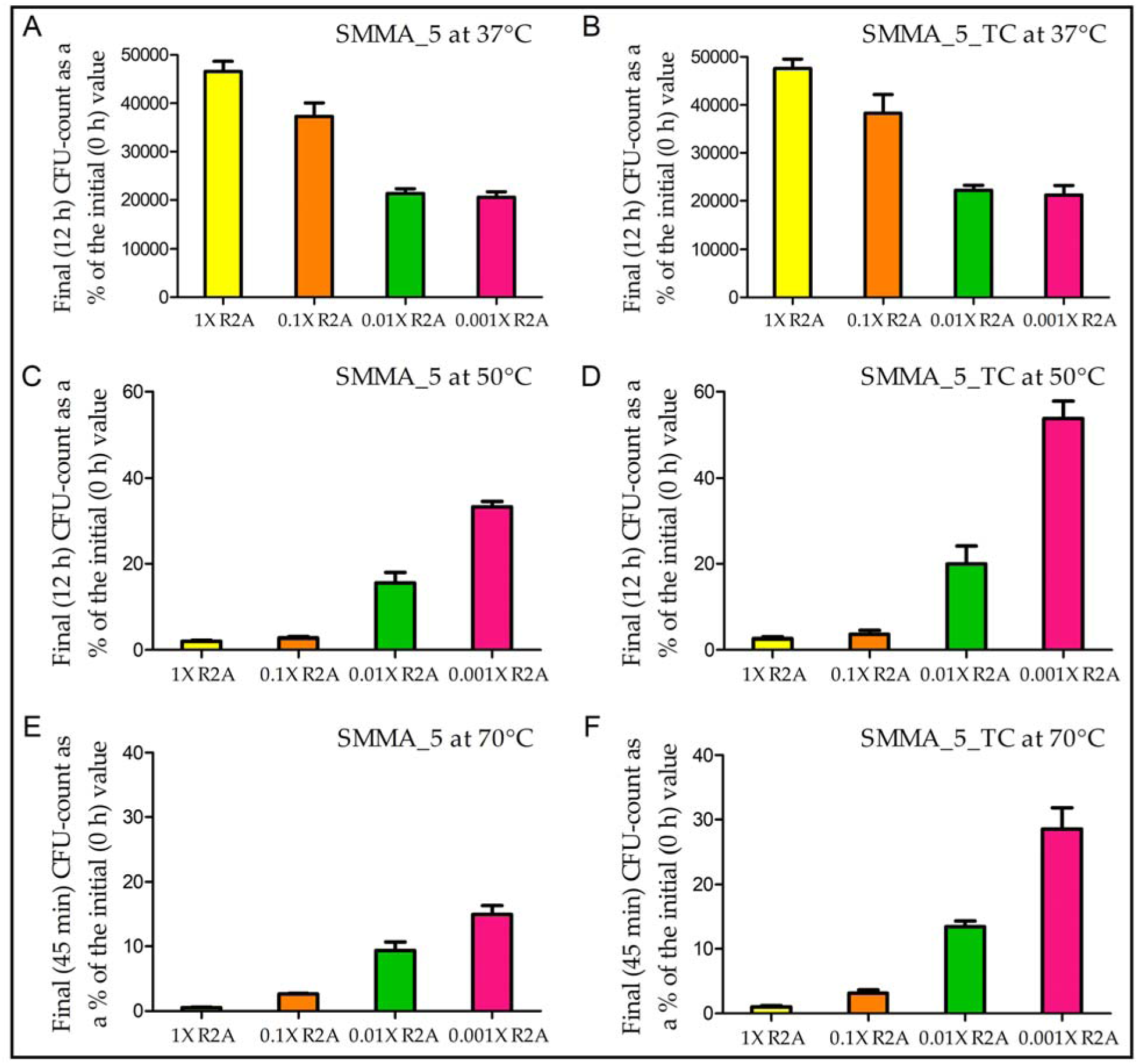
Increase or decrease in the CFU-count of SMMA_5 (panels A, C and E) and SMMA_5_TC (panels B, D and F) in 1X R2A, 0.1X R2A, 0.01 XR2A and 0.001 XR2A media after incubation at 37°C (panels A and B), 50°C (panels C and D) and 70°C (panels E and F). (**A**) final (12 h) CFU-counts of SMMA_5 in the different media at 37°C, represented as percentages of the corresponding initial (0 h) CFU-counts; (**B**) final (12 h) CFU-counts of SMMA_5_TC in the different media at 37°C, represented as percentages of the corresponding initial (0 h) CFU-counts; (**C**) final (12 h) CFU-counts of SMMA_5 in the different media at 50°C, represented as percentages of the corresponding initial (0 h) CFU-counts; (**D**) final (12 h) CFU-counts of SMMA_5_TC in the different media at 50°C, represented as percentages of the corresponding initial (0 h) CFU-counts; (**E**) final (45 minutes) CFU-counts of SMMA_5 in the different media at 70°C, represented as percentages of the corresponding initial (0 h) CFU-counts; (**F**) final (45 minutes) CFU-counts of SMMA_5_TC in the different media at 70°C, represented as percentages of the corresponding initial (0 h) CFU-counts. The data shown in panels A to F are averages obtained from three different experiments; error bars indicate the standard deviations of the data. Irrespective of the incubation temperature, data recorded for 1X R2A, 0.1X R2A, 0.01X R2A and 0.001X R2A media are represented by yellow, orange, green and pink bars respectively.

At 37°C, 12 h incubation in 0.1X, 0.01X and 0.001X R2A resulted in progressively lower cellular growth yields, as compared to what was recorded in R2A (1X), for both SMMA_5 and SMMA_5_TC (Fig. 8A and B; Fig. S3A and B).

After 12 h incubation at 50°C in 0.1X, 0.01X and 0.001X R2A, SMMA_5 retained 2.7%, 15.6% and 33.3% of the initial CFU-count respectively (Fig. 8C and S3C), whereas SMMA_5_TC retained 3.5%, 20.0% and 53.8% of the initial CFU-count respectively (Fig. 8D and S3D).

After 45 minutes at 70°C in 0.1X, 0.01X and 0.001X R2A, SMMA_5 cultures retained 2.6%, 9.4% and 14.9% of the initial CFU-count respectively (Fig. 8E and S3E), whereas SMMA_5_TC retained 3.1%, 13.4% and 28.5% of the initial CFU-count respectively (Fig. 8F and S3F).

### pH 7.5 is optimum for thermal endurance

The optimum pH for growth or CFU-retention at 37°C was tested in R2A, while the same at 50°C and 70°C was tested in 0.001X R2A. The media types were chosen based on their best support for growth or CFU-retention at the respective temperatures (Fig. 8); pH optima were not determined in MST medium because its salt ingredients precipitate out at pH >7.0 and temperature ≥50°C. For both SMMA_5 and SMMA_5_TC, R2A having pH 7.5 supported maximum growth after 12 h at 37°C (Fig. S4A and B; Fig. S5A and B); likewise, 0.001X R2A having pH 7.5 supported maximum CFU-retention after 12 h at 50°C (Fig. S4C and D; Fig. S5C and D), as well as 45 minutes at 70°C (Fig. S4E and F; Fig. S5E and F).

### Rapid growth is not sustainable amid high heat

SMMA_5_TC, when grown at 70°C in 0.001X R2A having an initial pH 7.5 and fortified simultaneously with 4 mM Na_2_B_4_O_7_.10H_2_O, 1 mM LiOH.H_2_O and 10 mM C_5_H_11_NO_2_, exhibited a 1.4 time increase in CFU-count (from 1.9 × 10^6^ mL^-1^ to 2.73 × 10^6^ mL^-1^, as averages of three different experiments where standard deviations of the data were <2% of the means) after 45 minutes of incubation; but after 1 h, no CFU (or for that matter FDA-stained cell) was left in the culture.

### Genomic uniqueness of the hot spring *Paracoccus*

The 3.09 Mb and 3.06 Mb draft genomes of SMMA_5 and SMMA_5_TC, as represented in 283 and 247 contigs (all >200 bp long), had G+C contents of 65.78% and 65.83% respectively (Table S1). SMMA_5 contained 3,234 potential open reading frames (ORFs) or coding sequences (CDSs), out of which 2498 and 503 encoded known and unknown (hypothetical) proteins respectively; 16S, 23S, and 5S rRNA genes were identified together with 49 tRNA genes. The completeness level of this genome was estimated to be 98.23% (Table S1), based on the presence of 556, out of the total 568, *Paracoccus*-specific conserved marker genes curated in CheckM database. The draft genome of SMMA_5_TC, on the other hand, was found to include 3,163 potential CDSs, out of which 2511 and 477 encoded known and unknown proteins respectively; 16S, 23S, and 5S rRNA genes were identified alongside 47 tRNA genes. Completeness of the SMMA_5_TC genome was 98.41% (Table S1), based on the presence of 554 *Paracoccus*-specific marker genes. The average isoelectric point (pI) of all the putative proteins encoded by SMMA_5 and SMMA_5_TC were predicted as 6.86 and 6.87 respectively; corresponding values for all other *Paracoccus* species except *Paracoccus contaminans* were lower (Table S2; Fig. S6).

Pan genome analysis (PGA) involving SMMA_5, and 44 other *Paracoccus* species having near-complete genome sequences available in the GenBank (Table S1), revealed 822 core genes for the genus. While SMMA_5 had 1742 accessory, 335 unique and 30 exclusively-absent genes, the corresponding numbers in the comparators ranged between 1528-3451, 159-817 and 0-45 (Table S3). Of the 335 unique genes of SMMA_5, 186 encoded hypothetical proteins, whereas 149 coded for putative functional proteins, of which, again, only 83 were ascribable to one or more clusters of orthologous group (COG) categories: predominantly, DNA replication, recombination and repair; transcription; secondary metabolites biosynthesis, transport and catabolism; inorganic ion transport and metabolism; and cell wall/membrane/envelope biogenesis (Table S4).

The metabolic potentials of SMMA_5 were compared with those of the phylogenomically closest *Paracoccus* species (see Fig. 1) on the basis of their gene-content under different COG categories. Two-way clustering of the genomes analyzed and COG categories identified revealed close functional relationships of SMMA_5 with *P. dinitrificans* and *Paracoccus thiocyanatus* on one hand, and *P. aminophilus* and *Paracoccus sulfurioxidans* on the other (Fig. 9A). Notably, in the phylogenomic tree, the first two species were closely related to SMMA_5, but the second pair diverged widely (Fig. 1). *P*. *pantotrophus P*. *yeei*, *P*. *kondratievae*, *P*. *aminovorans* and *P*. *lutimaris* showed the next closest relationships with SMMA_5 in the two-way dendogram (Fig. 9A), but out of these five species only the first four shared the same branch with SMMA_5 in the phylogenomic tree whereas *P*. *lutimaris* was in a completely distinct branch (Fig. 1). *P*. *versutus*, which was phylogenomically closely related to SMMA_5 (Fig. 1), diverged widely from the new isolate in terms of gene-content under different COG categories (Fig. 9A). Statistically analysis further revealed the two COG categories (i) cell wall/membrane/envelope biogenesis, and (ii) amino acid transport and metabolism, to be exclusively enriched in SMMA_5; lipid transport and metabolism was found to be enriched in SMMA_5 and *P. aminovorans* (Fig. 9B).

**Figure 9.**
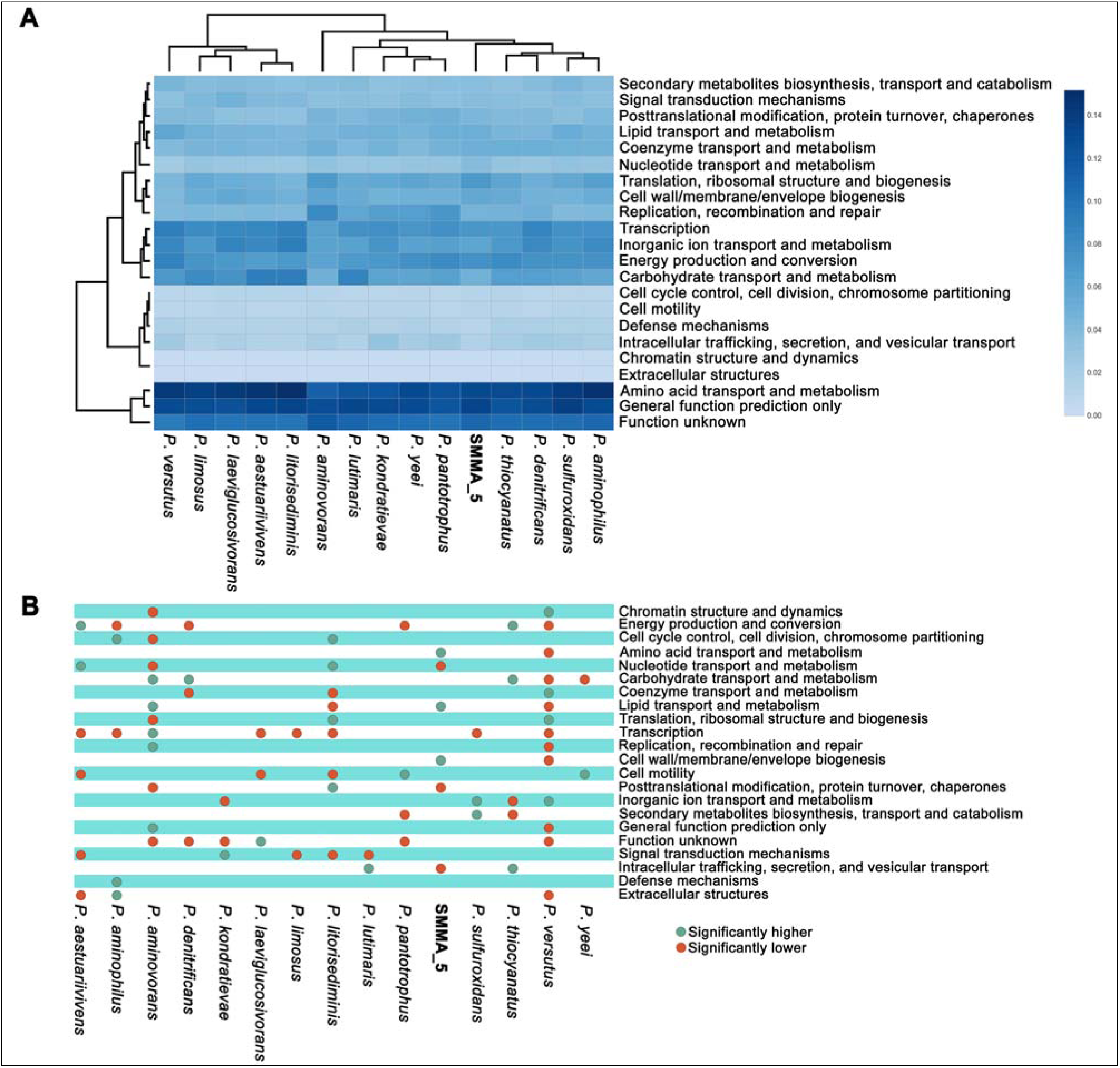
Functional analysis of the genomes of SMMA_5 and its phylogenetically closest *Paracoccus* species (the latter were identified based on the phylogeny of 92 conserved marker gene sequences, Fig. 1). (**A**) Heat map comparing the richness of the metabolic/functional categories across the genomes, determined in terms of the number of Clusters of Orthologous Groups (COGs) of Proteins that are ascribed to the categories in individual genomes; a two-dimensional clustering is also shown, involving 15 bacterial genomes on one hand and the 22 functional categories of COGs on the other; color gradient of the heat map varied from high (deep blue) to low (faint blue), through moderate (light blue), richness of the categories across the genomes. (**B**) Statistically significant, high (green circles) or low (red circles) richness of the functional categories of COG, detected across the genomes via Chi Square test with p < 0.001.

### Genomic aspects potentially linked to thermal adaptation

The genomes of both SMMA_5 and SMMA_5_TC encompassed a large repertoire of heat shock proteins and molecular chaperones. These included (i) one copy each of the co-chaperone GroES, the heat shock protein HspQ, the heat-inducible transcriptional repressor HrcA, the Hsp33 family molecular chaperone HslO, the co-chaperone GrpE, and the RNA polymerase sigma 32 factor RpoH; (ii) two copies each of the genes encoding the chaperonin GroEL, Hsp20 family protein, and the Hsp70 family protein DnaK; and (iii) three copies of the molecular chaperone DnaJ (Hsp40). Genes for heat shock protein Hsp90 and RNA polymerase sigma E factor, which are typically associated with heat stress management in *E*. *coli*, were not found in the hot spring *Paracoccus* genomes (Table S5). On the other hand, the presence of genes for the redox-regulated, Hsp33-family chaperone HslO – which is present in most thermophilic bacteria to protect thermally/oxidatively damaged proteins from irreversible aggregation – distinguished the genomes of SMMA_5 and SMMA_5_TC from those of *E*. *coli* and *P*. *denitrificans*. Furthermore, the genomes of SMMA_5 and SMMA_5_TC encoded several other proteins that can be potentially decisive in thermal adaptation by means of their regulated expression (Table S6).

The draft genomes of SMMA_5 and SMMA_5_TC showed 99.9748% sequence similarity over a total alignment length of 3061077 nucleotides (unaligned portions of the two genomes were attributable to their incomplete sequencing, rather than the presence of unique genes). Of the total 43 instances of single nucleotide polymorphism (SNP) detected across the alignment, 18 were in protein-coding genes, 18 were in frame- shifted pseudogenes, whereas seven were in non-coding regions (Table S7). Out of the 18 nucleotide substitutions detected in protein-coding genes, 13 were nonsynonymous (ns) SNPs while five were synonymous (silent) mutations (sSNPs). Of the 13 nsSNPs, again, nine and four involved transitions and transversions respectively, while seven involved such radical replacements of amino acids (e.g., polar to non-polar, or aromatic-group-containing to alkyl-group-containing, amino acids, or vice versa) that may lead to changes in the corresponding protein structure and function (Table S7).

## DISCUSSIONS

A complex set of geobiological and biophysical factors worked in conjunction with specialized genomic attributes to confer thermal endurance, or even moderate thermophilicity, to the hot-spring-dwelling relative of mesophilic *Paracoccus*. While the thermally unconditioned isolate SMMA_5 had an intrinsically higher frequency of CFU- retention at 50°C and 70°C, compared with other *Paracoccus* (and *E*. *coli*), its sibling SMMA_5_TC, which had been thermally conditioned for more than 30 transfer cycles, showed either growth or yet higher frequencies of CFU-retention at these temperatures. Corroboratively, thermal conditioning for only five transfer cycles brought about, on an average, eight and four times increases in the CFU-retention frequencies of SMMA_5 at 50°C and 70°C respectively. Geochemical and microbial solutes further improved the growth/survival efficacies at 50°C and 70°C, as did extreme oligotrophy at pH 7.5.

Overall it seems that during their residence in the solute-poor, circum-neutral pH, and near-boiling water of Lotus Pond (7, 12), cell populations of this novel *Paracoccus* had already acquired such heritable structural and functional attributes which are conducive for the mitigation of high heat. *Ex situ*, the potential cellular memories conferring heat endurance (or moderate thermophilicity) were on the wane in the thermally unconditioned culture (SMMA_5), while the same were retained relatively better in the thermally conditioned variant (SMMA_5_TC).

### Prior experience of high temperature enhances heat endurance or confers moderate thermophilicity

Pre-exposure to sub-lethal stress conditions is known to enhance a microorganism’s resilience to higher levels of stress (30). Bacteria are endowed with diverse memory mechanisms, including those which involve the inheritance and propagation of epigenetic states associated with adaptive advantages (31, 32). Of the various metabolic systems that are known to involve heritable epigenetic memory, ion-channel-mediated signaling and ion flux machineries (33–35) appear to be pertinent to thermal endurance in view of the observed influence of environmental solutes on growth and survival at high temperatures.

Under almost all the culture conditions tested, the standard deviations of the CFU- retention data for SMMA_5_TC were higher than those for SMMA_5. Concurrent to this trend, the sibling strains generated via one or five cycles of thermal conditioning of SMMA_5, did not show mutually equal frequencies of CFU-retention at any of the high-temperatures tested. These data collectively indicated that in the hot-spring-dwelling *Paracoccus*, thermal conditioning via potential trait memorization could be a multifactorial, and asymmetrically inherited (32), process that involves a “bet and hedging” strategy, where heterogeneous populations of phenotypically-distinct but genotypically-identical cells are formed and maintained by a bacterium to ensure that one sub-population or the other can always withstand, and adapt to, unknown environmental challenges of the future (36).

### Role of environmental solutes in enhancing thermal endurance or eliciting moderate thermophilicity

Out of the three environmental solutes tested, the alkali metal lithium was most effective in increasing the frequency of CFU-retention, as well as the percentage of viable cells, at 70°C, for both SMMA_5 and SMMA_5_TC, as compared to their respective MST phenotypes. At 50°C too, lithium fortification of MST increased the frequency of CFU- retention and CFU-growth for SMMA_5 and SMMA_5_TC respectively. These phenotypes were consistent with previous observations where *Listeria monocytogenes* exposed to 62.8°C for 10-20 minutes was recovered after 48-144 h, by adding 7 g L^-1^ LiCl to the 30°C revival cultures (18). Thermoprotective effect of lithium could be linked, but not necessarily restricted, to its activities at the periphery of cells. Li^+^ efficiently forms hydrogen bonds with nearby water molecules, as well as charged surfaces such as cell membranes, thereby modifying their intrinsic van der Waals, and electrostatic, forces (37). Li^+^ also competes for ligand-binding sites of biomacromolecules that are otherwise reserved for Na^+^ or Mg^2+^ (38). By virtue of these biophysical maneuvers Li^+^ can not only stabilize biomacromolecules but also enhance the permeability of those constituting cellular membranes, which in turn can enhance the entry of other thermo-/osmo- protective solutes (39, 40).

The compatible solute glycine-betaine also significantly increased the frequency of CFU-retention, as well as the percentage of viable cells, at 70°C, for both SMMA_5 and SMMA_5_TC. Furthermore, at 50°C, glycine-betaine fortification of MST brought about an unprecedented CFU-growth for SMMA_5, and a small but definite increase in the existing CFU-growth of SMMA_5_TC. Glycine-betaine is taken-up/synthesized in large amounts by halophilic/halotolerant microbes as their secondary response to high external salt concentrations (28). However, besides acting as osmotic balancers, compatible solutes can also impart a general stabilizing effect on biomacromolecules (41, 42). As the highly-soluble glycine-betaine molecules accumulate within the cell, their preferential exclusion from the immediate hydration sphere of proteins leads to a non-homogeneous distribution within the cell-water, thereby causing a thermodynamic disequilibrium (43). This disequilibrium is minimized via reduction in the volume of cell-water from which the solute is excluded; this, in turn, is achieved via reduction in the surface-area or volume of the proteins through increases in sub-unit assembly and stabilization of secondarily-folded tertiary structures (43). Such biophysical maneuvers at the molecular level can add-up at the cellular level to confer tolerance against any physicochemical stressor that tends to disrupt the system by increasing the disorder within biomacromolecules (43, 44).

Supplementing MST with boron also led to a small but definite increase in the frequency of CFU-retention, as well as the percentage of viable cells, at 70°C, for both SMMA_5 and SMMA_5_TC. At 50°C, boron addition to MST marginally increased the frequency of CFU-retention and CFU-growth for SMMA_5 and SMMA_5_TC respectively. Boron possesses unique bonding properties that can expand the biophysical functions of macromolecules including their modes of Lewis acidity. *In vitro* insertion of boron to proteins in the form of boronoalanine has been shown to allow dative bond-mediated and site-dependent protein Lewis acid–base-pairing, which in turn can generate new functions, including stability in the face of high temperature or proteolytic attack (45). Furthermore, it is theoretically not impossible that boron bonds with membrane fatty acids to form organoborane complexes which in turn provide cross-linkages, and thereby stability, for structures such as cellular membranes.

### Oligotrophy as a strategy for survival at high temperature

After 12 h incubation at 37°C, cellular growth yield of both SMMA_5 and SMMA_5_TC was higher in 1X R2A than in any of the dilution grades of the medium. But, after 12 h incubation at 50°C, as well as after 45 minutes at 70°C, all the dilution grades of R2A supported the retention of higher CFU percentages than undiluted R2A, for both SMMA_5 and SMMA_5_TC. When compared with the corresponding MST phenotype, CFU-retention in the different dilution grades of R2A was

- higher for SMMA_5, but not SMMA_5_TC, when incubation was carried out for 12 h at 50°C, and
- higher for both SMMA_5 and SMMA_5_TC when incubation was carried out for 45 minutes at 70°C.

Thus, SMMA_5 and SMMA_5_TC were found to be best adapted to cope 50°C temperature under oligotrophic and chemolithoautotrophic conditions respectively, whereas both strains were best adapted to cope 70°C under oligotrophic condition. These findings, in concurrence with the different effects exhibited by the environmental solutes on thermal endurance, indicated that the hot spring *Paracoccus* potentially modulates its metabolic response to thermal stress with increasing temperature, and such response regulation potentially involves a differential control of the global transcriptome, orchestrated over prolonged thermal conditioning. Overall, the survival strategy amid high heat appears to be aimed at slowing down growth, and prioritizing cell-system maintenance over proliferation.

Oligotrophy also affords collateral advantages in combating a wide variety of stresses including high heat (46). It entails a constant ability to take up whatever little nutrients are available in the cells’ chemical milieu by using low-specificity, high-affinity, and low-activation-energy, transport systems (47). However, constant nutrient uptake via low-specificity transport systems implies that hydrogen peroxide (H_2_O_2_) and other reactive oxygen species (ROS) abundant in hydrothermal environments also get unrestricted entry into the cell, as do the potentially thermoprotective inorganic solutes and small organic molecules. Furthermore, cells develop intrinsic resistance against H_2_O_2_/ROS toxicity via enhanced expression of genes responsible for membrane detoxification, and protein repair and maintenance (48), which in turn protect the cells’ macromolecules against the destabilizing effects of heat.

### Genomic underpinnings of thermal endurance

The hot spring *Paracoccus* possessed 335 unique genes, plus a large repertoire of genes involved in protein/DNA quality management under stress conditions. While many of its genes have potential roles in thermal adaptation, the genome as a whole is enriched in genes for cell wall/membrane/envelope biogenesis, and transport and metabolism of amino acids and lipids. Thermal conditioning, or the lack of it, appears to have a small but definite effect on the genome. This is consistent with the fact that temperature influences the rate and nature of mutations, and thereby natural selection, evolution, and biogeography of a microorganism (49). Of the total 43 SNPs identified across the aligned portions of the SMMA_5_TC and SMMA_5 genomes, 18 were in protein-coding genes. Out of these 18, again, 13 and five were nsSNPs and sSNPs respectively: a ratio that is potentially indicative of adaptive evolution (50) or relaxed selective constraints (51) in the thermally conditioned (SMMA_5_TC) lineage. Of the 13 nsSNPs detected in protein coding genes, nine and four involved transition and transversion mutations respectively, while seven resulted in radical amino acid replacements having potentials to change protein structures and functions in such ways that can impact activities/phenotypes (Table S7). While radical amino acid replacements often turn out to be beneficial mutations that enhance protein functions (52), those encountered in the putative selenide, water dikinase protein SelD, sulfate adenylyltransferase subunit 2 CysD, and DNA polymerase III subunit gamma/tau, appear to hold major implications for the enhanced thermal endurance or moderate thermophilicity of SMMA_5_TC.

Bacteria have a large repertoire of selenocysteine-containing proteins, and both SelD and CysD play vital roles in their biosynthesis. While the role of selenoproteins in cellular stress management is well documented (53), CysD is also involved in the first step of sulfite formation during the assimilatory reduction of sulfate to sulfide (54). Since ROS is a major driver of cell death during thermal stress (55, 56), the highly reduced sulfide species produced during sulfate reduction can potentially defend microbial cells amid high heat (57). The efficiency of these processes may well be enhanced by potentially modified homologs of SelD and CysD.

Gamma and tau subunits of DNA polymerase III, the stoichiometric components of the replicative complex, are engaged in loading the processivity clamp beta (58, 59). As hyperthermia inhibits DNA replication by either slowing down or arresting the replication forks depending on the temperature level and cell type (60), it is not unlikely that the DNA polymerase III subunit gamma/tau of SMMA_5_TC has potential structural advantages for high-temperature functioning.

### The putative proteome of hot spring *Paracoccus* is biophysically adapted to the environment

With the elevation of temperature and/or increase in the time of heat exposure, conformational changes in the structure of proteins decrease their solubility and tend to precipitate them out of the solution. Solubility of most proteins increase up to the 40- 50°C temperature range, and start decreasing irreversibly beyond these levels, depending on the pH of the medium, and also its ionic strength (61). In this scenario, higher electrostatic repulsion between the molecules of a protein, compared with the degree of hydrophobic interactions between them, can promote solubility via enhanced interaction with the solvent. At pH values above their pI, net charge of proteins become negative, so repulsion between their molecules, and thereby their solubility (interactions with water), increases (61). The average pI values predicted for all the putative proteins of SMMA_5 and SMMA_5_TC were 6.86 and 6.87 respectively, with 61.2% and 61.1% of all proteins of the two strains having pI values <6.86 and <6.87 respectively.

Furthermore, in oligotrophic media, pH 7.5 was found to be optimum for growth at 37°C, and CFU-retention at 50°C and 70°C, concurrent to which, 65.8% and 65.7% of all SMMA_5 and SMMA_5_TC proteins were predicted to have pI <7.5 respectively. In this context it is further noteworthy that the measured pH of Lotus Pond’s vent-water from where SMMA_5 was isolated ranged between 7.2 and 8.0, corresponding to *in situ* temperatures 85°C and 78°C (7). In this way, both *in vitro* pH optimum and the *in situ* pH of the habitat were above the average pI predicted for the total proteomes of SMMA_5 and SMMA_5_TC. Moreover, these *in vitro* or *in situ* pH values, at high temperatures, represent considerably alkaline environments, which in the light of the above biophysical considerations appear to be favorable for the conformational integrity, and the solubility, of the majority of proteins of this microorganism.

### Metabolic deceleration as central to thermal endurance

MST incubation at 50°C and 70°C, with or without lithium, boron or glycine-betaine, showed that a greater proportion of cells always remain metabolically active and viable, compared to what percentage of cells remain ready to divide and form colonies upon withdrawal of heat. Corroboratively, though one or two generations of CFU-growth was achieved over a period of 12 h at 50°C in MST, rapid cell division at a rate lower than the strain’s normal generation time (in MST at 37°C, generation time for both SMMA_5 and SMMA_5_TC was ∼90 minutes) was deleterious to the survival of the entire population at 70°C. All these were reflective of the fact that an overall deceleration of metabolism, underplaying cell division and aimed at achieving a cost-effective maintenance of the cell-system, was central to the thermal endurance strategy of the hot spring *Paracoccus*. Development of “viable but not readily culturable” states in high proportions under different stress conditions is a widespread phenomenon in bacteria (62, 63). At the level of the genome, metabolic deceleration could involve intricate regulation of loci attributed to growth and cell division, oxidative stress management, quality control of biomacromolecules, maintenance of cell wall/envelope integrity, membrane permeability, transport, and energy production/utilization. Concomitantly, thermal conditioning, whether *in situ* or *in vitro*, apparently helps memorize the regulatory states of global gene expression which are central to metabolic slow-down and cell-system maintenance. Eventually, at the level of the biomacromolecules, mitigation of the disordering effects of heat requires such intrinsic entropy-minimizing mechanisms (1, 64) which can accomplish a fine balance between structural integrity (needed to avoid denaturation) and flexibility (necessary to preserve functionality): while the hot spring *Paracoccus* ought to possess native biophysical contrivances to stabilize its macromolecular structures amid high heat (and thereby lower the energy cost of cell-system maintenance), environmental solutes, on their part, appear to augment the indigenous stability-conferring mechanisms. Future studies of transcriptomics, proteomics, and metabolomics, resolved along the dimensions of time, temperature and biogeochemical variables are needed to fully conclude how hot spring mesophiles survive high temperatures over time and space.

## MATERIALS AND METHODS

### Bacterial strains, media and culture conditions

The facultative chemolithoautotroph SMMA_5 was isolated, as described previously, via enrichment in MST medium that essentially contained a modified basal and mineral salts (MS) solution supplemented with thiosulfate (7, 12). The strain was deposited to the Microbial Type Culture Collection and Gene Bank (MTCC), Chandigarh, India with the public accession number MTCC 12601. Since its isolation SMMA_5 was maintained in R2A medium by growing the culture at 37°C and storing at 4°C, with a standard transfer interval of 28 days. From the main culture, the thermally conditioned variant SMMA_5_TC was created and maintained in the following way. A freshly inoculated R2A broth culture of SMMA_5 was first incubated at 50°C for 12 h and then shifted to 37°C for another 12 h. While no turbidity appeared in the culture flask over the 50°C incubation phase, growth corresponding to an OD_600_ of ∼0.6 was obtained at the end of 12 h at 37°C. The culture flask was stored at 25°C and transferred following the above procedure after an interval of 14 days (Fig. 4A). The comparator strains *P*. *pantotrophus* LMG 4218 and *E*. *coli* K-12 were maintained in Luria Broth by growing the cultures at 37°C and storing at 4°C, with a standard transfer interval of 28 days.

### Phylogenetic identification based on 16S rRNA or 92 conserved marker gene sequences

The 16S rRNA gene of the new isolate SMMA_5 was PCR-amplified and sequenced (GenBank accession number LN869532) using the *Bacteria*-specific universal primers 27f and 1492r (65). Pair-wise evolutionary distances between SMMA_5 and other *Paracoccus* species were determined based on their aligned 16S rRNA gene sequences. Subsequent to this, a neighbor-joining tree was constructed following majority rule and strict consensus out of 1000 phylogenetic trees, using the software package MEGAX (66). To check the robustness of the tree’s branching pattern, bootstrap values were calculated based on 1000 replicates.

After the *de novo* sequencing and assembly of their genomes (methods given below), the phylogeny of SMMA_5 and SMMA_5_TC was reconstructed based on 92 conserved marker genes, in relation to the other species of *Paracoccus* for which whole genome sequence were available in the GenBank database (a total of 44 non-redundant genomes for different *Paracoccus* species were included in the analysis on a “one genome per species” basis; two more genomes belonging to *Paracoccus* species were also there in the database but they were not considered due to their low levels of completeness; Table S1). Majority of the genes considered for phylogeny reconstruction (i.e., 67 out of 92), belonged to that functional category for clusters of orthologous groups (COGs) which involves translation, including ribosome structure and biogenesis.

The up-to-date bacterial core gene (UBCG) set was used to identify and align these marker genes (67). Final tree construction from the aligned marker genes was created using RAxML version 8 (68). Final visualization of the resultant tree was done using interactive Tree of Life (iTOL) version 4 (69).

### Growth experiments

Growth or decline in the CFU-count of *Paracoccus* cultures at different temperatures (37-70°C) was first tested in MST medium (pH 7.0) containing 20 mM thiosulfate as the sole source of energy and electrons (70). Subsequently, SMMA_5 and SMMA_5_TC were tested at different temperatures in Luria broth, and then in different concentrations of R2A (diluted up to 1000 times, thereby affording an extremely low absolute concentration of organic carbon) that in turn were adjusted to different levels of pH (7.0- 9.0). Corresponding experiments for *E*. *coli* K-12 were carried out in MS supplemented with 4 g L^-1^ dextrose [MSD, pH 7.0 (70)]. Growth or decline in the CFU-count of SMMA_5 and SMMA_5_TC at different temperatures was further tested in MST supplemented with 4 mM Na_2_B_4_O_7_.10H_2_O (i.e., 16 mM B), 1 mM LiOH.H_2_O (i.e., 1 mM Li) and/or 10 mM glycine-betaine or N,N,N trimethylglycine (C_5_H_11_NO_2_); the three media types were referred to as MSTB, MSTL and MSTG respectively. Notably, the boron and lithium concentrations used in the culture media were close to those recorded in the habitat of SMMA_5 (7, 14).

To test the growth or decline in the CFU-count of a given strain under a particular condition, a seed culture was first prepared in the strain’s designated maintenance medium. Inoculum from the log phase seed culture was transferred to the experimental broth, and incubated at the temperature specified for the experiment. To determine the number of CFUs present mL^-1^ of an experimental culture at a given time-point of incubation (including the 0 h), its various dilution grades [created using 0.9% (w/v) NaCl] were plated in triplicates on to R2A (for SMMA_5 and SMMA_5_TC) or Luria (for LMG 4218 and K-12) agar, and single colonies counted in each of them after 36 h incubation at 37°C. Colony-counts in the different dilution-plates were multiplied by their respective dilution factors, then summed-up across the plates, and finally averaged to get the number of CFUs that were present mL^-1^ of the experimental culture. Cellular growth yield after a given time-period of incubation was expressed as what percentage of the 0 h CFU-count was present in the culture after that period of incubation.

### Testing the effect of thermal-conditioning cycles on growth or CFU-retention

SMMA_5 was subjected to one or five thermal-conditioning cycles (Fig. 4B), both of which started with 1% inoculum transfer to fresh R2A broth, but subsequently involved one or five iteration(s) of the following incubation cycle respectively: 12 h at 50°C and then 12 h at 37°C. In the case of five iterations, two consecutive incubation cycles were separated by an interval of 48 h at 25°C. In either case (whether thermal-conditioning cycles were one or five), cells from the last 37°C-grown culture were dilution plated on R2A agar, and six discrete colonies were selected as six individual sibling strains of SMMA_5. Finally, each member of the two strain-sets - designated as C1_S1 to C1_S6 (Fig. 5A, B, E and F) and C5_S1 to C5_S6 (Fig. 5C, D, G and H), according as they originated from one or five conditioning cycles - was tested individually for its CFU- growth/decline at different temperatures following the procedure already described above.

### Determining the percentage of viable cells by flow cytometry

CFU-count gave the estimate of what percentage of cells in a culture at a given time- point of incubation retained the ability to divide readily, i.e. to form colonies in R2A/Luria agar plates after 36 h at 37°C. On the other hand, what percentage of the cell population remained viable or metabolically active was tested by checking the cells’ ability to accumulate the non-toxic and non-fluorescent molecule fluorescein diacetate and hydrolyze it to fluorescein, which in turn was detected via flow cytometry (71). Post incubation, cells were precipitated from a 100 mL volume of the test culture by centrifugation at 6000 *g* for 20 minutes at 4°C. The supernatant was discarded and the cell pellet resuspended in 2 mL 0.9% NaCl solution. 4 µl of FDA (Sigma, USA) solution (5 mg FDA mL^-1^ dimethyl sulfoxide) was added to the resuspended solution following which the cells were incubated for 15-20 minutes at 37°C. Incubated cells were washed and resuspended again in 0.9% NaCl solution (final resuspension was done in 500 µl) and analyzed using a FACSVerse flow cytometer (Becton Dickinson, USA).

Fluorescence was measured through the excitation wavelengths 475-495 nm and the emission wavelengths 520-530 nm. 10000 randomly-taken cells were analyzed for each sample and dot plots were generated using the fluorescence level of each cell as a function of its forward scattering of a 488 nm wave detected by a photodiode array detector. The data were analyzed using the software BD FACSuite (Becton Dickinson) by specific quadrant gating for each experiment that in turn were predetermined based on unstained samples. In all the individual gating experiments, <2% cells within the FDA-less controls exhibited fluorescence: this was the maximum proportion at which false positives could be there within the different values reported for viable cell percentage.

### Determining glycine-betaine concentration in spent medium

After incubation in MST broth for 12 h at 50°C, SMMA_5 or SMMA_5_TC cells were harvested from 100 mL spent medium by centrifugation at 6000 *g* for 20 minutes. The supernatant was lyophilized down to 1 mL and then filtered by passing through a 0.22 µm cellulose acetate filter (Sartorius Stedim Biotech, Germany). 25 µL of the filtrate was analyzed by high performance liquid chromatography (HPLC) followed by standard UV detection (72), using a Waters platform that encompassed a FlexInject sample-injector, 1525 Binary HPLC pump, a 2998 photodiode array detector, and the software Breeze 2.0 (Waters Corporation, USA). Isocratic elution was carried out in a Hypersil SCX column having 5 µm particle size, 250 mm length and 4.6 mm inner diameter (Phenomenex, USA), using a mixture of disodium phosphate buffer (0.05 M, pH 4.7) and methanol (95:5%, v/v) as the mobile phase (injection volume: 25 µL; flow rate: 1 mL minutes ^-1^ at 25°C; detection wavelength: 195 nm). 0, 2, 4, 8 and 10 mM glycine-betaine standards (Sigma, USA) were used to generate the calibration curve. Glycine-betaine in the sample was identified by comparing its retention time with that of the standard; purity of chromatographic peaks was checked using Breeze 2.0. Quantification was based on the area of the chromatographic peak using the calibration curve Y = mX + C, where Y is the peak-area; X is the concentration of glycine-betaine; m is the slope of the calibration curve; C is the curve intercept for the sample.

### Analysis of genomes and putative proteomes

The whole genome shotgun sequences of SMMA_5 and SMMA_5_TC were determined using an Ion S5 system (Thermo Fisher Scientific, USA) as described previously (73, 74). The assembled genome sequences were annotated for potential ORFs, or CDSs, using the Prokaryotic Genome Annotation Pipeline of the National Center for Biotechnology Information, USA. Completeness of the new genomes, or that of the comparator *Paracoccus* species, was determined using CheckM 1.0.12 on the basis of what percentage of the genus-specific marker genes was present in the genome considered (75). Pairwise DNA-DNA hybridization experiments were carried out using the online software Genome-to-Genome Distance Calculator 3.0 (76). Orthologous genes-based average nucleotide identity between genome pairs was calculated using ChunLab’s online ANI Calculator (77).

To identify potential modifications in the genome of SMMA_5_TC with respect to the SMMA_5 genome, the two sequences were first aligned using the software MAFFT (78) with default parameters. The software SNP-sites (79) was then used to identify single nucleotide polymorphisms (SNPs) from the alignment; in order to enumerate the different transition and transversion events, the VCF file generated was processed with the help of VCFtools (80), while the software SnpEff (81) was used to annotate all the SNPs at the nucleotide as well as translated amino acid sequence levels.

The core, and species-specific unique, genes of *Paracoccus* members including SMMA_5 were delineated with the help of the bacterial pan genome analysis pipeline BPGA 1.3 (82) as described previously (14). This analysis involved the SMMA_5 genome plus the 44 non-redundant genome sequences of *Paracoccus* species (each having a completeness level of ≥97%) available in the GenBank (Table S1). In the working dataset, one genome sequence was included per species, preferably that of the type strain, unless the same was unavailable in the GenBank. SMMA_5_TC was left out because two sibling genomes with identical gene-contents would have distorted the results pertaining to the identification of unique genes.

Putative protein sequence catalogs were derived from the genomes of SMMA_5, SMMA_5_TC and the 44 comparator species of *Paracoccus* which have near-complete genome sequences available in the GenBank. Subsequent to this, the pI profile of each catalog was determined using the software Isoelectric Point Calculator that utilizes the Henderson-Hasselbach equation and calculates pKa values for all the individual protein sequences present in a catalog (83).

For SMMA_5 and 14 other *Paracoccus* species occupying the same clade in the phylogenomic tree (Fig. 1), complete gene catalogs were downloaded from GenBank. Within each catalog, genes attributable to COGs of proteins (84) were selected, and then COG-counts under individual functional categories were determined. Whether COG-count under a functional category was significantly high or low in a particular genome was determined by Chi Square test, as described previously (14), using the contingency chart shown in Table S8 (a *P* value less than 0.001 was considered as the cut-off for inferring whether the presence of COGs under a category was high or low for a species). Hierarchical clustering was carried out to quantitatively decipher the relatedness between the genomes in terms of their enrichment of various COG categories (85). The heat map illustrating the results of hierarchical cluster analysis was constructed with the help of an R script using complete linkage method (14).

## Supporting information

Supplemental Tables, Figures and References

Supplemental Tables that are larger than one A4 size page in width

## DATA AVAILABILITY

The assembled whole genome sequences of SMMA_5 and SMMA_5_TC were deposited to GenBank with the accession numbers WIAB00000000 and WIAC00000000 respectively. The two raw read datasets were deposited to the Sequence Read Archive of the National Center for Biotechnology Information, USA, under the common BioProject PRJNA296849, with the run accession numbers SRR10281900 and SRR10281899 respectively.

## SUPPLEMENTAL MATERIAL

Supplemental materials in the form of a PDF file named “Supplementary_Information.pdf”, and an Excel file named “Supplementary_Dataset.xlsx”, accompany this paper.

## ACKNOWLEDGEMENTS

This research was financed by Bose Institute (via intra-mural faculty grants) as well as the Science and Engineering Research Board (SERB), Government of India (GoI) (SERB grant number was EMR/2016/002703). N.M. received fellowships from SERB and Bose Institute. C.R. got a fellowship from the University Grants Commission, GoI. S.C. received a fellowship from the Department of Biotechnology, GoI. J.S. and S.D. obtained there fellowships from Council of Scientific and Industrial Research, GoI. S.B. received a fellowship from Bose Institute.

## AUTHOR STATEMENT

WG conceived the research program, designed the experiments, interpreted the results, and wrote the paper. NM planned and performed the experiments, harnessed all the modules of the study, and analyzed and curated the data. CR led the studies of genomics and also carried out experiments of physiology. SC, SD and SB carried out the physiological studies while JS performed studies of genomics. RC made critical intellectual contributions to the overall work as well as the paper. All the authors read and vetted the paper. The authors have no conflict of interest to declare.

## REFERENCES

1. Vieille C, Zeikus GJ. 2001. Hyperthermophilic Enzymes: sources, uses, and molecular mechanisms for thermostability. Microbiol Mol Biol Rev 65:1–43. https://doi.org/10.1128/MMBR.65.1.1-43.2001

2. Berezovsky IN, Shakhnovich EI. 2005. Physics and evolution of thermophilic adaptation. Proc Natl Acad Sci U S A 102:12742–12747. https://doi.org/10.1073/pnas.0503890102

3. Shih T.W, Pan T.M. 2011. Stress responses of thermophilic *Geobacillus* sp. NTU 03 caused by heat and heat-induced stress. Microbiol Res 166:346–359. https://doi.org/10.1016/j.micres.2010.08.001

4. Goh KM, Gan HM, Chan K-G, Chan GF, Shahar S, Chong CS, Kahar UM, Chai KP. 2014. Analysis of *Anoxybacillus* genomes from the aspects of lifestyle adaptations, prophage diversity, and carbohydrate metabolism. PLoS One 9:e90549. https://doi.org/10.1371/journal.pone.0090549

5. Power JF, Carere CR, Lee CK, Wakerley GLJ, Evans DW, Button M, White D, Climo MD, Hinze AM, Morgan XC, McDonald IR, Cary SC, Stott MB. 2018. Microbial biogeography of 925 geothermal springs in New Zealand. Nat Commun 9:2876. https://doi.org/10.1038/s41467-018-05020-y

6. Colman DR, Lindsay MR, Boyd ES. 2019. Mixing of meteoric and geothermal fluids supports hyperdiverse chemosynthetic hydrothermal communities. Nat Commun 10:681. https://doi.org/10.1038/s41467-019-08499-1

7. Roy C, Mondal N, Peketi A, Fernandes S, Mapder T, Volvoikar SP, Haldar PK, Nandi N, Bhattacharya T, Mazumdar A, Chakraborty R, Ghosh W. 2020a. Geomicrobial dynamics of Trans-Himalayan sulfur–borax spring system reveals mesophilic bacteria’s resilience to high heat. J Earth Syst Sci 129:157. https://doi.org/10.1007/s12040-020-01423-y

8. Wemheuer B, Taube R, Akyol P, Wemheuer F, Daniel R. 2013. Microbial diversity and biochemical potential encoded by thermal spring metagenomes derived from the Kamchatka Peninsula. Archaea 2013:136714. https://doi.org/10.1155/2013/136714

9. Chan CS, Chan K-G, Tay Y-L, Chua Y-H, Goh KM. 2015. Diversity of thermophiles in a Malaysian hot spring determined using 16S rRNA and shotgun metagenome sequencing. Front Microbiol 6:177. https://doi.org/10.3389/fmicb.2015.00177

10. Ghosh W, Roy C, Roy R, Nilawe P, Mukherjee A, Haldar PK, Chauhan NK, Bhattacharya S, Agarwal A, George A, Pyne P, Mandal S, Rameez MJ, Bala G. 2015. Resilience and receptivity worked in tandem to sustain a geothermal mat community amidst erratic environmental conditions. Sci Rep 5:12179. https://doi.org/10.1038/srep12179

11. Menzel P, Gudbergsdóttir SR, Rike AG, Lin L, Zhang Q, Contursi P, Moracci M, Kristjansson JK, Bolduc B, Gavrilov S, Ravin N, Mardanov A, Bonch-Osmolovskaya E, Young M, Krogh A, Peng X. 2015. Comparative metagenomics of eight geographically remote terrestrial hot springs. Microb Ecol 70:411–424. https://doi.org/10.1007/s00248-015-0576-9

12. Roy C, Alam M, Mandal S, Haldar PK, Bhattacharya S, Mukherjee T, Roy R, Rameez MJ, Misra AK, Chakraborty R, Nanda AK, Mukhopadhyay SK, Ghosh W. 2016. Global association between thermophilicity and vancomycin susceptibility in Bacteria. Front Microbiol 7:412. https://doi.org/10.3389/fmicb.2016.00412

13. Roy C, Bakshi U, Rameez MJ, Mandal S, Haldar PK, Pyne P, Ghosh W. 2019. Phylogenomics of an uncultivated, aerobic and thermophilic, photoheterotrophic member of Chlorobia sheds light into the evolution of the phylum Chlorobi. Comput Biol Chem 80:206–216. https://doi.org/10.1016/j.compbiolchem.2019.04.001

14. Roy C, Rameez MJ, Haldar PK, Peketi A, Mondal N, Bakshi U, Mapder T, Pyne P, Fernandes S, Bhattacharya S, Roy R, Mandal S, O’Neill WK, Mazumdar A, Mukhopadhyay SK, Mukherjee A, Chakraborty R, Hallsworth JE, Ghosh W. 2020b. Microbiome and ecology of a hot spring-microbialite system on the Trans-Himalayan Plateau. Sci Rep 10:5917. https://doi.org/10.1038/s41598-020-62797-z

15. Mondal N, Peketi A, Mapder T, Roy C, Mazumdar A, Chakraborty R, Ghosh W. 2022. Indus and Nubra Valley hot springs affirm the geomicrobiological specialties of Trans-Himalayan hydrothermal systems. J Earth Syst Sci 131:12. https://doi.org/10.1007/s12040-021-01757-1

16. Baker GC, Gaffar S, Cowan DA, Suharto AR. 2001. Bacterial community analysis of Indonesian hot springs. FEMS Microbiol Lett 200:103–109. https://doi.org/10.1111/j.1574-6968.2001.tb10700.x

17. Ghilamicael AM, Budambula NLM, Anami SE, Mehari T, Boga HI. 2017. Evaluation of prokaryotic diversity of five hot springs in Eritrea. BMC Microbiol 17:203. https://doi.org/10.1186/s12866-017-1113-4

18. Mendonca AF, Knabel SJ. 1994. A novel strictly anaerobic recovery and enrichment system incorporating lithium for detection of heat-injured *Listeria monocytogenes* in pasteurized milk containing background microflora. Appl Environ Microbiol 60:4001– 4008. https://doi.org/10.1128/aem.60.11.4001-4008.1994

19. Rudolph B, Gebendorfer KM, Buchner J, Winter J. 2010. Evolution of *Escherichia coli* for growth at high temperatures. J Biol Chem 285:19029–19034. https://doi.org/10.1074/jbc.M110.103374

20. Anggarini S, Murata M, Kido K, Kosaka T, Sootsuwan K, Thanonkeo P, Yamada M. 2020. Improvement of thermotolerance of *Zymomonas mobilis* by genes for reactive oxygen species-scavenging enzymes and heat shock proteins. Front Microbiol 10:3073. https://doi.org/10.3389/fmicb.2019.03073

21. Matsushita K, Azuma Y, Kosaka T, Yakushi T, Hoshida H, Akada R, Yamada M. 2016. Genomic analyses of thermotolerant microorganisms used for high-temperature fermentations. Biosci Biotechnol Biochem 80:655–668. https://doi.org/10.1080/09168451.2015.1104235

22. Kosaka T, Nakajima Y, Ishii A, Yamashita M, Yoshida S, Murata M, Kato K, Shiromaru Y, Kato S, Kanasaki Y, Yoshikawa H, Matsutani M, Thanonkeo P, Yamada M. 2019. Capacity for survival in global warming: Adaptation of mesophiles to the temperature upper limit. PLoS One 14:e0215614. https://doi.org/10.1371/journal.pone.0215614

23. Ghosh W, Mallick S, Haldar PK, Pal B, Maikap SC and Gupta SKD. 2012. Molecular and cellular fossils of a mat-like microbial community in geothermal boratic sinters. Geomicrobiol J 29 879–885. https://doi.org/10.1080/01490451.2011.635761

24. Gupta RD, Arora S. 2017. Characteristics of the soils of Ladakh region of Jammu and Kashmir. J Soil Water Conserv 16:260. https://doi.org/10.5958/2455-7145.2017.00037.6

25. Cho JC, Giovannoni SJ. 2004. Cultivation and growth characteristics of a diverse group of oligotrophic marine Gammaproteobacteria. Appl Environ Microbiol 70:432– 440. https://doi.org/10.1128/AEM.70.1.432-440.2004

26. Boch J, Kempf B, Schmid R, Bremer E. 1996. Synthesis of the osmoprotectant glycine betaine in *Bacillus subtilis*: characterization of the gbsAB genes. J Bacteriol 178:5121–5129. https://doi.org/10.1128/jb.178.17.5121-5129.1996

27. Rosenstein R, Futter-Bryniok D, Götz F. 1999. The choline-converting Pathway in *Staphylococcus xylosus* C2A: genetic and physiological characterization. J Bacteriol 181:2273–2278. https://doi.org/10.1128/JB.181.7.2273-2278.1999

28. Cánovas D, Vargas C, Kneip S, Morón M-J, Ventosa A, Bremer E, Nieto JJY 2000. Genes for the synthesis of the osmoprotectant glycine betaine from choline in the moderately halophilic bacterium *Halomonas elongata* DSM. Microbiology 146:455– 463. https://doi.org/10.1099/00221287-146-2-455

29. Sleator RD, Hill C. 2002. Bacterial osmoadaptation: the role of osmolytes in bacterial stress and virulence. FEMS Microbiol Rev 26:49–71. https://doi.org/10.1111/j.1574-6976.2002.tb00598.x

30. Flahaut S, Frere J, Boutibonnes P, Auffray Y. 1997. Relationship between the thermotolerance and the increase of DnaK and GroEL synthesis in *Enterococcus faecalis* ATCC19433. J Basic Microbiol 37:251–258. https://doi.org/10.1002/jobm.3620370404

31. Casadesús J, D’Ari R. 2002. Memory in bacteria and phage. BioEssays 24:512– 518. https://doi.org/10.1002/bies.10102

32. Govers SK, Mortier J, Adam A, Aertsen A. 2018. Protein aggregates encode epigenetic memory of stressful encounters in individual *Escherichia coli* cells. PLoS Biol 16:e2003853. https://doi.org/10.1371/journal.pbio.2003853

33. Larkin JW, Zhai X, Kikuchi K, Redford SE, Prindle A, Liu J, Greenfield S, Walczak AM, Garcia-Ojalvo J, Mugler A, Süel GM. 2018. Signal percolation within a bacterial community. Cell Syst 7:137–145. https://doi.org/10.1016/j.cels.2018.06.005

34. Lee CK, Anda J de, Baker AE, Bennett RR, Luo Y, Lee EY, Keefe JA, Helali JS, Ma J, Zhao K, Golestanian R, O’Toole GA, Wong GCL. 2018. Multigenerational memory and adaptive adhesion in early bacterial biofilm communities. Proc Natl Acad Sci U S A 115:4471–4476. https://doi.org/10.1073/pnas.1720071115

35. Yang CY, Bialecka-Fornal M, Weatherwax C, Larkin JW, Prindle A, Liu J, Garcia- Ojalvo J, Süel GM. 2020. Encoding membrane-potential-based memory within a microbial community. Cell Syst 10:417–423. https://doi.org/10.1016/j.cels.2020.04.002

36. Bruhn-Olszewska B, Szczepaniak P, Matuszewska E, Kuczyńska-Wiśnik D, Stojowska-Swędrzyńska K, Algara MM, Laskowska E. 2018. Physiologically distinct subpopulations formed in *Escherichia coli* cultures in response to heat shock. Microbiol Res 209:33–42. https://doi.org/10.1016/j.micres.2018.02.002

37. Roark TC, Palacio LA, Gurnev PA, Ray BD, Petrache HI. 2012. Interactions of lithium ions with lipid membranes. Biophys J 102:96a. https://doi.org/10.1016/j.bpj.2011.11.545

38. Jakobsson E, Argüello-Miranda O, Chiu SW, Fazal Z, Kruczek J, Nunez-Corrales S, Pandit S, Pritchet L. 2017. Towards a unified understanding of lithium action in basic biology and its significance for applied biology. J Membr Biol 250:587–604. https://doi.org/10.1007/s00232-017-9998-2

39. Pitzurra M, Szybalski W. 1959. Formation and multiplication of spheroplasts of *Escherichia coli* in the presence of lithium chloride. J Bacteriol 77:614–620. https://doi.org/10.1128/jb.77.5.614-620.1959

40. Hesketh JE, Loudon JB, Reading HW, Glen AIM. 1978. The effect of lithium treatment on erythrocyte membrane ATPase activities and erythrocyte ion content. Br J Clin Pharmacol 5:323–329. https://doi.org/10.1111/j.1365-2125.1978.tb01715.x

41. Colaço C, Sen S, Thangavelu M, Pinder S, Roser B. 1992. Extraordinary stability of enzymes dried in trehalose: simplified molecular biology. Nat Biotechnol 10:1007–1011. https://doi.org/10.1038/nbt0992-1007

42. Qu Y, Bolen CL, Bolen DW. 1998. Osmolyte-driven contraction of a random coil protein. Proc Natl Acad Sci U S A 95:9268–9273. https://doi.org/10.1073/pnas.95.16.9268

43. Welsh DT. 2000. Ecological significance of compatible solute accumulation by micro-organisms: from single cells to global climate. FEMS Microbiol Rev 24:263– 290. https://doi.org/10.1111/j.1574-6976.2000.tb00542.x

44. Lippert K, Galinski EA. 1992. Enzyme stabilization be ectoine-type compatible solutes: protection against heating, freezing and drying. Appl Microbiol Biotechnol 37:61–65. https://doi.org/10.1007/BF00174204

45. Mollner TA, Isenegger PG, Josephson B, Buchanan C, Lercher L, Oehlrich D, Hansen DF, Mohammed S, Baldwin AJ, Gouverneur V, Davis BG. 2021. Post-translational insertion of boron in proteins to probe and modulate function. Nat Chem Biol 17:1245–1261. https://doi.org/10.1038/s41589-021-00883-7

46. Hartke A, Giard JC, Laplace JM, Auffray Y. 1998. Survival of *Enterococcus faecalis* in an oligotrophic microcosm: changes in morphology, development of general stress resistance, and analysis of protein synthesis. Appl Environ Microbiol 64:4238–4245. https://doi.org/10.1128/AEM.64.11.4238-4245.1998

47. 47. Poindexter JS. 1981. Oligotrophy. 63–89. *In* Alexander, M (ed.), Advances in Microbial Ecology. Springer US, Boston, MA. https://doi.org/10.1007/978-1-4615-8306-6_2

48. Meslé MM, Beam JP, Jay ZJ, Bodle B, Bogenschutz E, Inskeep WP. 2017. Hydrogen peroxide cycling in high-temperature acidic geothermal springs and potential implications for oxidative stress response. Front Mar Sci 4:130. https://doi.org/10.3389/fmars.2017.00130

49. Chu X-L, Zhang B-W, Zhang Q-G, Zhu B-R, Lin K, Zhang D-Y. 2018. Temperature responses of mutation rate and mutational spectrum in an *Escherichia coli* strain and the correlation with metabolic rate. BMC Evol Biol 18: 126. https://doi.org/10.1186/s12862-018-1252-8

50. Messier W, Stewart C-B. 1997. Episodic adaptive evolution of primate lysozymes. Nature 385:151–154. https://doi.org/10.1038/385151a0

51. Crandall K, Hillis D. 1997. Rhodopsin evolution in the dark. Nature 387:667–668. https://doi.org/10.1038/42628

52. Dagan T, Talmor Y, Graur D. 2002. Ratios of Radical to conservative amino acid replacement are affected by mutational and compositional factors and may not be indicative of positive Darwinian selection. Mol Biol Evol 19:1022–1025. https://doi.org/10.1093/oxfordjournals.molbev.a004161

53. Cao L, Tang J, Li Q, Xu J, Jia G, Liu G, Chen X, Shang H, Cai J, Zhao H. 2016. Expression of selenoprotein genes is affected by heat stress in IPEC-J2 Cells. Biol Trace Elem Res 172:354–360. https://doi.org/10.1007/s12011-015-0604-0

54. Leyh TS, Vogt TF, Suo Y. 1992. The DNA sequence of the sulfate activation locus from *Escherichia coli* K-12. J Biol Chem 267:10405–10410. https://doi.org/10.1016/S0021-9258(19)50034-5

55. Marcén M, Ruiz V, Serrano MJ, Condón S, Mañas P. 2017. Oxidative stress in *E*. *coli* cells upon exposure to heat treatments. Int J Food Microbiol 241:198–205. https://doi.org/10.1016/j.ijfoodmicro.2016.10.023

56. Hong Y, Zeng J, Wang X, Drlica K, Zhao X. 2019. Post-stress bacterial cell death mediated by reactive oxygen species. Proc Natl Acad Sci U S A 116:10064–10071. https://doi.org/10.1073/pnas.1901730116

57. Hatzios SK, Bertozzi CR. 2011. The regulation of sulfur metabolism in *Mycobacterium tuberculosis*. PLoS Pathog 7:e1002036. https://doi.org/10.1371/journal.ppat.1002036

58. Flower AM, McHenry CS. 1990. The gamma subunit of DNA polymerase III holoenzyme of *Escherichia coli* is produced by ribosomal frameshifting. Proc Natl Acad Sci U S A 87:3713–3717. https://doi.org/10.1073/pnas.87.10.3713

59. Walker JR, Hervas C, Ross JD, Blinkova A, Walbridge MJ, Pumarega EJ, Park MO, Neely HR. 2000. *Escherichia coli* DNA polymerase III tau- and gamma-subunit conserved residues required for activity in vivo and in vitro. J Bacteriol 182:6106– 6113. https://doi.org/10.1128/JB.182.21.6106-6113.2000

60. Velichko AK, Petrova NV, Kantidze OL, Razin SV. 2012. Dual effect of heat shock on DNA replication and genome integrity. Mol Biol Cell 23:3450–3460. https://doi.org/10.1091/mbc.E11-12-1009

61. Zayas JF. 1997. Solubility of Proteins, p. 6–75. In Zayas, JF (ed.), Functionality of Proteins in Food. Springer, Berlin, Heidelberg. https://doi.org/10.1007/978-3-642-59116-7_2

62. Oliver JD. 2005. The viable but nonculturable state in bacteria. J Microbiol 43:93– 100.

63. Li L, Mendis N, Trigui H, Oliver JD, Faucher SP. 2014. The importance of the viable but non-culturable state in human bacterial pathogens. Front Microbiol 5. https://doi.org/10.3389/fmicb.2014.00258

64. Fields PA. 2001. Review: Protein function at thermal extremes: balancing stability and flexibility. Comp Biochem Physiol A Mol Integr Physiol 129:417–431. https://doi.org/10.1016/S1095-6433(00)00359-7

65. Crocetti GR, Hugenholtz P, Bond PL, Schuler A, Keller J, Jenkins D, Blackall LL. 2000. Identification of polyphosphate-accumulating organisms and design of 16S rRNA-directed probes for their detection and quantitation. Appl Environ Microbiol 66:1175–1182. https://doi.org/10.1128/AEM.66.3.1175-1182.2000

66. Kumar S, Stecher G, Li M, Knyaz C, Tamura K. 2018. MEGA X: Molecular evolutionary genetics analysis across computing platforms. Mol Biol Evol 35:1547– 1549. https://doi.org/10.1093/molbev/msy096

67. Na SI, Kim YO, Yoon SH, Ha S, Baek I, Chun J. 2018. UBCG: Up-to-date bacterial core gene set and pipeline for phylogenomic tree reconstruction. J Microbiol 56:280–285. https://doi.org/10.1007/s12275-018-8014-6

68. Stamatakis A. 2014. RAxML version 8: a tool for phylogenetic analysis and post- analysis of large phylogenies. Bioinformatics 30:1312–1313. https://doi.org/10.1093/bioinformatics/btu033

69. Letunic I, Bork P. 2019. Interactive tree of life (iTOL) v4: recent updates and new developments. Nucleic Acids Res 47:W256–W259. https://doi.org/10.1093/nar/gkz239

70. Pyne P, Alam M, Rameez MJ, Mandal S, Sar A, Mondal N, Debnath U, Mathew B, Misra AK, Mandal AK, Ghosh W. 2018. Homologs from sulfur oxidation (Sox) and methanol dehydrogenation (Xox) enzyme systems collaborate to give rise to a novel pathway of chemolithotrophic tetrathionate oxidation. Mol Microbiol 109:169–191. https://doi.org/10.1111/mmi.13972

71. Battin TJ. 1997. Assessment of fluorescein diacetate hydrolysis as a measure of total esterase activity in natural stream sediment biofilms. Sci Total Environ 198:51– 60. https://doi.org/10.1016/S0048-9697(97)05441-7

72. Zamarreño A, Cantera RG, Garcia-Mina JM. 1997. Extraction and determination of glycine betaine in liquid fertilizers. J Agric Food Chem 45:774–776. https://doi.org/10.1021/jf960342h

73. Bhattacharya S, Roy C, Mandal S, Sarkar J, Rameez MJ, Mondal N, Mapder T, Chatterjee S, Pyne P, Alam M, Haldar PK, Roy R, Fernandes S, Peketi A, Chakraborty R, Mazumdar A, Ghosh W. 2020. Aerobic microbial communities in the sediments of a marine oxygen minimum zone. FEMS Microbiol Lett 367:fnaa157. https://doi.org/10.1093/femsle/fnaa157

74. Sen S, Saha T, Bhattacharya S, Nidhi, Mondal N, Ghosh W, Chakraborty R. 2020. Draft genome sequences of two boron-tolerant, arsenic-resistant, gram-positive bacterial strains, *Lysinibacillus* sp. OL1 and *Enterococcus* sp. OL5, Isolated from boron-fortified cauliflower-growing field soils of northern West Bengal, India. Microbiol Resour Announc 9:e01438–19. https://doi.org/10.1128/MRA.01438-19

75. Parks DH, Imelfort M, Skennerton CT, Hugenholtz P, Tyson GW. 2015. CheckM: assessing the quality of microbial genomes recovered from isolates, single cells, and metagenomes. Genome Res 25:1043–1055. https://doi.org/10.1101/gr.186072.114

76. Meier-Kolthoff JP, Carbasse JS, Peinado-Olarte RL, Göker M. 2022. TYGS and LPSN: a database tandem for fast and reliable genome-based classification and nomenclature of prokaryotes. Nucleic Acids Res 50:D801–D807. https://doi.org/10.1093/nar/gkab902

77. Yoon S-H, Ha S, Lim J, Kwon S, Chun J. 2017. A large-scale evaluation of algorithms to calculate average nucleotide identity. Antonie Van Leeuwenhoek 110:1281–1286. https://doi.org/10.1007/s10482-017-0844-4

78. Katoh K, Misawa K, Kuma K, Miyata T. 2002. MAFFT: a novel method for rapid multiple sequence alignment based on fast Fourier transform. Nucleic Acids Res 30:3059–3066. https://doi.org/10.1093/nar/gkf436

79. Page AJ, Taylor B, Delaney AJ, Soares J, Seemann T, Keane JA, Harris SRY 2016. SNP-sites: rapid efficient extraction of SNPs from multi-FASTA alignments. Microb Genomics 2:e000056. https://doi.org/10.1099/mgen.0.000056

80. Danecek P, Auton A, Abecasis G, Albers CA, Banks E, DePristo MA, Handsaker RE, Lunter G, Marth GT, Sherry ST, McVean G, Durbin R, 1000 Genomes Project Analysis Group. 2011. The variant call format and VCFtools. Bioinformatics 27: 2156–2158. https://doi.org/10.1093/bioinformatics/btr330

81. Cingolani P, Platts A, Wang LL, Coon M, Nguyen T, Wang L, Land SJ, Lu X, Ruden DM. 2012. A program for annotating and predicting the effects of single nucleotide polymorphisms, SnpEff: SNPs in the genome of *Drosophila melanogaster* strain w1118; iso-2; iso-3. Fly (Austin) 6:80–92. https://doi.org/10.4161/fly.19695

82. Chaudhari NM, Gupta VK, Dutta C. 2016. BPGA- an ultra-fast pan-genome analysis pipeline. Sci Rep 6:24373. https://doi.org/10.1038/srep24373

83. Kozlowski LP. 2016. IPC – Isoelectric Point Calculator. Biol Direct 11:55. https://doi.org/10.1186/s13062-016-0159-9

84. Tatusov RL, Fedorova ND, Jackson JD, Jacobs AR, Kiryutin B, Koonin EV, Krylov DM, Mazumder R, Mekhedov SL, Nikolskaya AN, Rao BS, Smirnov S, Sverdlov AV, Vasudevan S, Wolf YI, Yin JJ, Natale DA. 2003. The COG database: an updated version includes eukaryotes. BMC Bioinform 4:41. https://doi.org/10.1186/1471-2105-4-41

85. Köhn HF, Hubert LJ. 2015. Hierarchical Cluster Analysis, p. 1–13. In Wiley StatsRef: Statistics Reference Online. John Wiley & Sons, Ltd. https://doi.org/10.1002/9781118445112.stat02449.pub2

