## Supplemental Tables, Figures and References for "Thermal endurance by a hot-spring-dwelling phylogenetic relative of the mesophilic *Paracoccus*"


---

### Contents

#### Supplementary Tables


**Table S1.** The 44 comparator species of *Paracoccus* which have near-complete genome sequences available in the GenBank and so were considered for the present studies of comparative genomics.

**Table S2.** Bioinformatically predicted average isoelectric points determined individually for the putative protein contents of SMMA_5, SMMA_5_TC, and the 44 comparator species of *Paracoccus*.

**Table S3.** The numbers of core, accessory, and unique, genes present in the genomes of SMMA_5, and the 44 comparator species of *Paracoccus*.

**Table S4.** The 335 unique genes of SMMA_5, compared with other *Paracoccus* species, listed alongside the characteristics of the putative proteins encoded. Since this Table is larger than one A4 page in width it has been provided as an Excel sheet named Table S4, within the Excel Workbook named Supplementary_Dataset.

**Table S5.** Chaperones and heat shock proteins encoded by SMMA_5, SMMA_5_TC, *Paracoccus denitrificans*, *Escherichia coli*, and two representative thermophilic taxa that co-inhabited the Lotus Pond hot spring with SMMA_5, according to Roy et al. (2020a).

**Table S6.** Putative gene products encoded from the genomes of SMMA_5 as well as SMMA_5_TC that can potentially play decisive roles in thermal adaptation by means of regulated gene expression.

**Table S7.** Single nucleotide substitutions detected across the completely aligned segments of the SMMA_5 and SMMA_5_TC genomes. Since this Table is larger than one A4 page in width it has been provided as an Excel sheet named Table S7, within the Excel Workbook named Supplementary_Dataset.

**Table S8.** The contingency table that was used to determine by Chi-square test, whether individual COG-counts under different functional categories across *Paracoccus* species including SMMA_5 were significantly high or low. Since this Table is larger than one A4 page in width it has been provided as an Excel sheet named Table S8, within the Excel Workbook named Supplementary_Dataset.

### Supplementary References

Full reference has been given for every paper cited within the Supplementary Materials.

### Supplementary Figures

**Figure S1.** 16S rRNA gene sequence based neighbor-joining tree showing the phylogenetic relationships between the new isolate SMMA_5 and representative strains of existing *Paracoccus* species.

**Figure S2.** Increase or decrease in the CFU-counts of SMMA_5 and SMMA_5_TC, during incubation in MST, MSTB, MSTL and MSTG, at 37°C, 50°C and 70°C.

**Figure S3.** Increase or decrease in the CFU-counts of SMMA_5 and SMMA_5_TC, during incubation in the different dilution grades of R2A, at 37°C, 50°C and 70°C.

**Figure S4.** Increase or decrease in the CFU-counts of SMMA_5 and SMMA_5_TC, during incubation at 37°C, 50°C and 70°C in 1X R2A or 0.001X R2A medium having different pH levels.

**Figure S5.** Final CFU-counts of SMMA_5 and SMMA_5_TC as percentages of the initial levels, after incubation at 37°C, 50°C and 70°C, in 1X R2A or 0.001X R2A medium having different pH levels.

**Figure S6.** Average isoelectric points (pI values) predicted bioinformatically, and individually, for the putative protein contents of SMMA_5, SMMA_5_TC, and the 44 other *Paracoccus* species considered for comparative analysis (see Table S1 for the details of the comparator genomes).

### Supplementary Tables

**Table S1.** The 44 comparator species of *Paracoccus* which have near-complete genome sequences available in the GenBank[1] and so were considered for the present studies of comparative genomics.

| | Name of the organism | Reference sequence ID of the genome | Genome size (Mb) | G+C content (%) | Genome completeness (%) |
|---|---|---|---|---|---|
| 1 | *Paracoccus acridae* CGMCC 1.15419$^T$ | GCF_014642735.1 | 3.99 | 65.3 | 99.09 |
| 2 | *Paracoccus aerius* KCTC 42845$^T$ | GCF_014656455.1 | 4.23 | 64.9 | 99.09 |
| 3 | *Paracoccus aestuarii* DSM 19484$^T$ | GCF_003594815.1 | 3.75 | 67.7 | 98.79 |
| 4 | *Paracoccus aestuariivivens* NBRC 111993$^T$ | GCF_009711225.1 | 4.56 | 61.4 | 99.62 |
| 5 | *Paracoccus alcaliphilus* DSM 8512$^T$ | GCF_900110285.1 | 4.61 | 64.3 | 99.09 |
| 6 | *Paracoccus alkanivorans* 4-2$^T$ | GCF_003697785.1 | 4.66 | 62.1 | 99.70 |
| 7 | *Paracoccus alkenifer* DSM 11593$^T$ | GCF_900108405.1 | 3.19 | 67.3 | 98.28 |
| 8 | *Paracoccus aminophilus* JCM 7686$^T$ | NC_022041.1 | 3.61 | 63.4 | 99.29 |
| 9 | *Paracoccus aminovorans* JCM 7685$^T$ | NZ_LN832559.1 | 3.11 | 67.2 | 98.48 |
| 10 | *Paracoccus chinensis* CGMCC 1.7655$^T$ | GCF_900102885.1 | 3.63 | 68.1 | 98.79 |
| 11 | *Paracoccus contaminans* RKI 16-01929$^T$ | NZ_CP020612.1 | 2.94 | 68.7 | 98.17 |
| 12 | *Paracoccus denitrificans* ASM406373v1 | NZ_CP035090.1, NZ_CP035091.1 | 4.59 | 66.8 | 99.70 |
| 13 | *Paracoccus gahaiensis* KCTC42687$^T$ | GCF_005048225.1 | 4.18 | 67.5 | 99.09 |
| 14 | *Paracoccus haematequi* CIP 111624$^T$ | GCF_900631945.1 | 4.10 | 66.6 | 99.02 |
| 15 | *Paracoccus haeundaensis* CGMCC 1.8012 | NZ_VDDC00000000.1 | 4.10 | 66.2 | 99.39 |
| 16 | *Paracoccus hibiscisoli* CCTCC AB 2016182$^T$ | GCF_005048265.1 | 3.96 | 67.1 | 98.79 |
| 17 | *Paracoccus homiensis* DSM 17862$^T$ | GCF_900111675.1 | 3.87 | 63.8 | 99.39 |
| 18 | *Paracoccus isoporae* DSM 22220$^T$ | GCF_900101865.1 | 3.52 | 65.8 | 98.18 |
| 19 | *Paracoccus jeotgali* CBA4604$^T$ | NZ_CP025583.1 | 3.12 | 66.8 | 97.68 |
| 20 | Paracoccus kondratievae BJQ0001 | NZ_CP045072.1, NZ_CP045073.1 | 3.77 | 62.9 | 99.44 |
| 21 | *Paracoccus laeviglucosivorans* DSM 100094$^T$ | GCF_900182695.1 | 4.30 | 63.1 | 99.62 |
| 22 | *Paracoccus liaowanqingii* 2251$^T$ | NZ_CP038439.1 | 3.27 | 68.2 | 98.42 |

| | | | | | |
|---|---|---|---|---|---|
| 23 | *Paracoccus limosus* JCM 17370$^T$ | GCF_009711185.1 | 3.91 | 66.1 | 99.62 |
| 24 | *Paracoccus litorisediminis* NBRC 112902$^T$ | GCF_009711205.1 | 5.37 | 63.6 | 99.62 |
| 25 | *Paracoccus lutimaris* CECT 8525$^T$ | GCF_003337565.1 | 4.21 | 65.2 | 99.60 |
| 26 | *Paracoccus marcusii* CGMCC 1.8602 | GCF_006151785.1 | 3.67 | 66.9 | 99.39 |
| 27 | *Paracoccus pantotrophus* ACCC10489 | NZ_CP058689.1, NZ_CP058690.1 | 3.99 | 67.3 | 99.39 |
| 28 | *Paracoccus salipaludis* WN007$^T$ | GCF_002287065.1 | 3.67 | 68.6 | 99.44 |
| 29 | *Paracoccus saliphilus* DSM 18447$^T$ | GCF_900156835.1 | 4.57 | 61.1 | 99.39 |
| 30 | *Paracoccus sanguinis* DSM 29303$^T$ | GCF_900106665.1 | 3.59 | 70.7 | 99.90 |
| 31 | *Paracoccus sediminis* DSM 26170$^T$ | GCF_900188295.1 | 3.65 | 66.0 | 99.48 |
| 32 | *Paracoccus seriniphilus* DSM 14827$^T$ | GCF_900199195.1 | 4.20 | 61.6 | 99.09 |
| 33 | *Paracoccus siganidrum* M26$^T$ | GCF_003709565.1 | 4.90 | 67.9 | 99.09 |
| 34 | *Paracoccus solventivorans* DSM 6637$^T$ | GCF_900142875.1 | 3.38 | 68.7 | 98.79 |
| 35 | *Paracoccus sphaerophysae* HAMBI 3106$^T$ | GCF_000763805.1 | 3.36 | 69.1 | 98.74 |
| 36 | *Paracoccus subflavus* GY0581$^T$ | GCF_004310345.1 | 3.18 | 65.6 | 98.69 |
| 37 | *Paracoccus sulfuroxidans* CGMCC 1.5364$^T$ | GCF_007830335.1 | 4.04 | 64.2 | 99.62 |
| 38 | *Paracoccus suum* SC2-6$^T$ | NZ_CP030918.1 | 3.25 | 66.9 | 99.19 |
| 39 | *Paracoccus thiocyanatus* ATCC 700171$^T$ | GCF_900156255.1 | 3.52 | 67.2 | 99.55 |
| 40 | *Paracoccus tibetensis* CGMCC 1.8925$^T$ | GCF_900102505.1 | 3.91 | 68.3 | 99.39 |
| 41 | *Paracoccus versutus* DSM 582$^T$ | GCF_003387045.1 | 5.63 | 67.6 | 99.39 |
| 42 | *Paracoccus yeei* ASM274949v1 | NZ_CP024422.1 | 3.59 | 67.3 | 99.55 |
| 43 | *Paracoccus zeaxanthinifaciens* ATCC 21588$^T$ | NZ_ATUJ00000000.1 | 3.04 | 67.2 | 97.97 |
| 44 | *Paracoccus zhejiangensis* J6$^T$ | NZ_CP025430.1 | 3.99 | 65.8 | 98.99 |

1 Genome sequences are also available in the public databases for *Paracoccus halophilus* CGMCC 1.6117$^T$ (GCF_900111785.1) and *Paracoccus mutanolyticus* RSP02$^T$ (NZ_CP030239.1), but they were not included in the present analyses because of their low levels of completeness (66.67% and 25.23% respectively).

**Table S2.** Bioinformatically predicted average isoelectric points (pI values) determined individually for the putative protein contents of SMMA_5, SMMA_5_TC, and the 44 comparator species of *Paracoccus* which have near-complete genome sequences available in the GenBank.

| | Name of the organism | No. of proteins encoded by the genome | Average pI | SD of average pI |
|---|---|---|---|---|
| 1 | SMMA_5 | 3001 | 6.86 | 2.088327 |
| 2 | SMMA_5_TC | 2988 | 6.87 | 2.093058 |
| 3 | *Paracoccus acridae* CGMCC 1.15419$^T$ | 3886 | 6.56 | 2.023760 |
| 4 | *Paracoccus aerius* KCTC 42845$^T$ | 4125 | 6.61 | 2.043153 |
| 5 | *Paracoccus aestuarii* DSM 19484$^T$ | 3680 | 6.78 | 2.189871 |
| 6 | *Paracoccus aestuariivivens* NBRC 111993$^T$ | 4258 | 6.48 | 1.877906 |
| 7 | *Paracoccus alcaliphilus* DSM 8512$^T$ | 4581 | 6.66 | 2.125421 |
| 8 | *Paracoccus alkanivorans* 4-2$^T$ | 4380 | 6.39 | 1.925626 |
| 9 | *Paracoccus alkenifer* DSM 11593$^T$ | 2997 | 6.64 | 2.089086 |
| 10 | *Paracoccus aminophilus* JCM 7686$^T$ | 3387 | 6.56 | 1.914766 |
| 11 | *Paracoccus aminovorans* JCM 7685$^T$ | 2902 | 6.73 | 2.071890 |
| 12 | *Paracoccus chinensis* CGMCC 1.7655$^T$ | 3545 | 6.68 | 2.112931 |
| 13 | *Paracoccus contaminans* RKI 16-01929$^T$ | 2686 | 6.95 | 2.194853 |
| 14 | *Paracoccus denitrificans* ASM406373v1 | 4378 | 6.68 | 2.030358 |
| 15 | *Paracoccus gahaiensis* KCTC42687$^T$ | 3871 | 6.55 | 2.091142 |
| 16 | *Paracoccus haematequi* CIP 111624$^T$ | 4055 | 6.69 | 2.087728 |
| 17 | *Paracoccus haeundaensis* CGMCC 1.8012 | 3911 | 6.58 | 2.097424 |
| 18 | *Paracoccus hibiscisoli* CCTCC AB 2016182$^T$ | 3711 | 6.67 | 2.128472 |
| 19 | *Paracoccus homiensis* DSM 17862$^T$ | 3840 | 6.46 | 2.002623 |
| 20 | *Paracoccus isoporae* DSM 22220$^T$ | 3355 | 6.34 | 2.032360 |
| 21 | *Paracoccus jeotgali* CBA4604$^T$ | 2853 | 6.45 | 2.022062 |
| 22 | Paracoccus kondratievae BJQ0001 | 2165 | 6.64 | 1.997151 |
| 23 | *Paracoccus laeviglucosivorans* DSM 100094$^T$ | 4233 | 6.57 | 1.976051 |
| 24 | *Paracoccus liaowanqingii* 2251$^T$ | 3021 | 6.56 | 2.114653 |
| 25 | *Paracoccus limosus* JCM 17370$^T$ | 3707 | 6.72 | 2.000091 |
| 26 | *Paracoccus litorisediminis* NBRC 112902$^T$ | 5058 | 6.56 | 1.934091 |

| | | | | |
|---|---|---|---|---|
| 27 | *Paracoccus lutimaris* CECT 8525$^T$ | 3971 | 6.57 | 1.965089 |
| 28 | *Paracoccus marcusii* CGMCC 1.8602 | 3638 | 6.58 | 2.110701 |
| 29 | *Paracoccus pantotrophus* ACCC10489 | 3735 | 6.76 | 2.057470 |
| 30 | *Paracoccus salipaludis* WN007$^T$ | 3444 | 6.70 | 2.110246 |
| 31 | *Paracoccus saliphilus* DSM 18447$^T$ | 4388 | 6.26 | 1.909184 |
| 32 | *Paracoccus sanguinis* DSM 29303$^T$ | 3401 | 6.66 | 2.134532 |
| 33 | *Paracoccus sediminis* DSM 26170$^T$ | 3538 | 6.67 | 2.097230 |
| 34 | *Paracoccus seriniphilus* DSM 14827$^T$ | 3990 | 6.37 | 1.937665 |
| 35 | *Paracoccus siganidrum* M26$^T$ | 4600 | 6.64 | 2.075268 |
| 36 | *Paracoccus solventivorans* DSM 6637$^T$ | 3198 | 6.62 | 2.099228 |
| 37 | *Paracoccus sphaerophysae* HAMBI 3106$^T$ | 2822 | 6.67 | 2.125007 |
| 38 | *Paracoccus subflavus* GY0581$^T$ | 3063 | 6.63 | 2.039024 |
| 39 | *Paracoccus sulfuroxidans* CGMCC 1.5364$^T$ | 3926 | 6.40 | 1.911463 |
| 40 | *Paracoccus suum* SC2-6$^T$ | 3052 | 6.62 | 2.088351 |
| 41 | *Paracoccus thiocyanatus* ATCC 700171$^T$ | 3446 | 6.80 | 2.090078 |
| 42 | *Paracoccus tibetensis* CGMCC 1.8925$^T$ | 3740 | 6.67 | 2.116551 |
| 43 | *Paracoccus versutus* DSM 582$^T$ | 5446 | 6.76 | 2.067002 |
| 44 | *Paracoccus yeei* ASM274949v1 | 3402 | 6.73 | 2.048450 |
| 45 | *Paracoccus zeaxanthinifaciens* ATCC 21588$^T$ | 2893 | 6.36 | 2.024594 |
| 46 | *Paracoccus zhejiangensis* J6$^T$ | 3733 | 6.28 | 1.898157 |

**Table S3.** The numbers of core, accessory, and unique, genes present in the genomes of SMMA_5, and the 44 comparator species of *Paracoccus* which have near-complete genome sequences available in the GenBank.

| | Name of the organism | No. of core genes | No. of accessory genes | No. of unique genes | No. of exclusively absent genes |
|---|---|---|---|---|---|
| 1 | SMMA_5 | 822 | 1742 | 335 | 30 |
| 2 | *Paracoccus acridae* CGMCC 1.15419$^T$ | 822 | 2590 | 299 | 3 |
| 3 | *Paracoccus aerius* KCTC 42845$^T$ | 822 | 2720 | 380 | 3 |
| 4 | *Paracoccus aestuarii* DSM 19484$^T$ | 822 | 2324 | 306 | 2 |
| 5 | *Paracoccus aestuariivivens* NBRC 111993$^T$ | 822 | 2863 | 394 | 0 |
| 6 | *Paracoccus alcaliphilus* DSM 8512$^T$ | 822 | 2806 | 579 | 4 |
| 7 | *Paracoccus alkanivorans* 4-2$^T$ | 822 | 2836 | 611 | 2 |
| 8 | *Paracoccus alkenifer* DSM 11593$^T$ | 822 | 1887 | 196 | 0 |
| 9 | *Paracoccus aminophilus* JCM 7686$^T$ | 822 | 1764 | 679 | 11 |
| 10 | *Paracoccus aminovorans* JCM 7685$^T$ | 822 | 1754 | 246 | 1 |
| 11 | *Paracoccus chinensis* CGMCC 1.7655$^T$ | 822 | 2268 | 286 | 1 |
| 12 | *Paracoccus contaminans* RKI 16-01929$^T$ | 822 | 1528 | 211 | 15 |
| 13 | *Paracoccus denitrificans* ASM406373v1 | 822 | 2917 | 398 | 0 |
| 14 | *Paracoccus gahaiensis* KCTC42687$^T$ | 822 | 2672 | 255 | 1 |
| 15 | *Paracoccus haematequi* CIP 111624$^T$ | 822 | 2671 | 425 | 3 |
| 16 | *Paracoccus haeundaensis* CGMCC 1.8012 | 822 | 2685 | 221 | 1 |
| 17 | *Paracoccus hibiscisoli* CCTCC AB 2016182$^T$ | 822 | 2476 | 243 | 0 |
| 18 | *Paracoccus homiensis* DSM 17862$^T$ | 822 | 2390 | 512 | 5 |
| 19 | *Paracoccus isoporae* DSM 22220$^T$ | 822 | 1880 | 582 | 0 |
| 20 | *Paracoccus jeotgali* CBA4604$^T$ | 822 | 1561 | 407 | 9 |
| 21 | Paracoccus kondratievae BJQ0001 | 822 | 2328 | 302 | 0 |
| 22 | *Paracoccus laeviglucosivorans* DSM 100094$^T$ | 822 | 2641 | 552 | 0 |
| 23 | *Paracoccus liaowanqingii* 2251$^T$ | 822 | 1937 | 204 | 11 |
| 24 | *Paracoccus limosus* JCM 17370$^T$ | 822 | 2418 | 355 | 1 |
| 25 | *Paracoccus litorisediminis* NBRC 112902$^T$ | 822 | 3163 | 817 | 0 |

| | | | | | |
|---|---|---|---|---|---|
| 26 | *Paracoccus lutimaris* CECT 8525$^T$ | 822 | 2467 | 484 | 1 |
| 27 | *Paracoccus marcusii* CGMCC 1.8602 | 822 | 2470 | 159 | 2 |
| 28 | *Paracoccus pantotrophus* ACCC10489 | 822 | 2474 | 282 | 0 |
| 29 | *Paracoccus salipaludis* WN007$^T$ | 822 | 2168 | 245 | 0 |
| 30 | *Paracoccus saliphilus* DSM 18447$^T$ | 822 | 2707 | 686 | 0 |
| 31 | *Paracoccus sanguinis* DSM 29303$^T$ | 822 | 2122 | 345 | 0 |
| 32 | *Paracoccus sediminis* DSM 26170$^T$ | 822 | 2313 | 294 | 0 |
| 33 | *Paracoccus seriniphilus* DSM 14827$^T$ | 822 | 2267 | 744 | 3 |
| 34 | *Paracoccus siganidrum* M26$^T$ | 822 | 3223 | 365 | 0 |
| 35 | *Paracoccus solventivorans* DSM 6637$^T$ | 822 | 1983 | 268 | 0 |
| 36 | *Paracoccus sphaerophysae* HAMBI 3106$^T$ | 822 | 1646 | 228 | 45 |
| 37 | *Paracoccus subflavus* GY0581$^T$ | 822 | 1947 | 224 | 0 |
| 38 | *Paracoccus sulfuroxidans* CGMCC 1.5364$^T$ | 822 | 2370 | 615 | 0 |
| 39 | *Paracoccus suum* SC2-6$^T$ | 822 | 1581 | 555 | 16 |
| 40 | *Paracoccus thiocyanatus* ATCC 700171$^T$ | 822 | 2268 | 213 | 1 |
| 41 | *Paracoccus tibetensis* CGMCC 1.8925$^T$ | 822 | 2293 | 457 | 0 |
| 42 | *Paracoccus versutus* DSM 582$^T$ | 822 | 3451 | 666 | 0 |
| 43 | *Paracoccus yeei* ASM274949v1 | 822 | 2101 | 361 | 12 |
| 44 | *Paracoccus zeaxanthinifaciens* ATCC 21588$^T$ | 822 | 1796 | 231 | 6 |
| 45 | *Paracoccus zhejiangensis* J6$^T$ | 822 | 2268 | 584 | 3 |

**Table S5.** Chaperones and heat shock proteins encoded by SMMA_5, SMMA_5_TC, *Paracoccus denitrificans*, *Escherichia coli*, and two representative thermophilic taxa (*Thermus aquaticus* and *Caldicellulosiruptor saccharolyticus*) that co-inhabited the Lotus Pond hot spring with SMMA_5, according to Roy et al. (2020a).

| **Name of the protein** | **Known function of the protein** | **Number of genes present in the different bacterial genomes (accession numbers have been given) for the protein considered (protein IDs have been given for the homologs present)** | | | | | | **Reference** |
|---|---|---|---|---|---|---|---|---|
| | | ***E. coli* O157:H7str. Sakai (NC_002695.2)** | ***P. denitrificans* PD1222 (NC_008686.1; NC_008687.1)** | **SMMA_5 (WIAB00000000)** | **SMMA_5_TC (WIAC00000000)** | ***T. aquaticus* Y51MC23 (NZ_CP010822.1)** | ***C. saccharolyticus* DSM8903 (NC_009437.1)** | |
| Chaperonin GroEL | High expression levels of GroEL (along with GroES) in bacteria protect proteins at high temperature. | 1 (NP_313151.1) | 1 (WP_011749890.1) | 2 (MSU17203.1, MSU17645.1) | 2 (MST19413.1, MST19880.1) | 1 (WP_003049018.1) | 1 (WP_011916830.1) | Genevaux et al., 2004 |
| Co-chaperone GroES | High expression levels of GroES (along with GroEL) in bacteria protect proteins in high temperature. | 1 (NP_313150.1) | 1 (WP_019353022.1) | 1 (MSU17644.1) | 1 (MST19881.1) | 1 (WP_003049015.1) | 1 (WP_011916829.1) | Genevaux et al., 2004 |
| Heat shock protein Hsp90 | Helps DnaK and GroEL for their functions at the time of thermal stress. Absence of HSP90 delays the recovery of cells from heat shock. | 1 (NP_308553.1) | 0 | 0 | 0 | 0 | 0 | Maleki et al., 2016 |
| Heat shock protein HspQ | Involved in the degradation of certain denatured proteins at the time of heat shock stress. | 1 (NP_309077.2) | 1 (WP_011747158.1) | 1 (MSU15669.1) | 1 (MST18354.1) | 0 | 0 | Shimuta et al., 2004 |
| Heat-inducible transcriptional repressor HrcA | Negative regulator of *grpE-dnaK-dnaJ* and *groELS* operons. Heat | 0 | 1 (WP_011746359.1) | 1 (MSU16083.1) | 1 (MST18385.1) | 0 | 1 (WP_011917271.1) | Hitomi et al., 2003 |

| | | | | | | | | |
|---|---|---|---|---|---|---|---|---|
| | stress causes the dissociation of HrcA from the operon complex, thereby facilitating the transcription of *groE* and *dnaK* operons. | | | | | | | |
| Hsp20 family protein | Chaperones protecting proteins against heat induced denaturation and aggregation. | 2 (NP_308041.1, NP_310905.3) | 4 (WP_011748492.1, WP_011749882.1, WP_011749414.1, WP_011748492.1) | 2 MSU17441.1, MSU17533.1) | 2 (MST19677.1, MST19772.1) | 4 (WP_003047493.1, WP_003045810.1, WP_003049394.1, WP_003049384.1) | 1 (WP_011917576.1) | Li et al., 2012 |
| Hsp33 family molecular chaperone HslO | Redox regulator of molecular chaperones. Protects thermally and oxidatively damaged proteins from irreversible aggregation. | 0 | 0 | 1 (MSU18148.1) | 1 (MST20546.1) | 1 (WP_003045607.1) | 1 (WP_011916464.1) | Barbirz et al., 2000 |
| Hsp70 family protein/ molecular chaperone DnaK | Molecular chaperones that help to avoid the formation of erroneous protein configurations during stressful conditions. | 0 | 2 (WP_011750484.1, WP_011748586.1) | 2 (MSU17549.1, MSU17162.1) | 2 (MST19803.1, MST19983.1) | 1 (WP_060474026.1) | 1 (WP_011917273.1) | Ghazaei, 2017 |
| Molecular chaperone DnaJ/DnaJ domain-containing protein/ chaperone Hsp40 | Prevents aggregation of damaged proteins at the time of hyperosmotic and heat shock. It also stimulates the activity of DnaK. | 2 (NP_308711.1, NP_308042.1) | 2 (WP_011748587.1, WP_011746852.1) | 3 (MSU17161.1, MSU16547.1, MSU16807.1) | 3 (MST19984.1, MST20384.1, MST18153.1) | 1 (WP_003045304.1) | 1 (WP_011917274.1) | Liberek et al., 1991 |

| | | | | | | | | |
|---|---|---|---|---|---|---|---|---|
| Nucleotide exchange factor GrpE/Co-chaperone GrpE | GrpE actively responds to heat shock by preventing the aggregation of denatured proteins. | 1 (NP_311503.1) | 1 (WP_011746358.1) | 1 (MSU16084.1) | 1 (MST18384.1) | 1 (WP_003045305.1) | 1 (WP_011917272.1) | Harrison et al., 1997 |
| RNA polymerase sigma E factor | sigma-E responds to periplasmic protein stress, increased levels of periplasmic lipopolysaccharide after heat shock and oxidative stress and also controls protein processing in the extracytoplasmic compartment. | 1 (NP_311466.1) | 0 | 0 | 0 | 0 | 0 | Raina et al., 1995 |
| RNA polymerase sigma factor RpoH/ RNA polymerase sigma 32 factor RpoH | Heat shock leads to intracellular accumulation of protein, which initiates the transcription of heat shock genes, global transcriptional regulators and genes involved in maintaining membrane functionality and homeostasis. | 1 (NP_312337.1) | 1 (WP_011748503.1) | 1 (MSU17450.1) | 1 (MST19687.1) | 0 | 0 | Straus et al., 1987 |

**Table S6.** Putative gene products encoded from the genomes of SMMA_5 as well as SMMA_5_TC that can potentially play decisive roles in thermal adaptation by means of regulated gene expression.

| | Name of the gene product | Locus tag in SMMA_5 | Locus tag in SMMA_5_TC | Function of the protein | Reference |
|---|---|---|---|---|---|
| 1 | ATP-dependent chaperone ClpB | GB879_14550 | GB880_14620 | ATP-dependent chaperone ClpB that belongs to the Caseinolytic Protease (Clp) family and rescues proteins from an aggregated state, thereby playing decisive roles in thermal adaptation. Expression of ClpB is regulated by heat stress, and its malfunctioning renders bacteria critically sensitive to high temperature. | Allan et al., 1998; Lee et al., 2003 |
| 2 | ATP-dependent Clp protease proteolytic subunit | GB879_01125, GB879_03845 | GB880_03440, GB880_03970 | ATP-dependent caseinolytic protease proteolytic subunit ClpP; this serine protease renders the proteolysis of defective and misfolded proteins of diverse structures and functions, and in doing so plays a key role in bacterial proteostasis alongside the widespread Lon and FtsH proreases. Proteins targeted by ClpP include those involved in biofilm formation, cell-cycle progression and cell division, cell motility, damage repair, heat-stress response, nutrient starvation, stationary phase adaptation, transcription regulation, and various other metabolisms. | Sauer and Baker, 2011; Moreno-Cinos et al., 2019 |
| 3 | ATP-dependent protease | GB879_15030 | GB880_15130 | A member of the LON superfamily of ATP-dependent proteases that catalyze the rapid turnover of short-lived regulatory proteins, as well as damaged, denatured, misfolded or mutant proteins. In doing so LON proteases control radiation resistance, cell division, filamentation, capsular polysaccharide production, and survival under starvation. | Jonas et al., 2013; Pinti et al., 2016 |
| 4 | ATP-dependent zinc metalloprotease FtsH | GB879_04900 | GB880_13585 | ATP-dependent zinc metallopeptidase acts as a processive for cytoplasmic and membrane proteins. It involved in the quality control of integral membrane proteins and also degrades proteins that have not been assembled into complexes such as SecY, $F_0$ATPase subunit a and YccA, and also cytoplasmic proteins sigma-32, LpxC, KdtA and phage lambda cII protein etc. With the help of ATP, it generally dislocates membrane-spanning and periplasmic segments of the protein into the cytoplasm for degradation. It also degrades C-terminal-tagged cytoplasmic proteins which are tagged with an 11-amino-acid nonpolar destabilizing tail via a mechanism involving the 10SA (SsrA) stable RNA. The FtsH regulates LpxC and KdtA expression; so, it is required for synthesis of protein and lipid components in lipopolysaccharide. | Katz and Ron, 2008 |

| | | | | | |
|---|---|---|---|---|---|
| 5 | peptidase | GB879_00935; GB879_01660; GB879_04085; GB879_05795; GB879_07345; GB879_09215; GB879_12600; GB879_14905 | GB880_03250; GB880_03810; GB880_05495; GB880_07355; GB880_07510; GB880_12215; GB880_14670 | The peptidyl-prolyl cis-trans isomerase (rotamase); rotamases function as protein folding chaperones by increasing the rate of protein folding via *cis*-proline / *trans*-proline interconversion in sync with the metabolic condition of the cell. | Weissbach et al., 2002 |
| 6 | peptide-methionine (S)-S-oxide reductase MsrA | GB879_09060 | GB880_07665 | The peptide-methionine (S)-S-oxide reductase MsrA, known to repair oxidation-inactivated proteins by reducing their methionine sulfoxides back to methionine. | Weissbach et al., 2002 |
| 7 | elongation factor Ts | GB879_00375 | GB880_00370 | The elongation factor Ts (EF-Ts) that converts the inactive form of the elongation factor Tu (EF-Tu*GDP) to its active form (EF-Tu*GTP) capable of delivering aminoacyl-tRNAs to the ribosome. In this way, EF-Ts recycles EF-Tu to complete the elongation cycle. | Kudlicki et al., 1997; Wieden et al., 2002 |
| 8 | polyhydroxyalkanoate synthesis repressor PhaR | GB879_03465 | GB880_03080 | Under nutrient-limited and stressed conditions, many microorganisms produce PHAs (linear polyesters produced by bacterial fermentation of sugar or lipids) as intracellular carbon and energy storage materials; PHA granules are separated from the cytoplasm by an amphiphilic layer of small proteins called phasins (PhaP). PhaR acts as a repressor of the *phaP* gene as well as that of *phaR* located downstream of *phaP*. It senses the onset of PHA synthesis as well as the enlargement of PHA granules through direct binding to PHA, thereby allowing PhaP synthesis to proceed in sync with the PHA concentration in the cell. However, free PhaR is apparently incapable of sensing mature PHA granules that are already covered with PhaP and/or other proteins. | Maehara et al., 2002; de Almeida et al., 2007 |
| 9 | Ppx/GppA family phosphatase | GB879_00860 | GB880_12560 | The exopolyphosphate / guanosine pentaphosphate (Ppx/GppA) phosphatase that plays a key role in bacterial survival, starvation-induced stringent response, motility, biofilm formation, sporulation and resistance to complement-mediated killing. While the exopolyphosphatase activity degrades poly-P into a smaller branches of inorganic phosphate (these can serve as energy sources for the synthesis of sugars, nucleosides, and proteins, besides acting as activating precursors for fatty acids, phospholipids, polypeptides, and | Reizer et al., 1993; Srivatsan and Wang, 2008; Malde et al., 2014 |

| | | | | | |
|---|---|---|---|---|---|
| | | | | nucleic acids), the guanosine pentaphosphate (pppGpp) phosphorhydrolase activity generates guanosine tetraphosphate (ppGpp) to regulate stringent response in bacteria. (p)ppGpp, in turn, inhibits RNA synthesis in the event of amino acids scarcity, thereby decreasing translation and conserving amino acids within the bacterial cells; ppGpp triggers growth arrest, and also up-regulates several stress response genes such as those involved in amino acid uptake (from the surrounding media) and biosynthesis. | |
| 10 | SsrA-binding protein SmpB | GB879_13720 | GB880_12560 | The protein SmpB that in association with small stable RNA A (SsrA) molecules earmark incomplete proteins for degradation through cotranslational addition of peptide tags to their C-terminal ends. While such modification of biosynthesis-stalled / biosynthesis-interrupted proteins facilitates their degradation by intracellular proteases, jammed or obstructed ribosomes are also cleared / rescued in the process. | Karzai et al., 2000 |
| 11 | addiction module toxin RelE | GB879_03675 | GB880_02870 | The addiction module toxin RelE, a sequence-specific and ribosome-dependent mRNA endoribonuclease that inhibits translation (and thereby cell growth) during amino acid starvation. | Pandey and Gerdes, 2005 |
| 12 | recombination protein RecR | GB879_11710 | GB880_11420 | The protein RecR that participates in recombinational DNA repair. | Webb et al., 1997 |
| 13 | FtsH protease activity modulator HflK | GB879_11040 | GB880_10470 | The inner membrane proteins HflK and HflC that act together (in a heterodimeric form) as the modulator (negative regulator) of the protease activity of FtsH, a membrane-anchored energy-dependent protease essential for the quality control of integral membrane proteins, regulation of lipopolysaccharide biosynthesis, and regulation of heat shock response. | Kihara et al., 1996; Langklotz et al., 2012 |
| 14 | protease modulator HflC | GB879_11035 | GB880_10465 | The inner membrane proteins HflK and HflC that act together (in a heterodimeric form) as the modulator (negative regulator) of the protease activity of FtsH, a membrane-anchored energy-dependent protease essential for the quality control of integral membrane proteins, regulation of lipopolysaccharide biosynthesis, and regulation of heat shock response. | Kihara et al., 1996; Langklotz et al., 2012 |
| 15 | TerC/Alx family metal homeostasis membrane protein | GB879_10695 | GB880_14190 | Alx, a membrane protein belonging to the TerC family, which facilitates the increase of intracellular manganese (Mn) concentration, thereby playing a key role in Mn homeostasis in conjunction with other ptroteins such as MntP / MntE and UPF0016 that are involved in bacterial Mn transport. Mn constitutes cofactors of enzymes that provide protection against oxidative damage. | Zeinert et al., 2018 |

| | | | | | |
|---|---|---|---|---|---|
| | | | | Nevertheless, excess Mn is toxic, so Mn homeostasis via efficient export-import is central to cellular health (especially under stress conditions). | |
| 16 | recombinase RecA | GB879_02755 | GB880_00745 | The recombinase enzyme RecA of the SOS response machinery, a programmed DNA repair regulatory network that - at the time of environmental stress and adverse effect - not only restores DNA but also confers genetic variability, thereby facilitating adaptation and evolution. | Verbenko, 2017 |
| 17 | BolA/IbaG family iron-sulfur metabolism protein | GB879_06850; GB879_07740 | GB880_06030; GB880_14695 | The BolA/IbaG homologs are reportedly required for maintaining proper cell shape and cell envelope integrity, membrane permeability, motility, and biofilm formation, synthesis and trafficking (assembly and delivery) of iron-sulfur (Fe-S) clusters to a wide variety of proteins (enzymes) that need these evolutionarily ancient cofactors, binding DNA and modulating transcription, and/or modulating bacterial viability amidst diverse environmental stressors. | Guinote et al., 2012, 2014; Dressaire et al., 2015;Fleurie et al., 2019; Talib et al., 2021 |
| 18 | RNA chaperone Hfq | GB879_13320; | GB880_13375 | Hfq renders small-RNA-mediated regulation of target mRNAs (by promoting rapid base-pairing between small RNAs and target mRNAs) to launch quick adaptive response to various stress conditions. The protein is known to modulate $\sigma^{E}$-mediated response to envelope stress and $\sigma^{32}$-mediated response to cytoplasmic stress. | Guisbert et al., 2007; Wagner, 2013 |
| 19 | thioredoxin TrxC | GB879_08690 | GB880_10600 | Bacterial thioredoxin, TrxC, involved in oxidative stress response. *trxC* expression is regulated by the transcriptional activator OxyR in response to oxidative stress: under conditions such as increased hydrogen peroxide concentration and disrupted redox pathway operations, the oxidized activator binds directly to the promoter of the *trxC* gene to induce its expression. | Ritz et al., 2000 |
| 20 | peptide-methionine (R)-S-oxide reductase MsrB | GB879_10530 | GB880_01400 | The thioredoxin-linked enzyme MsrB [peptide-methionine (R)-S-oxide reductase] involved in repairing oxidatively-damaged methionine residues of proteins (conversion of methionine sulfoxide to methionine). | Ezraty et al., 2005 |
| 21 | thioredoxin family protein | GB879_07660 | GB880_07060 | A thioredoxin superfamily member containing a typical TRX domain with a CXXC motif and the members of this family act as protein disulfide oxidoreductases to alter the redox state of other proteins via reversible oxidation of the latters' dithiol sites. | Bardischewsky and Friedrich 2001; Orawski et al., 2007; Carius et al., 2009 |
| 22 | excinuclease ABC subunit | GB879_02880 | GB880_00870 | The subunit A (UvrA) of Excinuclease ABC, also known as UvrABC. UvrA works alongside UvrB, UvrC, DNA helicase II and DNA polymerase I to | Truglio et al., |

| | | | | | |
|---|---|---|---|---|---|
| | UvrA | | | recognize, cleave (in an ATP-dependent manner), and resynthesize or repair damaged DNA. | 2006 |
| 23 | iron-sulfur cluster assembly accessory protein | GB879_03185; GB879_07035 | GB880_01190; GB880_02290 | It is an accessory protein for iron-sulfur cluster assembly. Iron-sulfur (Fe-S) clusters are required for critical biochemical pathways, including respiration, photosynthesis, and nitrogen fixation; while their highly-controlled assembly within bacterial cells help avoid toxicity from free iron and sulfide, multiple assembly pathways carry out basal cluster assembly, stress-responsive cluster assembly, and enzyme-specific cluster assembly. | Ayala-Castro et al., 2008 |
| 24 | nitronate monooxygenase | GB879_00355; GB879_03940 | GB880_00350; GB880_03875 | The nitronate monooxygenase (formerly known as 2-nitropropane dioxygenase), an FMN-dependent enzyme that uses $O_2$ to oxidize (anionic) alkyl nitronates as well as (neutral) nitroalkanes to the corresponding carbonyl compounds and nitrite. Notably, substrates of this enzyme, for example the nitro toxin propionate-3-nitronate, are potent inhibitors of succinate dehydrogenase and fumarase (in the Krebs cycle), so can halt energy production inside the cells. | Gadda and Francis, 2010; Francis et al., 2013 |
| 25 | SH3 domain-containing protein | GB879_03945; GB879_05745; GB879_12520; GB879_12560 | GB880_03870; GB880_05545; GB880_09265; GB880_09305 | SH3-domain-containing proteins are involved in signal transduction, regulation of cell proliferation, migration, and cytoskeletal modifications, via regulation of diverse enzymes, changing the subcellular localization of signaling pathway components, and mediating the formation of multiprotein complex assemblies. | Kurochkina and Guha, 2013 |
| 26 | ribonuclease G | GB879_04715 | GB880_04980 | The non-essential endoribonuclease RNase G that plays a global role in the stability of bacterial mRNAs, besides having potential roles in negatively regulating the expression of heat-shock response genes. | Bernardini et al., 2015; Bernardini and Martínez, 2017 |
| 27 | translation initiation factor IF-1 | GB879_04705 | GB880_04990 | The translation initiation factor IF-1 that consists mainly of an S1 RNA binding domain. IF-1, alongside IF-2 and IF-3, influence the kinetics as well as the stability of the ternary complex formed by the mRNA, fMet-tRNA, and ribosomal subunits. It enhances the rate of association and dissociation of the 70S ribosome subunit, plus the interaction of 30S ribosomal subunit with IF2 and IF3. Furthermore, IF-1 not only stimulates 30S complex formation, but also by binding to the A-site of the 30S ribosomal subunit governs mRNA initiation site selection. | Bycroft et al., 1997; Dahlquist and Puglisi, 2000; Croitoru et al., 2006 |
| 28 | DNA oxidative demethylase | GB879_00320 | GB880_00315 | The 2-oxoglutarate- and iron-dependent DNA repair (damage reversal) enzyme AlkB involved in releasing replication blocks in alkylated DNA, via | Falnes et al., 2002 |

| | | | | | |
|---|---|---|---|---|---|
| | AlkB | | | oxidative demethylation of 1-methyladenine residues. | |
| 29 | metal ABC transporter substrate-binding protein | GB879_01150 | GB880_03465 | The substrate-binding protein of an ATP-binding cassette (ABC) transporter system that putatively carries metal ions. | Maqbool et al., 2015 |
| 30 | RelA/SpoT family protein | GB879_01300 | GB880_01580 | The RelA produces pppGpp (or ppGpp) from ATP and GTP (or GDP), SpoT degrades ppGpp and can act as a ppGpp synthetase too: the two proteins respond specifically to amino acid starvation and diverse starvation stresses respectively. | Battesti and Bouveret, 2009 |
| 31 | DNA repair protein RadC | GB879_09715 | GB880_12510 | The RadC protein involved in repairing DNA damaged by UV and X-ray radiation; in *Escherichia coli* RadC is reported to render a RecG-like DNA recombination/repair function, even though homologs across the bacterial domain may have diversified functions. | Lombardo and Rosenberg, 2000 |
| 32 | DNA mismatch repair protein MutS | GB879_01650; GB879_09545; GB879_10075 | GB880_03820; GB880_08945; GB880_09915 | The MutS protein of the DNA mismatch repair system involved in detecting and refurbishing such nucleotides that have been erroneously inserted, deleted or incorporated during DNA replication, recombination, or damage repair. MutS homodimers initialize the mismatch repair system by detecting and binding to base-base, or insertion/deletion, mismatch-containing DNA. | Iyer et al., 2006 |
| 33 | N-acetylmuramic acid 6-phosphate etherase | GB879_08815 | GB880_08735 | N-acetylmuramic acid 6-phosphate etherase that plays central roles in cell wall metabolism by helping the bacteria grow on MurNAc, besides utilizing extraneous or the cells' own anhydro-N-acetylmuramic acid (that latter function contributes to cell wall recycling). | Uehara et al., 2006; Hadi et al., 2008 |
| 34 | metalloregulator ArsR/SmtB family transcription factor | GB879_04120; GB879_05570; GB879_05610; GB879_07675; GB879_08685; GB879_08750; GB879_08765; | GB880_05675; GB880_05715; GB880_07075; GB880_07545; GB880_10605; GB880_13000 | Metal-sensing transcription repressor of the ArsR/SmtB family, members of which interact with the promoter/operator regions of genes involved in the efflux of diverse metal ions. Upon sensing As+3, Cd+2, Sb+3, Zn+2, etc., SmtB/ArsR repressors dissociate from the promoter/operator regions of the genes that they had repressed thus far and in doing so facilitate the latters' activities which involve chelation, expulsion and/or redox transformation of the metal ions to reduce cellular toxicity. | Busenlehner et al., 2003; Osman and Cavet, 2010 |
| 35 | TetR family transcriptional | GB879_00680; GB879_08120; | GB880_04475; GB880_06655; | TetR family of transcriptional regulators (one component signal transduction proteins) interact with a vast array of ligands, and in doing so control diverse | Cuthbertson and Nodwell, 2013 |

| | | | | | |
|---|---|---|---|---|---|
| | regulator | GB879_08935; GB879_11840; GB879_15485 | GB880_08555; GB880_11625; GB880_14090 | genes involved in a wide variety of bacterial metabolisms such as antibiotic production, quorum sensing, small molecules efflux, cell division and the stress response. | |
| 36 | alpha/beta hydrolase | GB879_02250; GB879_13205 | GB880_04365; GB880_06975 | The α/β-hydrolase fold superfamily that in turn encompasses a wide variety of catalytically distinct entities such as epoxide hydrolases, esterases, chaperones/routers of other proteins, dehalogenases, lipases, lyases, peroxidases, proteases, and transferases and transporters. | Lenfant et al., 2013 |
| 37 | Rrf2 family transcriptional regulator | GB879_02165; GB879_02255; GB879_14685; GB879_15515 | GB880_04360; GB880_11310; GB880_13435; GB880_14860 | A transcriptional regulator (repressor) belonging to the Rrf2 family, members of which are known for redox-sensing under oxidative stress and infection conditions. | Loi et al., 2018 |
| 38 | excinuclease ABC subunit UvrB | GB879_00915 | GB880_03230 | The subunit B (UvrB) of Excinuclease ABC, also known as UvrABC. UvrB works alongside UvrA, UvrC, DNA helicase II and DNA polymerase I to recognize, cleave (in an ATP-dependent manner), and resynthesize or repair damaged DNA. | Truglio et al., 2006 |
| 39 | Do family serine endopeptidase | GB879_04280; GB879_04630 | GB880_02060; GB880_04655 | The Peptidase Do, a serine endopeptidase that is essential for degradation and clearing partially-unfolded or denatured proteins from the inner-membrane and periplasmic space, especially under high temperature conditions. | Seol et al., 1991 |
| 40 | porin | GB879_09415; GB879_10650 | GB880_09080; GB880_10195 | The porins are large transport proteins that allow passive diffusion and facilitate specific or non-specific entry of hydrophillic molecules into the cell. | Cabiaux et al., 2003 |
| 41 | Grx4 family monothiol glutaredoxin | GB879_06845 | GB880_06035 | A Grx4 family monothiol glutaredoxin that acts as an important regulator of overall redox, and iron, homeostasis of the cell, and is potentially regulated by guanosine 3′,5′-tetraphosphate (ppGpp) in stationary phase cells. | Fernandes et al., 2005; Attarian et al., 2018 |
| 42 | RNA-binding protein | GB879_02395; GB879_05450 | GB880_04215; GB880_05395 | A member of the YlxR group of conserved bacterial proteins that can bind RNA, and influence transcription, recombination, and genome stability. YlxR homologs also critically influence the expression of a large number of metabolic genes and sigma factor genes. | Ogura and Kanesaki, 2018 |
| 43 | nucleoside-diphosphate kinase | GB879_00415 | GB880_00410 | The nucleoside diphosphate kinase (NdK), an enzyme that keeps the concentrations of different nucleoside triphosphates within a cell in equilibrium by catalyzing the exchange of terminal phosphate between different nucleoside triphosphates (NTP) and diphosphates (NDP) in a reversible manner to produce new nucleoside triphosphates (XDP + YTP ←→ XTP + YDP, where X | Chakrabarty, 1998 |

| | | | | | |
|---|---|---|---|---|---|
| | | | | and Y represent different nitrogenous bases; consequently, NdK regulates the synthesis of cell surface polysaccharides and overall cellular growth. | |
| 44 | protein TolQ | GB879_11480 | GB880_11190 | The inner membrane protein TolQ that together with the other components of the Tol-Pal system (the inner membrane proteins TolA and TolR, the periplasmic TolB and the outer membrane protein Pal) form a network linking the inner and outer membranes and the peptidoglycan layer. Overexpression of TolQ in Gram-negative bacteria binds and sequesters the essential cell division protein FtsN, thereby depleting this key component of the divisome that is needed to activate septal peptidoglycan synthesis (division septum assembly) and constriction of the dividing cell. | Derouiche et al., 1995; Journet et al., 1999; Teleha et al., 2013; Liu et al., 2015 |
| 45 | protein-methionine-sulfoxide reductase catalytic subunit MsrP | GB879_14555 | GB880_14625 | MsrP, the catalytic subunit of protein-methionine sulfoxide reductase; MsrP is a part of the MsrPQ system that repairs oxidized periplasmic proteins containing methionine sulfoxide residues (Met-O), using respiratory chain electrons, thereby protecting these proteins from damage caused by reactive oxygen species. MsrPQ is essential for the maintenance of envelope integrity under stress, as it rescues diverse structurally-unrelated periplasmic proteins, such as the primary periplasmic chaperone SurA and the outer membrane lipoprotein Pal, from methionine oxidation. | Gennaris et al., 2015 |
| 46 | L,D-transpeptidase family protein | GB879_01220; GB879_01740; GB879_05510; GB879_06855; GB879_07215; GB879_11560; GB879_11890; GB879_12440; GB879_14030 | GB880_01500; GB880_03730; GB880_05455; GB880_06025; GB880_07220; GB880_09440; GB880_11270; GB880_13255; GB880_15010 | An L,D-transpeptidase, homologs of which render the unusual 3→3 (diaminopimelic acid → diaminopimelic acid) cross-linking between the short peptides of the peptidoglycan. Amidst elevated synthesis of the (p)ppGpp alarmone by RelA, L,D-transpeptidases can even act as the main peptidoglycan cross-linking enzyme in a bacterium, thereby enabling it to completely bypass the usual 4→3 (D-alanine → diaminopimelic acid) cross-linking activity of the D,D-transpeptidase. | Triboulet et al., 2013; Hugonnet et al., 2016 |
| 47 | protein-export chaperone SecB | GB879_09560 | GB880_08930 | The protein-export chaperone SecB required for the translocation of a large variety of pre-proteins or precursor proteins out of the cytoplasm in loosely folded but stable conformation; mechanistically, the role of SecB is to assist in the targeting of secretory proteins to the Sec-translocase. | Baars et al., 2006; Huang et al., 2016 |
| 48 | 4-hydroxyphenylp | GB879_14950 | GB880_15445 | 4-hydroxyphenylpyruvate dioxygenase responsible for converting 4-hydroxyphenylpyruvate to homogentisate. With decline in $O_2$ concentration, as | Coon et al., 1994; Ruzafa et al., |

| | | | | | |
|---|---|---|---|---|---|
| | yruvate dioxygenase | | | witnessed with increase in temperature, auto-oxidation and self-polymerization of homogentisate can lead to the formation of pyomelanin, a polyaromatic heteropolymer, which consists of numerous quinone moieties. Melanin pigments, in turn, confer versatile growth and survival advantages to microorganisms; these include protection against UV light, oxidative stress, and ionizing radiations; acting as a terminal electron acceptor; enhancing dissimilatory $Fe^{3+}$ reduction as well as $Fe^{2+}$ chelation; uranium complexation and immobilization. | 1995; Turick et al., 2009 |
| 49 | peptidylprolyl isomerase | GB879_02695; GB879_05110; GB879_05420; GB879_06020; GB879_13065; GB879_13070 | GB880_05050; GB880_05365; GB880_10860; GB880_11075; GB880_11080; | The peptidylprolyl isomerase (PPIase), a ubiquitous enzyme encountered across the three domains of life, and involved in catalyzing the *cis–trans* isomerisation of peptide bonds that are N-terminal to proline (Pro) residues within polypeptide chains. In doing so, PPIs often play chaperone-like roles in the folding of many newly synthesised proteins, and in turn control a wide range of cellular activities. | Fischer and Schmid, 1990; Schmid, 1995 |
| 50 | M48 family metalloprotease | GB879_00870; GB879_08635 | GB880_06610; GB880_12550; GB880_15910 | An M48 family metalloprotease; members of M48 have chaperone activities, play important roles in the maturation and quality-control of lipopolysaccharide besides influencing the insertion of other OMPs into the outer membrane. | Narita et al., 2013; Bryant et al., 2020 |

### Supplementary References


Allan, E., Mullany, P. and Tabaqchali, S., 1998. Construction and characterization of a *Helicobacter pylori* clpB mutant and role of the gene in the stress response. *Journal of bacteriology*, *180*, 426-429. https://doi.org/10.1128/JB.180.2.426-429.1998

Attarian, R., Hu, G., Sánchez-León, E., Caza, M., Croll, D., Do, E., Bach, H., Missall, T., Lodge, J., Jung, W.H. and Kronstad, J.W., 2018. The monothiol glutaredoxin Grx4 regulates iron homeostasis and virulence in *Cryptococcus neoformans*. *MBio*, *9*, e02377-18. https://doi.org/10.1128/mBio.02377-18

Ayala-Castro, C., Saini, A. and Outten, F.W., 2008. Fe-S cluster assembly pathways in bacteria. *Microbiology and Molecular Biology Reviews*, *72*, 110-125. https://doi.org/10.1128/MMBR.00034-07

Baars, L., Ytterberg, A.J., Drew, D., Wagner, S., Thilo, C., Van Wijk, K.J. and de Gier, J.W., 2006. Defining the role of the *Escherichia coli* chaperone SecB using comparative proteomics. *Journal of Biological Chemistry*, *281*, 10024-10034. https://doi.org/10.1074/jbc.M509929200

Barbirz, S., Jakob, U., Glocker, M.O., 2000. Mass spectrometry unravels disulfide bond formation as the mechanism that activates a molecular chaperone. J. Biol. Chem. 275, 18759-18766. https://doi.org/10.1074/jbc.M001089200

Bardischewsky, F. and Friedrich, C.G., 2001. The shxVW locus is essential for oxidation of inorganic sulfur and molecular hydrogen by *Paracoccus pantotrophus* GB17: a novel function for lithotrophy. *FEMS microbiology letters*, *202*, 215-220. https://doi.org/10.1111/j.1574-6968.2001.tb10806.x

Battesti, A. and Bouveret, E., 2009. Bacteria possessing two RelA/SpoT-like proteins have evolved a specific stringent response involving the acyl carrier protein-SpoT interaction. *Journal of bacteriology*, *191*, 616-624. https://doi.org/10.1128/jb.01195-08

Bernardini, A. and Martínez, J.L., 2017. Genome-wide analysis shows that RNase G plays a global role in the stability of mRNAs in *Stenotrophomonas maltophilia*. *Scientific reports*, *7*, 16016. https://doi.org/10.1038/s41598-017-16091-0

Bernardini, A., Corona, F., Dias, R., Sánchez, M.B. and Martínez, J.L., 2015. The inactivation of RNase G reduces the Stenotrophomonas maltophilia susceptibility to quinolones by triggering the heat shock response. *Frontiers in microbiology*, *6*, 1068. https://doi.org/10.3389/fmicb.2015.01068

Bryant, J.A., Cadby, I.T., Chong, Z.S., Boelter, G., Sevastsyanovich, Y.R., Morris, F.C., Cunningham, A.F., Kritikos, G., Meek, R.W., Banzhaf, M. and Chng, S.S., 2020. Structure-function characterization of the conserved regulatory mechanism of the *Escherichia coli* M48 metalloprotease BepA. *Journal of bacteriology*, *203*, e00434-20. https://doi.org/10.1128/JB.00434-20

Busenlehner, L.S., Pennella, M.A. and Giedroc, D.P., 2003. The SmtB/ArsR family of metalloregulatory transcriptional repressors: structural insights into prokaryotic metal resistance. *FEMS microbiology reviews*, *27*, 131-143. https://doi.org/10.1016/S0168-6445(03)00054-8

Bycroft, M., Hubbard, T.J., Proctor, M., Freund, S.M. and Murzin, A.G., 1997. The solution structure of the S1 RNA binding domain: a member of an ancient nucleic acid–binding fold. *Cell*, *88*, 235-242. https://doi.org/10.1016/S0092-8674(00)81844-9

Cabiaux, V., Vande Weyer, S. and Ruysschaert, J.M., 2003. Chapter 18 - The use of iiposomes to detect channel formation mediated by secreted bacterial proteins. Editor(s): H.T. Tien, A. Ottova-Leitmannova. *Membrane science and technology*, Elsevier, Volume *7*, pp.517-537. https://doi.org/10.1016/S0927-5193(03)80042-6

Carius, Y., Rother, D., Friedrich, C.G. and Scheidig, A.J., 2009. The structure of the periplasmic thiol–disulfide oxidoreductase SoxS from *Paracoccus pantotrophus* indicates a triple Trx/Grx/DsbC functionality in chemotrophic sulfur oxidation. *Acta Crystallographica Section D: Biological Crystallography*, *65*, 229-240. https://doi.org/10.1107/S0907444908043023

Chakrabarty, A.M., 1998. Nucleoside diphosphate kinase: role in bacterial growth, virulence, cell signalling and polysaccharide synthesis. *Molecular microbiology*, *28*, 875-882. https://doi.org/10.1046/j.1365-2958.1998.00846.x

Coon, S.L., Kotob, S., Jarvis, B.B., Wang, S., Fuqua, W.C. and Weiner, R.M., 1994. Homogentisic acid is the product of MelA, which mediates melanogenesis in the marine bacterium *Shewanella* *colwelliana* D. *Applied and environmental microbiology*, *60*, 3006-3010. https://doi.org/10.1128/aem.60.8.3006-3010.1994

Croitoru, V., Semrad, K., Prenninger, S., Rajkowitsch, L., Vejen, M., Laursen, B.S., Sperling-Petersen, H.U. and Isaksson, L.A., 2006. RNA chaperone activity of translation initiation factor IF1. *Biochimie*, *88*, 1875-1882. https://doi.org/10.1016/j.biochi.2006.06.017

Cuthbertson, L. and Nodwell, J.R., 2013. The TetR family of regulators. *Microbiology and Molecular Biology Reviews*, *77*, 440-475. https://doi.org/10.1128/MMBR.00018-13

Dahlquist, K.D. and Puglisi, J.D., 2000. Interaction of translation initiation factor IF1 with the *E. coli* ribosomal A site. *Journal of molecular biology*, *299*, 1-15. https://doi.org/10.1006/jmbi.2000.3672

De Almeida, A., Nikel, P.I., Giordano, A.M. and Pettinari, M.J., 2007. Effects of granule-associated protein PhaP on glycerol-dependent growth and polymer production in poly (3-hydroxybutyrate)-producing *Escherichia coli*. *Applied and environmental microbiology*, *73*, 7912-7916. https://doi.org/10.1128/AEM.01900-07

Derouiche, R., Benedetti, H., Lazzaroni, J.C., Lazdunski, C. and Lloubès, R., 1995. Protein complex within *Escherichia coli* inner membrane: TolA N-terminal domain interacts with TolQ and TolR proteins. *Journal of Biological Chemistry*, *270*, 11078-11084. https://doi.org/10.1074/jbc.270.19.11078

Dressaire, C., Moreira, R.N., Barahona, S., Alves de Matos, A.P. and Arraiano, C.M., 2015. BolA is a transcriptional switch that turns off motility and turns on biofilm development. *MBio*, *6*, e02352-14. https://doi.org/10.1128/mBio.02352-14

Ezraty, B., Aussel, L. and Barras, F., 2005. Methionine sulfoxide reductases in prokaryotes. *Biochimica et Biophysica Acta (BBA)-Proteins and Proteomics*, *1703*, 221-229. https://doi.org/10.1016/j.bbapap.2004.08.017

Falnes, P.Ø., Johansen, R.F. and Seeberg, E., 2002. AlkB-mediated oxidative demethylation reverses DNA damage in *Escherichia coli*. *Nature*, *419*, 178-182. https://doi.org/10.1038/nature01048

Fernandes, A.P., Fladvad, M., Berndt, C., Andrésen, C., Lillig, C.H., Neubauer, P., Sunnerhagen, M., Holmgren, A. and Vlamis-Gardikas, A., 2005. A novel monothiol glutaredoxin (Grx4) from *Escherichia coli* can serve as a substrate for thioredoxin reductase. *Journal of Biological Chemistry*, *280*, 24544-24552. https://doi.org/10.1074/jbc.M500678200

Fischer, G. and Schmid, F.X., 1990. The mechanism of protein folding. Implications of in vitro refolding models for de novo protein folding and translocation in the cell. *Biochemistry*, *29*, 2205-2212. https://doi.org/10.1021/bi00461a001

Fleurie, A., Zoued, A., Alvarez, L., Hines, K.M., Cava, F., Xu, L., Davis, B.M. and Waldor, M.K., 2019. A *Vibrio cholerae* BolA-like protein is required for proper cell shape and cell envelope integrity. *MBio*, *10*, e00790-19. https://doi.org/10.1128/mBio.00790-19

Francis, K., Smitherman, C., Nishino, S.F., Spain, J.C. and Gadda, G., 2013. The biochemistry of the metabolic poison propionate 3-nitronate and its conjugate acid, 3-nitropropionate. *IUBMB life*, *65*, 759-768. https://doi.org/10.1002/iub.1195

Gadda, G. and Francis, K., 2010. Nitronate monooxygenase, a model for anionic flavin semiquinone intermediates in oxidative catalysis. *Archives of biochemistry and biophysics*, *493*, 53-61. https://doi.org/10.1016/j.abb.2009.06.018

Genevaux, P., Keppel, F., Schwager, F., Langendijk-Genevaux, P.S., Hartl, F.U., Georgopoulos, C., 2004. In vivo analysis of the overlapping functions of DnaK and trigger factor. EMBO Rep. 5, 195-200. https://doi.org/10.1038/sj.embor.7400067

Gennaris, A., Ezraty, B., Henry, C., Agrebi, R., Vergnes, A., Oheix, E., Bos, J., Leverrier, P., Espinosa, L., Szewczyk, J. and Vertommen, D., 2015. Repairing oxidized proteins in the bacterial envelope using respiratory chain electrons. *Nature*, *528*, 409-412. https://doi.org/10.1038/nature15764

Ghazaei, C., 2017. Role and mechanism of the Hsp70 molecular chaperone machines in bacterial pathogens. Int. J. Med. Microbiol. 66, 259-265. https://doi.org/10.1099/jmm.0.000429

Guinote, I.B., Moreira, R.N., Barahona, S., Freire, P., Vicente, M. and Arraiano, C.M., 2014. Breaking through the stress barrier: the role of BolA in Gram-negative survival. *World Journal of Microbiology and Biotechnology*, *30*, 2559-2566. https://doi.org/10.1007/s11274-014-1702-4

Guinote, I.B., Moreira, R.N., Freire, P. and Arraiano, C.M., 2012. Characterization of the BolA homolog IbaG: a new gene involved in acid resistance. *Journal of microbiology and biotechnology*, *22*, 484-493. https://doi.org/10.4014/jmb.1107.07037

Guisbert, E., Rhodius, V.A., Ahuja, N., Witkin, E. and Gross, C.A., 2007. Hfq modulates the σE-mediated envelope stress response and the σ32-mediated cytoplasmic stress response in *Escherichia coli*. *Journal of bacteriology*, *189*, 1963-1973. https://doi.org/10.1128/JB.01243-06

Hadi, T., Dahl, U., Mayer, C. and Tanner, M.E., 2008. Mechanistic studies on N-acetylmuramic acid 6-phosphate hydrolase (MurQ): an etherase involved in peptidoglycan recycling. *Biochemistry*, *47*, 11547-11558. https://doi.org/10.1021/bi8014532

Harrison, C.J., Hayer-Hartl, M., Di Liberto, M., Hartl, F.U., Kuriyan, J., 1997. Crystal structure of the nucleotide exchange factor GrpE bound to the ATPase domain of the molecular chaperone DnaK. Science 276, 431-435. https://doi.org/10.1126/science.276.5311.431

Hitomi, M., Nishimura, H., Tsujimoto, Y., Matsui, H., Watanabe, K., 2003. Identification of a helix-turn-helix motif of *Bacillus thermoglucosidasius* HrcA essential for binding to the CIRCE element and thermostability of the HrcA-CIRCE complex, indicating a role as a thermosensor. J. Bacteriol. 185, 381-385. https://doi.org/10.1128/JB.185.1.381-385.2003

Huang, C., Rossi, P., Saio, T. and Kalodimos, C.G., 2016. Structural basis for the antifolding activity of a molecular chaperone. *Nature*, *537*, 202-206. https://doi.org/10.1038/nature18965

Hugonnet, J.E., Mengin-Lecreulx, D., Monton, A., den Blaauwen, T., Carbonnelle, E., Veckerle, C., Yves, V.B., van Nieuwenhze, M., Bouchier, C., Tu, K. and Rice, L.B., 2016. Factors essential for L, D-transpeptidase-mediated peptidoglycan cross-linking and β-lactam resistance in *Escherichia coli*. *Elife*, *5*, e19469. https://doi.org/10.7554/eLife.19469.001

Iyer, R.R., Pluciennik, A., Burdett, V. and Modrich, P.L., 2006. DNA mismatch repair: functions and mechanisms. *Chemical reviews*, *106*, 302-323. https://doi.org/10.1021/cr0404794

Jonas, K., Liu, J., Chien, P. and Laub, M.T., 2013. Proteotoxic stress induces a cell-cycle arrest by stimulating Lon to degrade the replication initiator DnaA. *Cell*, *154*, 623-636. https://doi.org/10.1016/j.cell.2013.06.034

Journet, L., Rigal, A., Lazdunski, C. and Bénédetti, H., 1999. Role of TolR N-terminal, central, and C-terminal domains in dimerization and interaction with TolA and TolQ. *Journal of bacteriology*, *181*, 4476-4484. https://doi.org/10.1128/JB.181.15.4476-4484.1999

Karzai, A.W., Roche, E.D. and Sauer, R.T., 2000. The SsrA–SmpB system for protein tagging, directed degradation and ribosome rescue. *Nature structural biology*, *7*, 449-455. https://doi.org/10.1038/75843

Katz, C. and Ron, E.Z., 2008. Dual role of FtsH in regulating lipopolysaccharide biosynthesis in Escherichia coli. *Journal of bacteriology*, *190*(21), pp.7117-7122.

Kihara, A., Akiyama, Y. and Ito, K., 1996. A protease complex in the *Escherichia coli* plasma membrane: HflKC (HflA) forms a complex with FtsH (HflB), regulating its proteolytic activity against SecY. *The EMBO Journal*, *15*, 6122-6131. https://doi.org/10.1002/j.1460-2075.1996.tb01000.x

Kudlicki, W., Coffman, A., Kramer, G. and Hardesty, B., 1997. Renaturation of rhodanese by translational elongation factor (EF) Tu: protein refolding by EF-Tu flexing. *Journal of Biological Chemistry*, *272*, 32206-32210. https://doi.org/10.1074/jbc.272.51.32206

Kurochkina, N. and Guha, U., 2013. SH3 domains: modules of protein–protein interactions. *Biophysical reviews*, *5*, 29-39. https://doi.org/10.1007/s12551-012-0081-z

Langklotz, S., Baumann, U. and Narberhaus, F., 2012. Structure and function of the bacterial AAA protease FtsH. *Biochimica Et Biophysica Acta (BBA)-Molecular Cell Research*, *1823*, 40-48. https://doi.org/10.1016/j.bbamcr.2011.08.015

Lee, S., Sowa, M.E., Watanabe, Y.H., Sigler, P.B., Chiu, W., Yoshida, M. and Tsai, F.T., 2003. The structure of ClpB: a molecular chaperone that rescues proteins from an aggregated state. *Cell*, *115*, 229-240. https://doi.org/10.1016/S0092-8674(03)00807-9

Lenfant, N., Hotelier, T., Bourne, Y., Marchot, P. and Chatonnet, A., 2013. Proteins with an alpha/beta hydrolase fold: relationships between subfamilies in an ever-growing superfamily. *Chemico-biological interactions*, *203*, 266-268. https://doi.org/10.1016/j.cbi.2012.09.003

Li, D.C., Yang, F., Lu, B., Chen, D.F., Yang, W.J., 2012. Thermotolerance and molecular chaperone function of the small heat shock protein HSP20 from hyperthermophilic archaeon, *Sulfolobus solfataricus* P2. Cell Stress Chaperones17, 103-108. https://doi.org/10.1007/s12192-011-0289-z

Liberek, K., Marszalek, J., Ang, D., Georgopoulos, C., Zylicz, M., 1991. *Escherichia coli* DnaJ and GrpE heat shock proteins jointly stimulate ATPase activity of DnaK. Proc. Natl. Acad. Sci. USA. 88, 2874-2878. https://doi.org/10.1073/pnas.88.7.2874

Liu, B., Persons, L., Lee, L. and de Boer, P.A., 2015. Roles for both FtsA and the FtsBLQ subcomplex in FtsN-stimulated cell constriction in *Escherichia coli*. *Molecular microbiology*, *95*, 945-970. https://doi.org/10.1111/mmi.12906

Loi, V.V., Busche, T., Tedin, K., Bernhardt, J., Wollenhaupt, J., Huyen, N.T.T., Weise, C., Kalinowski, J., Wahl, M.C., Fulde, M. and Antelmann, H., 2018. Redox-sensing under hypochlorite stress and infection conditions by the Rrf2-family repressor HypR in *Staphylococcus aureus*. *Antioxidants & redox signaling*, *29*, 615-636. https://doi.org/10.1089/ars.2017.7354

Lombardo, M.J. and Rosenberg, S.M., 2000. radC102 of *Escherichia coli* is an allele of recG. *Journal of bacteriology*, *182*, 6287-6291. https://doi.org/10.1128/JB.182.22.6287-6291.2000

Maehara, A., Taguchi, S., Nishiyama, T., Yamane, T. and Doi, Y., 2002. A repressor protein, PhaR, regulates polyhydroxyalkanoate (PHA) synthesis via its direct interaction with PHA. *Journal of bacteriology*, *184*, 3992-4002. https://doi.org/10.1128/JB.184.14.3992-4002.2002

Malde, A., Gangaiah, D., Chandrashekhar, K., Pina-Mimbela, R., Torrelles, J.B. and Rajashekara, G., 2014. Functional characterization of exopolyphosphatase/guanosine pentaphosphate phosphohydrolase (PPX/GPPA) of *Campylobacter jejuni*. *Virulence*, *5*, 521-533. https://doi.org/10.4161/viru.28311

Maleki, F., Afra Khosravi, A.N., Taghinejad, H., Azizian, M., 2016. Bacterial heat shock protein activity. J. Clin. Diagn. Res. 10, BE01-BE03. https://doi.org/10.7860/JCDR/2016/14568.7444

Maqbool, A., Horler, R.S., Muller, A., Wilkinson, A.J., Wilson, K.S. and Thomas, G.H., 2015. The substrate-binding protein in bacterial ABC transporters: dissecting roles in the evolution of substrate specificity. *Biochemical Society Transactions*, *43*, 1011-1017. https://doi.org/10.1042/BST20150135

Moreno-Cinos, C., Goossens, K., Salado, I.G., Van Der Veken, P., De Winter, H. and Augustyns, K., 2019. ClpP protease, a promising antimicrobial target. *International journal of molecular sciences*, *20*, 2232. https://doi.org/10.3390/ijms20092232

Narita, S.I., Masui, C., Suzuki, T., Dohmae, N. and Akiyama, Y., 2013. Protease homolog BepA (YfgC) promotes assembly and degradation of β-barrel membrane proteins in *Escherichia coli*. *Proceedings of the National Academy of Sciences of the United States of America*, *110*, E3612-E3621. https://doi.org/10.1073/pnas.1312012110

Ogura, M. and Kanesaki, Y., 2018. Newly identified nucleoid-associated-like protein YlxR regulates metabolic gene expression in *Bacillus subtilis*. *Msphere*, *3*, e00501-18. https://doi.org/10.1128/mSphere.00501-18

Orawski, G., Bardischewsky, F., Quentmeier, A., Rother, D. and Friedrich, C.G., 2007. The periplasmic thioredoxin SoxS plays a key role in activation in vivo of chemotrophic sulfur oxidation of *Paracoccus pantotrophus*. *Microbiology*, *153*, 1081-1086. https://doi.org/10.1099/mic.0.2006/004143-0

Osman, D. and Cavet, J.S., 2010. Bacterial metal-sensing proteins exemplified by ArsR–SmtB family repressors. *Natural product reports*, *27*, 668-680. https://doi.org/10.1039/B906682A

Pandey, D.P. and Gerdes, K., 2005. Toxin–antitoxin loci are highly abundant in free-living but lost from host-associated prokaryotes. *Nucleic acids research*, *33*, 966-976. https://doi.org/10.1093/nar/gki201

Pinti, M., Gibellini, L., Nasi, M., De Biasi, S., Bortolotti, C.A., Iannone, A. and Cossarizza, A., 2016. Emerging role of Lon protease as a master regulator of mitochondrial functions. *Biochimica et Biophysica Acta (BBA)-Bioenergetics*, *1857*, 1300-1306. https://doi.org/10.1016/j.bbabio.2016.03.025

Raina, S., Missiakas, D., Georgopoulos, C., 1995. The rpoE gene encoding the sigma E (sigma 24) heat shock sigma factor of *Escherichia coli*. EMBO J. 14, 1043-1055. https://doi.org/10.1002/j.1460-2075.1995.tb07085.x

Reizer, J., Reizer, A., Saier Jr, M.H., Bork, P. and Sander, C., 1993. Exopolyphosphate phosphatase and guanosine pentaphosphate phosphatase belong to the sugar kinase/actin/hsp 70 superfamily. *Trends in biochemical sciences*, *18*, 247-248. https://doi.org/10.1016/0968-0004(93)90172-j

Ritz, D., Patel, H., Doan, B., Zheng, M., Åslund, F., Storz, G. and Beckwith, J., 2000. Thioredoxin 2 Is Involved in the Oxidative Stress Response in *Escherichia coli*. *Journal of Biological Chemistry*, *275*, 2505-2512. https://doi.org/10.1074/jbc.275.4.2505

Roy, C., Mondal, N., Peketi, A., Fernandes, S., Mapder, T., Volvoikar, S.P., Haldar, P.K., Nandi, N., Bhattacharya, T., Mazumdar, A., Chakraborty, R. and Ghosh, W. 2020a. Geomicrobial dynamics of Trans-Himalayan sulfur–borax spring system reveals mesophilic bacteria's resilience to high heat. *Journal of Earth System Science,* 129:157. https://doi.org/10.1007/s12040-020-01423-y

Ruzafa, C., Sanchez-amat, A. and Solano, F., 1995. Characterization of the melanogenic system in *Vibrio cholerae*, ATCC 14035. *Pigment Cell Research*, *8*, 147-152. https://doi.org/10.1111/j.1600-0749.1995.tb00656.x

Sauer, R.T. and Baker, T.A., 2011. AAA+ proteases: ATP-fueled machines of protein destruction. *Annual review of biochemistry*, *80*, 587-612. https://doi.org/10.1146/annurev-biochem-060408-172623

Schmid, F.X., 1995. Protein folding: prolyl isomerases join the fold. *Current Biology*, *5*, 993-994. https://doi.org/10.1016/S0960-9822(95)00197-7

Seol, J.H., Woo, S.K., Jung, E.M., Yoo, S.J., Lee, C.S., Kim, K., Tanaka, K., Ichihara, A., Ha, D.B. and Chung, C.H., 1991. Protease Do is essential for survival of *Escherichia coli* at high temperatures: its identity with the htrA gene product. *Biochemical and biophysical research communications*, *176*, 730-736. https://doi.org/10.1016/s0006-291x(05)80245-1

Shimuta, T.R., Nakano, K., Yamaguchi, Y., Ozaki, S., Fujimitsu, K., Matsunaga, C., Noguchi, K., Emoto, A., Katayama, T., 2004. Novel heat shock protein HspQ stimulates the degradation of mutant DnaA protein in *Escherichia coli*. Genes Cells 9, 1151-1166. https://doi.org/10.1111/j.1365-2443.2004.00800.x

Srivatsan, A. and Wang, J.D., 2008. Control of bacterial transcription, translation and replication by (p)ppGpp. *Current opinion in microbiology*, *11*, 100-105. https://doi.org/10.1016/j.mib.2008.02.001

Straus, D.B., Walter, W.A., Gross, C.A., 1987. The heat shock response of *E. coli* is regulated by changes in the concentration of σ32. Nature 329, 348-351. https://doi.org/10.1038/329348a0

Talib, E.A. and Outten, C.E., 2021. Iron-sulfur cluster biogenesis, trafficking, and signaling: Roles for CGFS glutaredoxins and BolA proteins. *Biochimica et Biophysica Acta (BBA)-Molecular Cell Research*, *1868*, 118847. https://doi.org/10.1016/j.bbamcr.2020.118847

Teleha, M.A., Miller, A.C. and Larsen, R.A., 2013. Overexpression of the *Escherichia coli* TolQ protein leads to a null-FtsN-like division phenotype. *Microbiologyopen*, *2*, 618-632. https://doi.org/10.1002/mbo3.101

Triboulet, S., Dubée, V., Lecoq, L., Bougault, C., Mainardi, J.L., Rice, L.B., Ethève-Quelquejeu, M., Gutmann, L., Marie, A., Dubost, L. and Hugonnet, J.E., 2013. Kinetic features of L, D-transpeptidase inactivation critical for β-lactam antibacterial activity. *PloS one*, *8*, e67831. https://doi.org/10.1371/journal.pone.0067831

Truglio, J.J., Croteau, D.L., Van Houten, B. and Kisker, C., 2006. Prokaryotic nucleotide excision repair: the UvrABC system. *Chemical reviews*, *106*, 233-252. https://doi.org/10.1021/cr040471u

Turick, C.E., Beliaev, A.S., Zakrajsek, B.A., Reardon, C.L., Lowy, D.A., Poppy, T.E., Maloney, A. and Ekechukwu, A.A., 2009. The role of 4-hydroxyphenylpyruvate dioxygenase in enhancement of solid-phase electron transfer by *Shewanella oneidensis* MR-1. *FEMS microbiology ecology*, *68*, 223-235. https://doi.org/10.1111/j.1574-6941.2009.00670.x

Uehara, T., Suefuji, K., Jaeger, T., Mayer, C. and Park, J.T., 2006. MurQ etherase is required by *Escherichia coli* in order to metabolize anhydro-N-acetylmuramic acid obtained either from the environment or from its own cell wall. *Journal of bacteriology*, *188*, 1660-1662. https://doi.org/10.1128/jb.188.4.1660-1662.2006

Verbenko, V., 2017. SOS Repair, In Reference Module in Life Sciences, Elsevier, Amsterdam, Netherlands https://doi.org/10.1016/B978-0-12-809633-8.07178-8

Wagner, E.G.H., 2013. Cycling of RNAs on hfq. *RNA biology*, *10*, 619-626. https://doi.org/10.4161/rna.24044

Webb, B.L., Cox, M.M. and Inman, R.B., 1997. Recombinational DNA repair: the RecF and RecR proteins limit the extension of RecA filaments beyond single-strand DNA gaps. *Cell*, *91*, 347-356. https://doi.org/10.1016/s0092-8674(00)80418-3

Weissbach, H., Etienne, F., Hoshi, T., Heinemann, S.H., Lowther, W.T., Matthews, B., John, G.S., Nathan, C. and Brot, N., 2002. Peptide methionine sulfoxide reductase: structure, mechanism of action, and biological function. *Archives of Biochemistry and Biophysics*, *397*(2), pp.172-178. https://doi.org/10.1006/abbi.2001.2664

Wieden, H.J., Gromadski, K., Rodnin, D. and Rodnina, M.V., 2002. Mechanism of elongation factor (EF)-Ts-catalyzed nucleotide exchange in EF-Tu: contribution of contacts at the guanine base. *Journal of Biological Chemistry*, *277*, 6032-6036. https://doi.org/10.1074/jbc.M110888200

Zeinert, R., Martinez, E., Schmitz, J., Senn, K., Usman, B., Anantharaman, V., Aravind, L. and Waters, L.S., 2018. Structure–function analysis of manganese exporter proteins across bacteria. *Journal of Biological Chemistry*, *293*, 5715-5730. https://doi.org/10.1074/jbc.M117.790717

### Supplementary Figures

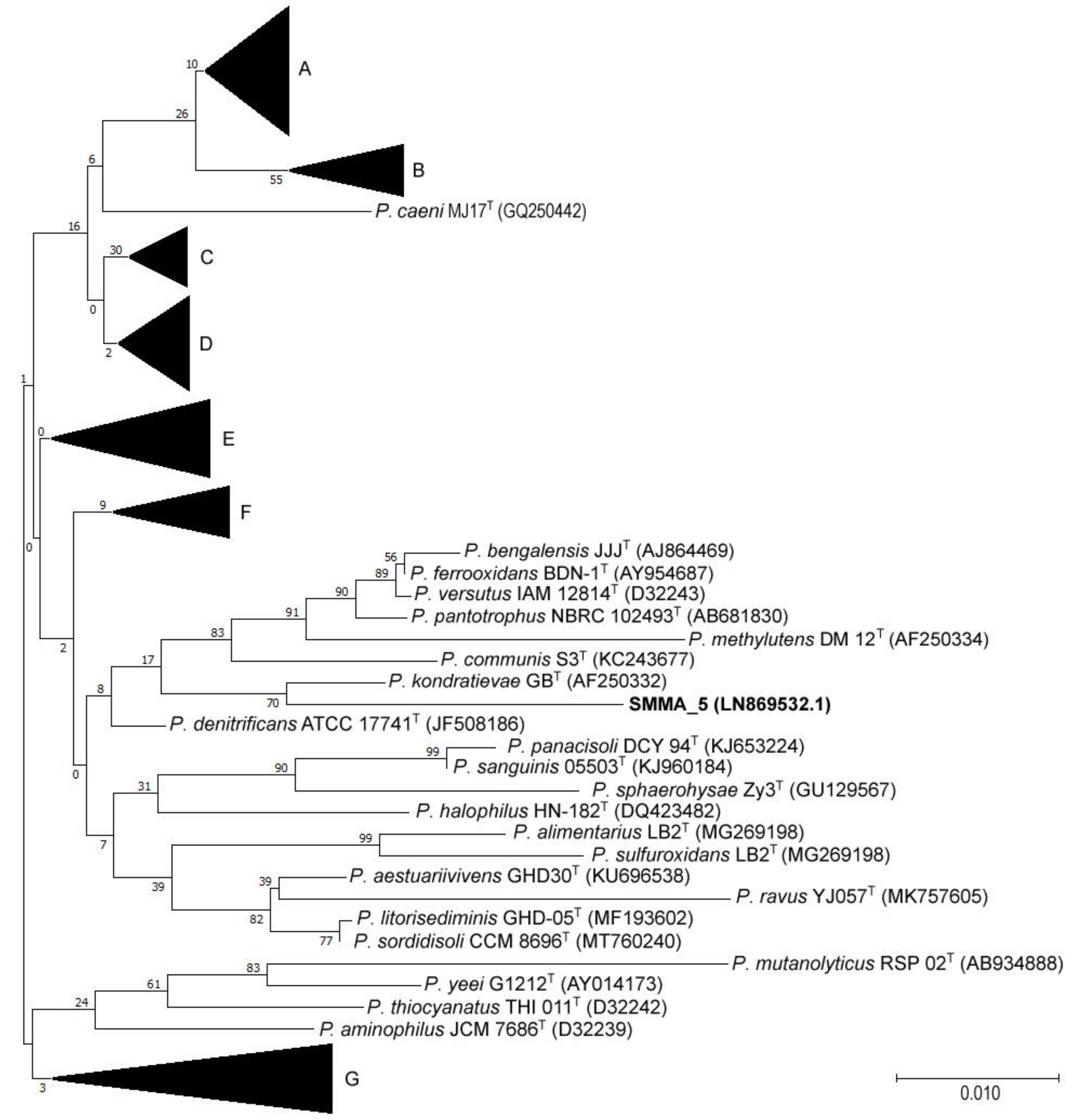


**Figure S1.** 16S rRNA gene sequence based neighbor-joining tree showing the phylogenetic relationships between the new isolate SMMA_5 and representative strains of existing *Paracoccus* species. The distance bar represents 10% nucleotide difference. Tree branches were condensed and marked as clades A to G. **Clade A** encompassed *P. haeundaensis* BC74171$^T$ (AY189743), *P. marcusii* MH1$^T$ (Y12703), *P. carotinifaciens* E396$^T$ (AB006899), *P. hibiscisoli* THG-T2.31$^T$ (KX456191), *P. nototheniae* 41R45$^T$ (MH065728), *P. gahaiensis* CUG 00006$^T$ (KT345705), *P. liaowanqingii* 2251$^T$ (MG798927), *P. seriniphilus* NBRC 100798$^T$ (AB681242), *P. fistulariae* 22-5$^T$ (GQ260189), *P. homiensis* DD-R11$^T$ (DQ342239), *P. halotolerans* CFH 90064$^T$ (KY039331),*P. zeaxanthinifaciens* ATCC 21588$^T$ (AF461158), *P.* aquimaris KF89$^T$ (KP716798), *P. indicus* IO390502$^T$ (MG845150) and *P. sediminilitoris* DSL-16$^T$ (MH491014); **Clade B** encompassed *P. aestuarii* B7$^T$ (EF660757), *P. hibisci* THG-T2.8$^T$ (KX456189), *P. pueri* THG-N2.35$^T$ (KX456187), *P. tibetensis* Tibet-S9a3$^T$ (DQ108402), *P. beibuensis* JLT1284$^T$ (EU650196) and *P. rhizosphaerae* CCM 7904$^T$

(MT760188); **Clade C** encompassed *P. angustae* E6$^{T}$ (KR052005), *P. fontiphilus* MVW-1$^{T}$ (LT223122), *P. sediminis* CMB17$^{T}$ (JX126474), *P. subflavus* GY0581$^{T}$ (MH880084), *P. haematequi* M1-83$^{T}$ (MH665405), *P*. acridae SCU-M53$^{T}$ (KT634253) and *P*. aerius 011410$^{T}$ (KX664462); **Clade D** encompassed *P. kocurii* JCM 7684$^{T}$ (D32241), *P. koreensis* Ch05$^{T}$ (AB187584), *P. jeotgali* CBA4604$^{T}$ (CP025583), *P. alkenifer* A901/1$^{T}$ (Y13827), *P. solventivorans* DSM 6637$^{T}$ (Y07705), *P. alcaliphilus* JCM 7364$^{T}$ (D32238), P. siganidrum M26$^{T}$ (JX398976), *P. alkanivorans* 4-2$^{T}$ (MH050839), *P. saliphilus* YIM 90738$^{T}$ (DQ923133), *P. oceanense* JLT1679$^{T}$ (HQ638977) and *P. stylophorae* KTW-16$^{T}$ (GQ281379); **Clade E** encompassed *P. chinensis* KS-11$^{T}$ (EU660389), *P. niistensis* NII-0918$^{T}$ (FJ842690), *P. salipaludis* WN007$^{T}$ (MF782381), *P. contaminans* WPAn02$^{T}$ (KX427102), *P. aeridis* JC501$^{T}$ (LT799401), *P*. marinus NBRC 100637$^{T}$ (AB681209), *P. endophyticus* SYSUP0003$^{T}$ (MH504123), *P*. luteus CFH 10530$^{T}$ (MK424273) and *P*. suum SC2-6$^{T}$ (MK027071); **Clade F** encompassed *P. aminovorans* JCM 7685$^{T}$ (D32240), *P. huijuniae* FLN-7$^{T}$ (EU725799), *P. mangrove* gyp-1$^{T}$ (LN879490), *P*. simplex F5$^{T}$ (MG938051), *P. laeviglucosivorans* 43P$^{T}$ (AB727354) and *P. limosus* NB88$^{T}$ (HQ336256); **Clade G** encompassed *P. cavernae* CECT 8482$^{T}$ (MT760298), *P. aurantiacus* TK008$^{T}$ (MN708965), *P. xiamenensis* 12-3$^{T}$ (MT137387), *P. pacificus* F14$^{T}$ (KF924610), *P. isoporae* SW-3$^{T}$ (FJ593906), *P. tegillarcae* BM15$^{T}$ (KJ789958), *P. lutimaris* HDM-25$^{T}$ (KJ451483) and *P. zhejiangensis* J6$^{T}$ (JN561152).

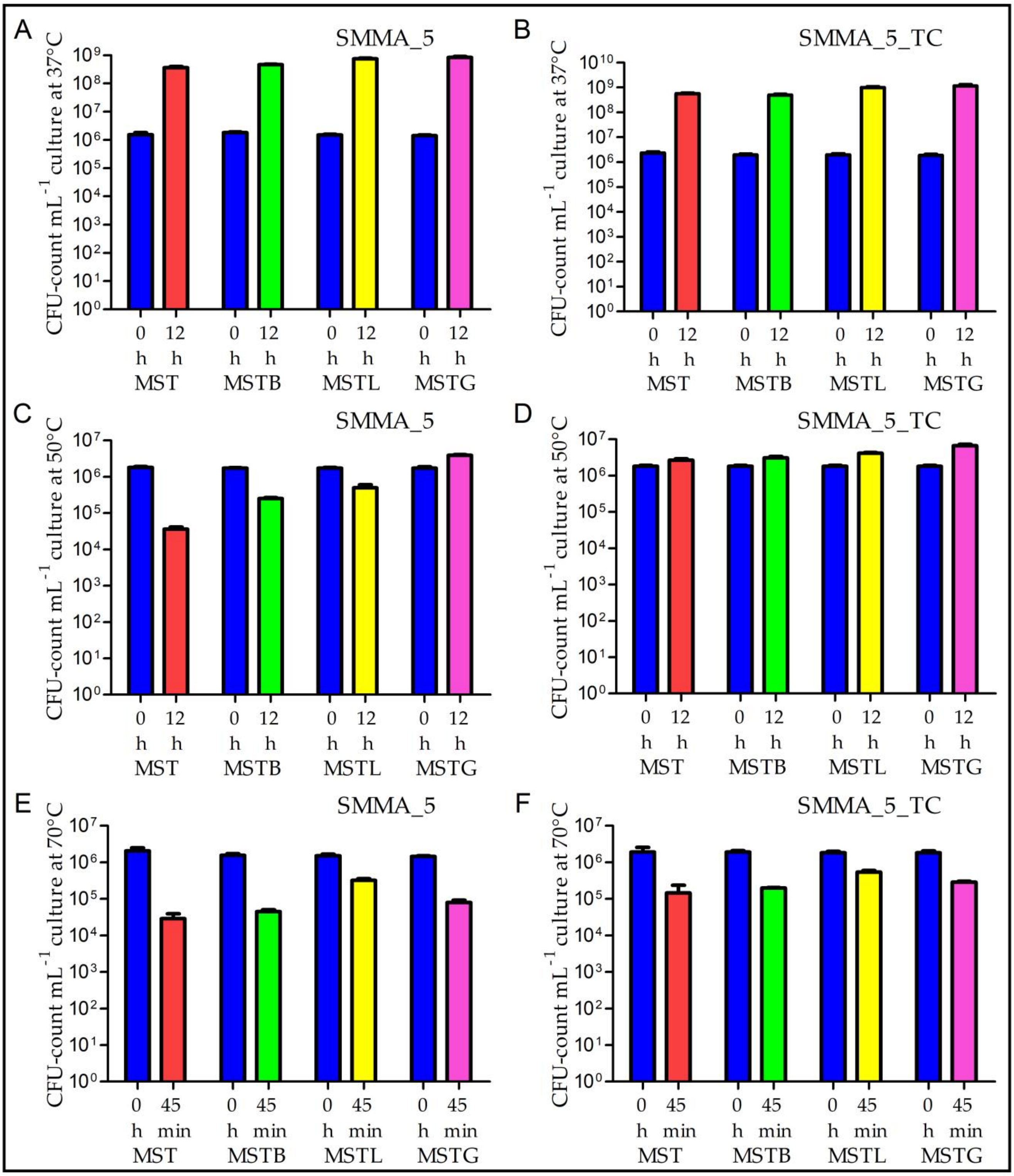


**Figure S2.** Increase or decrease in the CFU-counts of SMMA_5 and SMMA_5_TC, during incubation in MST, MSTB, MSTL and MSTG, at 37°C, 50°C and 70°C. (**A and B**) 0 h and 12 h CFU-counts recorded for SMMA_5 and SMMA_5_TC in the different MST variants at 37°C; (**C and D**) 0 h and 12 h CFU-counts recorded for SMMA_5 and SMMA_5_TC in the different MST variants at 50°C; (**E and F**) 0 h and 45 minute CFU-counts recorded for SMMA_5 and SMMA_5_TC in the different MST variants at 70°C. All the data shown in this figure are averages obtained from three different experiments; error bars indicate the standard deviations of the data. Across the panels, and irrespective of the bacterium considered, all 0 h data are represented by blue bars, while the data for incubations in MST, MSTB, MSTL and MSTG are represented by red, green, yellow and purple bars respectively.

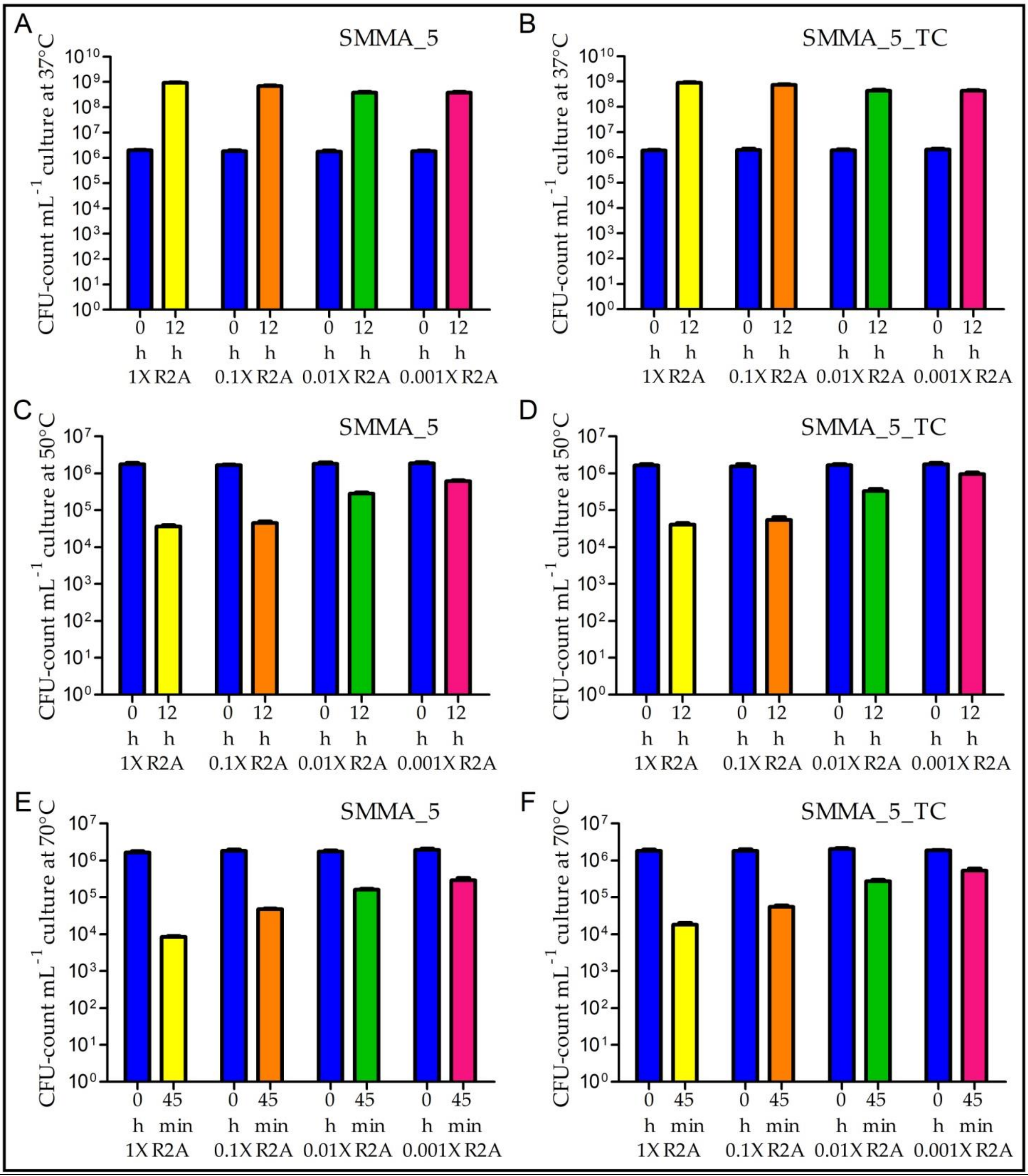


**Figure S3.** Increase or decrease in the CFU-counts of SMMA_5 and SMMA_5_TC, during incubation in the different dilution grades of R2A, at 37°C, 50°C and 70°C. (**A and B**) 0 h and 12 h CFU-counts recorded for SMMA_5 and SMMA_5_TC in the different dilution grades of R2A at 37°C; (**C and D**) 0 h and 12 h CFU-counts recorded for SMMA_5 and SMMA_5_TC in the different dilution grades of R2A at 50°C; (**E and F**) 0 h and 45 minute CFU-counts recorded for SMMA_5 and SMMA_5_TC in the different dilution grades of R2A at 70°C. All the data shown in this figure are averages obtained from three different experiments; error bars indicate the standard deviations of the data. Across the panels, and irrespective of the bacterium considered, all 0 h data are represented by blue bars, while the data for incubations in 1X R2A, 0.1X R2A, 0.01X R2A and 0.001X R2A are represented by yellow, orange, green and pink bars respectively.

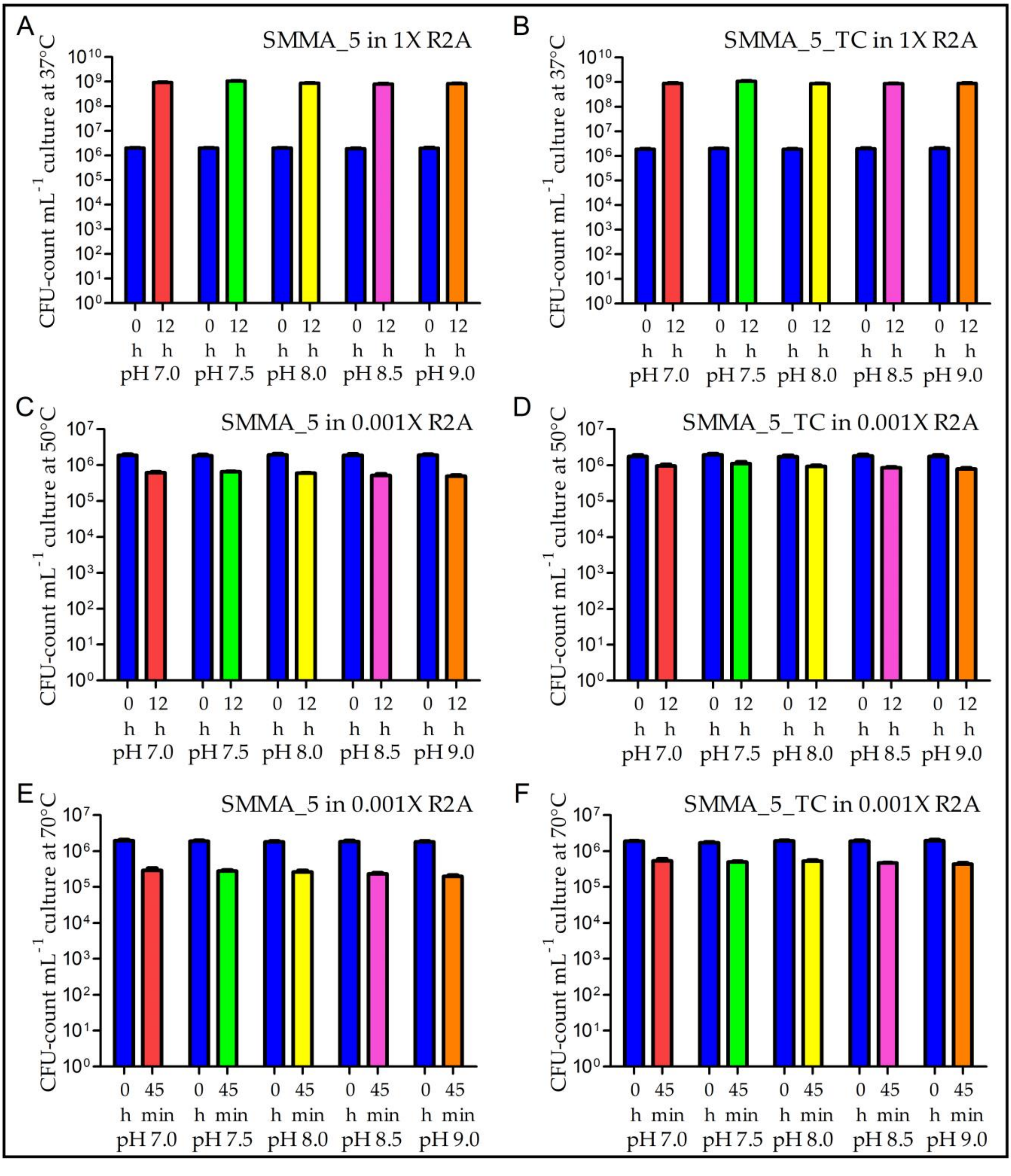


**Figure S4.** Increase or decrease in the CFU-counts of SMMA_5 and SMMA_5_TC, during incubation at 37°C, 50°C and 70°C in 1X R2A or 0.001X R2A medium having different pH levels. (**A and B**) 0 h and 12 h CFU-counts recorded for SMMA_5 and SMMA_5_TC at 37°C in 1X R2A having different pH levels; (**C and D**) 0 h and 12 h CFU-counts recorded for SMMA_5 and SMMA_5_TC at 50°C in 0.001X R2A having different pH levels; (**E and F**) 0 h and 45 minute CFU-counts recorded for SMMA_5 and SMMA_5_TC at 70°C in 0.001X R2A having different pH levels. All the data shown in this figure are averages obtained from three different experiments; error bars indicate the standard deviations of the data. Across the panels, and irrespective of the bacterium considered, all 0 h data are represented by blue bars, while the data for the incubations in R2A variants having pH 7.0, 7.5, 8.0, 8.5 and 9.0 are represented by red, green, yellow, purple and orange bars respectively.

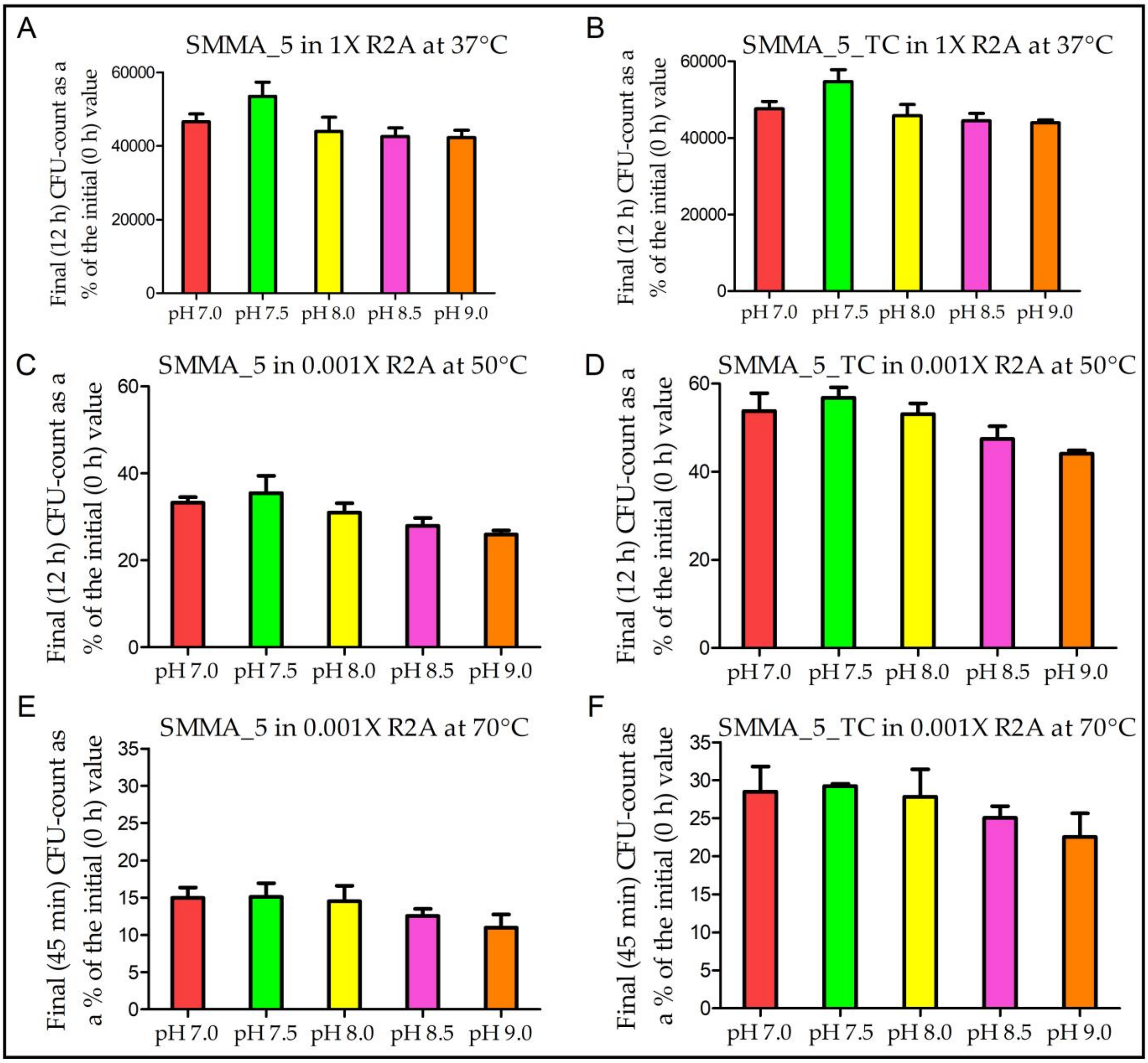


**Figure S5.** Final CFU-counts of SMMA_5 and SMMA_5_TC as percentages of the initial levels, after incubation at 37°C, 50°C and 70°C, in 1X R2A or 0.001X R2A medium having different pH levels. (**A and B**) final (12 h) CFU-counts of SMMA_5 and SMMA_5_TC at 37°C in 1X R2A having different pH levels, represented as percentages of the corresponding initial (0 h) CFU-counts; (**C and D**) final (12 h) CFU-counts of SMMA_5 and SMMA_5_TC at 50°C in 0.001X R2A having different pH levels, represented as percentages of the corresponding initial (0 h) CFU-counts; (**E and F**) final (45 minute) CFU-counts of SMMA_5 and SMMA_5_TC at 70°C in 0.001X R2A having different pH levels, represented as percentages of the corresponding initial (0 h) CFU-counts. All the data shown in this figure are averages obtained from three different experiments; error bars indicate the standard deviations of the data. Across the panels, and irrespective of the bacterium considered, all 0 h data are represented by blue bars, while the data for the incubations in R2A variants having pH 7.0, 7.5, 8.0, 8.5 and 9.0 are represented by red, green, yellow, purple and orange bars respectively.

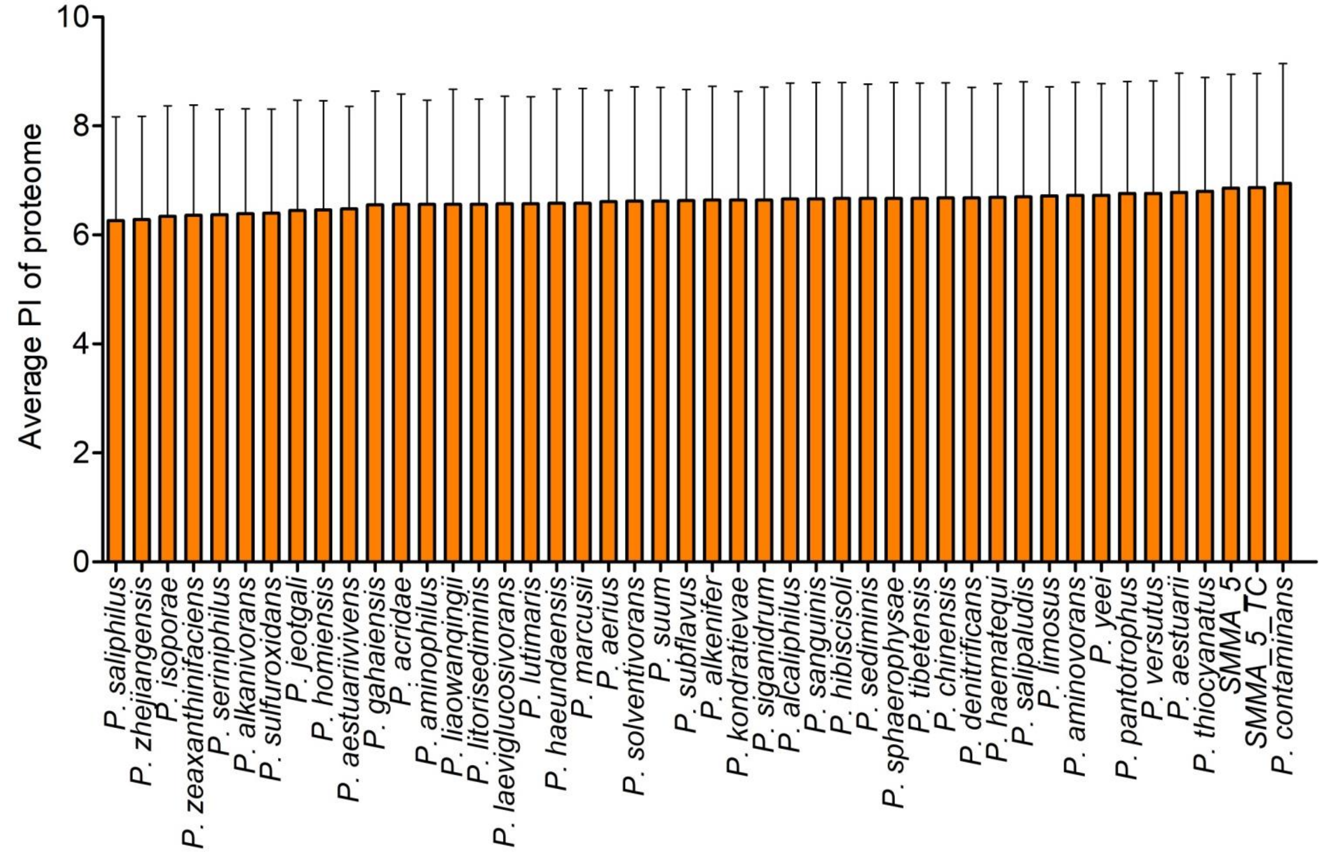


**Figure S6.** Average isoelectric points (pI values) predicted bioinformatically, and individually, for the putative protein contents of SMMA_5, SMMA_5_TC, and the 44 other *Paracoccus* species considered for comparative analysis (see Table S1 for the details of the comparator genomes).
